# A single-cell atlas characterizes dysregulation of the bone marrow immune microenvironment associated with outcomes in multiple myeloma

**DOI:** 10.1101/2024.05.15.593193

**Authors:** William C. Pilcher, Lijun Yao, Edgar Gonzalez-Kozlova, Yered Pita-Juarez, Dimitra Karagkouni, Chaitanya R. Acharya, Marina E Michaud, Mark Hamilton, Shivani Nanda, Yizhe Song, Kazuhito Sato, Julia T. Wang, Sarthak Satpathy, Yuling Ma, Jessica Schulman, Darwin D’Souza, Reyka G. Jayasinghe, Giulia Cheloni, Mojtaba Bakhtiari, Nick Pabustan, Kai Nie, Jennifer A. Foltz, Isabella Saldarriaga, Rania Alaaeldin, Eva Lepisto, Rachel Chen, Mark A. Fiala, Beena E Thomas, April Cook, Junia Vieira Dos Santos, I-ling Chiang, Igor Figueiredo, Julie Fortier, Michael Slade, Stephen T. Oh, Michael P. Rettig, Emilie Anderson, Ying Li, Surendra Dasari, Michael A Strausbauch, Vernadette A Simon, Immune Atlas Consortium, Adeeb H Rahman, Zhihong Chen, Alessandro Lagana, John F. DiPersio, Jacalyn Rosenblatt, Seunghee Kim-Schulze, Madhav V Dhodapkar, Sagar Lonial, Shaji Kumar, Swati S Bhasin, Taxiarchis Kourelis, Ravi Vij, David Avigan, Hearn J Cho, George Mulligan, Li Ding, Sacha Gnjatic, Ioannis S Vlachos, Manoj Bhasin

## Abstract

Multiple Myeloma (MM) remains incurable despite advances in treatment options. Although tumor subtypes and specific DNA abnormalities are linked to worse prognosis, the impact of immune dysfunction on disease emergence and/or treatment sensitivity remains unclear. We established a harmonized consortium to generate an Immune Atlas of MM aimed at informing disease etiology, risk stratification, and potential therapeutic strategies. We generated a transcriptome profile of 1,149,344 single cells from the bone marrow of 263 newly diagnosed patients enrolled in the CoMMpass study and characterized immune and hematopoietic cell populations. Associating cell abundances and gene expression with disease progression revealed the presence of a proinflammatory immune senescence-associated secretory phenotype in rapidly progressing patients. Furthermore, signaling analyses suggested active intercellular communication involving APRIL-BCMA, potentially promoting tumor growth and survival. Finally, we demonstrate that integrating immune cell levels with genetic information can significantly improve patient stratification.

## INTRODUCTION

Multiple myeloma (MM) is the second most prevalent hematological cancer, and its incidence continues to rise globally^1,2^. An estimated 35,780 new diagnoses and 12,540 deaths are projected for 2024 in the United States^3^. The emergence of myeloma-targeting biologic and immune-based therapies has led to significant improvements in patient outcomes^4^. Nevertheless, curative outcomes are characteristically elusive, and most MM patients eventually succumb to the disease. Disease evolution is associated with progressive immune dysregulation. With the recent FDA approval of immunotherapies, such as CAR-T cells and bispecific T cell engagers, understanding the immune elements in the myeloma microenvironment has become increasingly important for addressing disease emergence and/or response to treatment. Over the past 15 years, multiple studies^5–10^, including the Clinical Outcomes in Multiple Myeloma to Personal Assessment of Genetic Profiles (CoMMpass) registry^8,11^, have investigated the genomic landscape and diversity of MM as well as identified specific tumor subtypes and their underlying associations with clinical outcomes. Further, these studies have demonstrated that, like other cancers, MM tumors are multi-clonal, with their clonal makeup evolving over the course of the disease progression and exposure to treatments. Notably, prognostic models leveraging these genetic determinants are limited in their capacity to identify patients at high risk for early relapse. This suggests that latent, tumor-extrinsic factors contributing to patient prognosis are not captured by current models.

The bone marrow microenvironment (BMME) composition in MM has been identified as a factor affecting tumor progression and therapeutic outcomes. Recent studies have pointed to T cell exhaustion^12,13^ and the infiltration of immunomodulatory cell populations contributing to immunoediting and immune evasion in MM, such as myeloid-derived suppressor cells (MDSCs), regulatory T cells (T_reg_), Th17 cells, dendritic cells (DCs), and dysregulated natural killer (NK) cells, as well as tumor-associated neutrophils (TANs) and macrophages (TAMs)^14–17^. We hypothesized that profiling the BMME of newly diagnosed MM (NDMM) patients prior to treatment with standard myeloma therapies could reveal immune populations and signaling pathways associated with disease progression or clinical outcomes. Such insights can be used to refine current patient stratification tools including the revised International Staging System (R-ISS) and, importantly, inform strategies for target identification and rational integration of various immunotherapies in MM.

To this end, we generated a BMME Immune Atlas of NDMM patients from the Multiple Myeloma Research Foundation (MMRF) CoMMpass study (NCT01454297), which included corresponding detailed clinical and genomic information. Analyzing over 1.1 million cells from 263 NDMM patients, we identified immune populations and phenotypes associated with relapse risk and PFS.

## RESULTS

### A multiple myeloma bone marrow microenvironment cell atlas

To decipher the role of the BMME in MM outcomes, we profiled CD138^neg^ cells from 361 bone marrow (BM) aspirate samples (prior to treatment) obtained from 263 NDMM patients recruited in the CoMMpass study (**Figure 1a**). Patients who are either high cytogenetics risk or received doublet/triplet therapy, with or without autologous stem cell transplantation (ASCT), were preferentially selected from the CoMMpass study (n=1,143 patients) to be included in the study. This randomly selected sub-cohort was generally reflective of the CoMMpass study with similar (**Supplemental Table 1**) demographic and clinical characteristics, including median age (62.9 v. 64.1), percentage self-identified as African American (16.6 v. 17.5) ISS Stage 3 (27.9 v. 26.3) and cytogenetic high-risk^18^ (51.6 v. 53.2) (**Figure 1b, Supplemental Table 1)**. Therapeutically, 184 patients initially received a combination of proteasome inhibitors (PIs), immunomodulatory drugs (IMiDs), and steroids, while 135 underwent ASCT as first-line therapy. Overall, the study profiled before-treatment BM samples from 263 NDMM patients along with a subset of post-treatment samples (n = 98) (**Figure 1c**) using our previously standardized single-cell RNA sequencing (scRNA-seq) protocol^12,19,20^ (**Figure 1a**).

**Figure 1.**
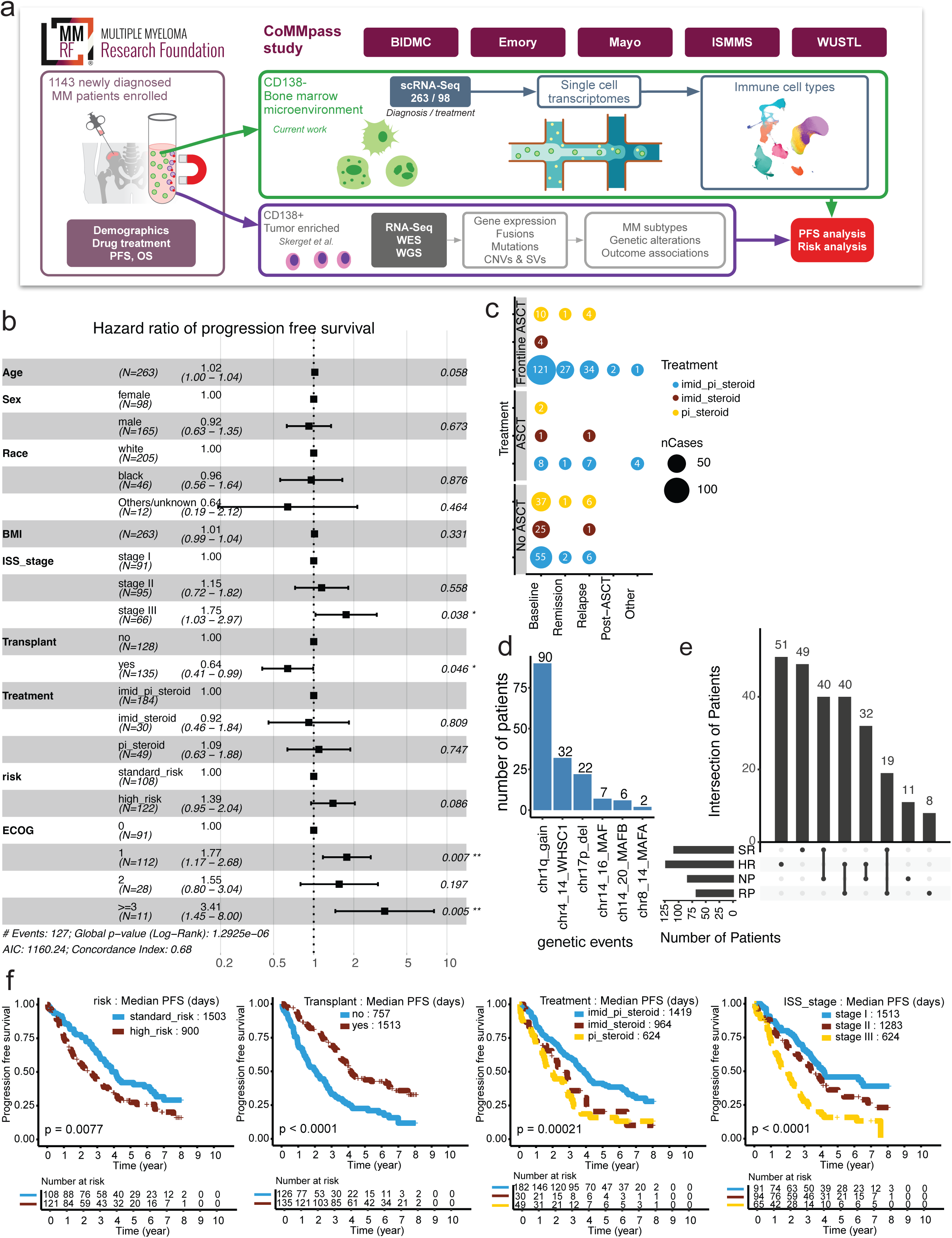
Overview of the immune atlas design, workflow, and patient characteristics. **(a)** Overview of the immune atlas study design, patient cohort, sample processing, and analysis workflow. **(b)** Clinical characteristics of patients (n = 263) in this study. The Forest plot illustrates the effect of various clinical features on progression-free survival (PFS). **(c)** Dot plot depicting the cross-section of samples based on autologous stem cell transplant (ASCT) and frontline treatment, where dot size indicates the number of patients and dot color indicates the type of treatment regimen. **(d)** Bar chart showing the total number of patients with each of the six genetic events used for risk classification. **(e)** Upset plot showing the intersection of patients categorized as standard-risk (SR) or high-risk (HR) and non-progressor (NP) or rapid progressor (RP) at baseline. **(f)** Kaplan-Meier curves display survival analysis for patients categorized based on risk stratification (HR vs SR), transplant as a frontline treatment, treatment type, and ISS staging.

By integrating single-cell transcriptomic profiles of CD138^neg^ and tumor-enriched genetic profiles of CD138^pos^ fraction of cells, we sought to examine how genetic alterations and the immune landscape correlate with outcomes (**Figure 1a**). Given previous findings from the CoMMpass study^11,18^, we used six genomics events to categorize patients into high-risk (HR, n = 123), if they met one or more of the following criteria: del17p13, t(14;16)[MAF], t(8;14)[MAFA], t(14;20)[MAFB], t(4;14)[WHSC1/MMSET/NSD2], gain of chromosome 1q; the remaining patients meeting none of these criteria were labeled as standard-risk (SR, n = 108) (**Figure 1d,e**). We additionally stratified patients based on their disease progression kinetics into rapid progressors (RP, n = 67), with progression events occurring within 18 months of diagnosis, and non-progressors (NP, n = 83), with durable remission for at least four years following treatment (**Figure 1e, Supplemental Figure 1**). Interestingly, although high-risk patients are mainly associated with rapid disease progression and vice versa, we identified 32 HR patients as NPs and 19 SR patients as RPs, indicating other factors, such as the immune environment, might play additional critical roles (**Figure 1e**). As expected, patients categorized as standard-risk had improved PFS relative to high-risk patients, suggesting our risk classification strategy was informative for predicting outcomes (**Figure 1f**). Additionally, survival analysis on other clinical variates also demonstrated that patients who either underwent BM transplants, received triplet treatment (PI, IMiD, and steroid), or were classified as ISS stage I presented significantly (*P* < 0.05) improved PFS (**Figure 1f**).

### Single-cell transcriptome profiling identifies traditional and rare cell types and subtypes of the myeloma BMME

Single-cell RNA-seq was performed on 361 samples collected at multiple time points from 263 patients, resulting in 1,149,344 high-quality BM cells (**Figure 2a**). On average, baseline samples consisted of T cells (30.51% CD8^+^, 23.39% CD4^+^), NK cells (6.82%), B cells (8.51%), Myeloid cells (12.20%), erythroblasts and erythrocytes (7.87%), and plasma cells (8.46%), with the remainder comprised of small, independent populations (HSCs, pDCs, Fibroblasts, 1.53%) (**Figure 2b,c**). Canonical lineage markers were used for cell type and subtype annotation with detailed information provided in the Supplemental Cell Population Annotation Dictionary (**Figure 2b, c, Supplemental Document 1**). For subsequent downstream analysis, we focused primarily on the baseline samples for all patients (n=263), unless otherwise specified.

**Figure 2.**
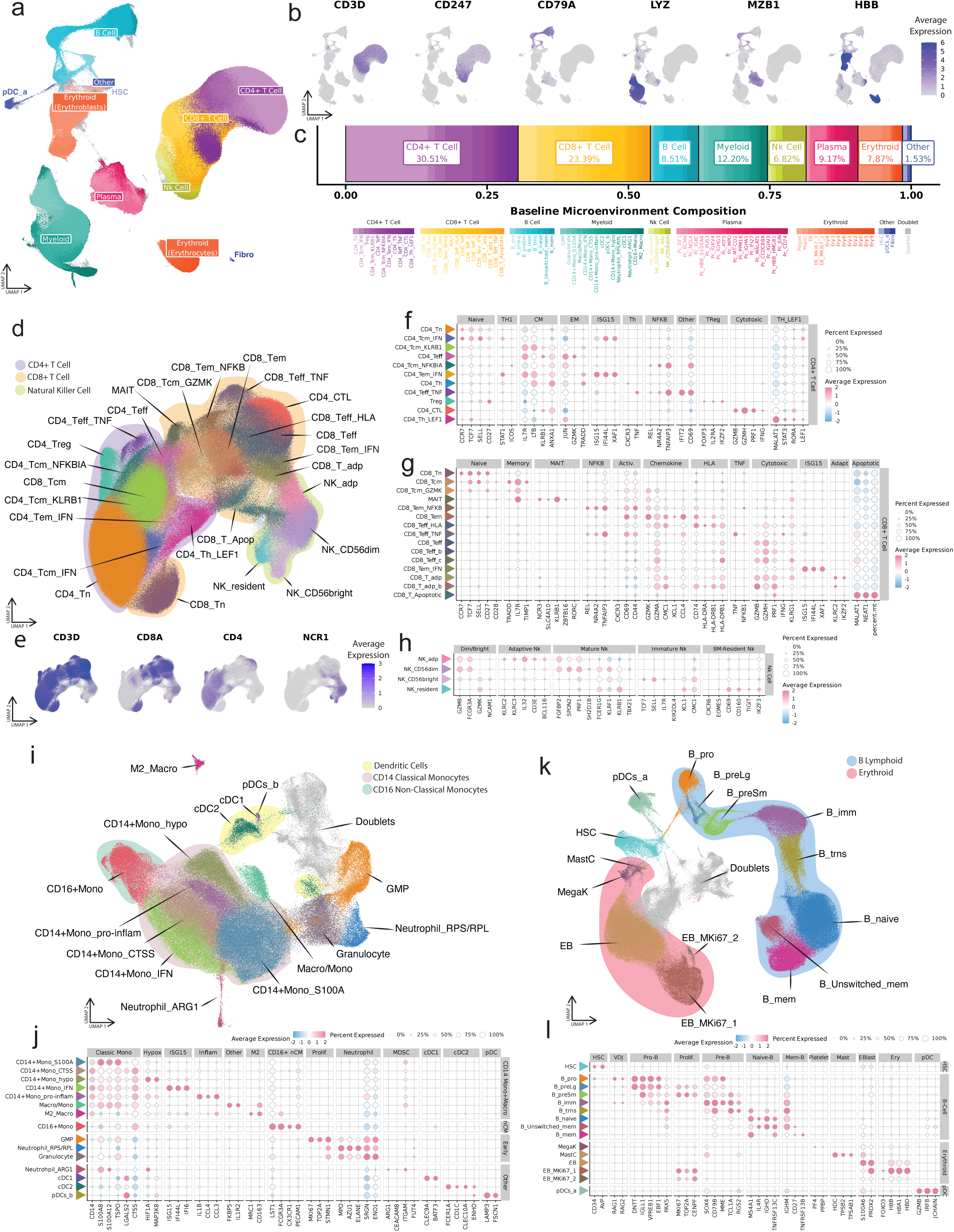
Single-cell immune atlas of multiple myeloma patient samples. **(a)** Uniform manifold approximation and projection (UMAP) embedding of 1,149,344 CD138^neg^ BMME cells collected from MM patients. A total of 106 clusters were observed, spanning five major compartments defined by canonical markers: T and NK cells, B cells and erythroblasts, myeloid cells, erythrocytes, and plasma cells. Populations identified as doublets are colored grey. **(b)** Feature plots displaying the normalized gene expression for a selection of lineage-specific markers. **(c)** A stacked bar chart, displaying the average per-patient cell type composition at baseline. Clusters are colored by their lineage and shaded by subtype. Doublet populations are omitted. **(d)** UMAP of the T Lymphocyte and Natural Killer compartment. Cells are colored by lineage *(CD4 T cells: purple; CD8 T cells: orange; NK cells: yellow)* and shaded by individual subtypes. The color legend is included in the corresponding dot plots (f-h). **(e)** Feature plots displaying the normalized gene expression per cell for markers to distinguish, CD4^+^, CD8^+^, and NK cells. **(f-h)** Dot plots displaying the average scaled expression of select marker genes used for precise cluster annotation. Expression is visualized on a red-blue color scale, with the size of each dot corresponding to the percent expression. Dot plots are split by lineage into **(f)** NK cells, **(g)** CD8^+^ T cells, and **(h)** CD4^+^ T cells. The colored triangle next to the cluster name matches the cluster color in the corresponding UMAP **(d)**. **(i)** UMAP of the myeloid compartment. Cells are shaded by their subtype. Doublet populations are colored grey. **(j)** Dot plot displaying the average scaled expression of select marker genes for precise cluster annotation in the myeloid compartment. Expression is visualized on a red-blue color scale, with the size of each dot corresponding to the percent expression. The triangle next to the cluster name matches the cluster color in the corresponding UMAP. **(k)** UMAP of the B cell and erythroblast compartment. Cells are colored by their lineage *(B cells: cyan; Erythrocytes: red; Other: dark blue)*, shaded by subtype. Doublet populations are colored grey **(l)** Dot plot displaying the average scaled expression of select marker genes used for precise cluster annotation in the B cell and erythroblast compartment. Expression is visualized on a red-blue color scale, with the size of each dot corresponding to the percent expression. See **Supplemental Document 1** for a detailed description of the annotation of all individual clusters. The colored triangle next to the cluster name matches the clusters in the corresponding UMAP **(k)**.

The T and NK cells compartment formed 30 clusters across CD4^+^ (11 clusters), CD8^+^ (15 clusters), and NK (4 clusters) cell populations (**Figure 2d, e**). CD4^+^ T cell clusters comprised naïve, central memory, effector memory, regulatory, and helper T cells (**Figure 2f, Supplemental Figure 2a**). This large-scale analysis also enabled the identification of rare cytotoxic CD4^+^ T cells with high expression of *GZMB* and *PRF1* markers. Similarly, the CD8^+^ T cell population also comprised multiple clusters of memory, and effector cells as well late activated effector subtypes (i.e., CD8_Teff_HLA) with intermediate-to-low expression of cytotoxic markers, but high expression of HLA-I and HLA-II class markers (**Figure 2g, Supplemental Figure 2b**). The NK cell clusters comprised classical CD56^+bright^, CD56^+dim^, as well as rare adaptive, and BM resident natural killer cell types (**Figure 2h**).

The myeloid lineage comprised 18 clusters of classical CD14^+^ and non-classical CD16^+^ monocytes, granulocytes, neutrophils, cDCs, pDCs, and macrophages (**Figure 2i, j**). The B cell compartment contained pro-B cells, immature transitional B cells, naïve, and memory B cells (**Figure 2k**). The compartment also captured immature hematopoietic populations, such as HSCs, mast cells, and erythroblasts. A small distinct population of mature erythrocytes was observed (**Figure 2a, Supplemental Figure 2c**) with nine subclusters exhibiting minimal patient-specific heterogeneity.

Additionally, we identified a population of plasma cells representing 9.17% of cells on average in baseline samples (**Supplemental Figure 2d**), likely comprising residual myeloma populations in the CD138^neg^ BM fraction, indicated by driver variants and copy number changes. Plasma cell subpopulations showed some associations with disease progression, with potential implications for patient outcomes (**Supplemental Figure 3a-c**).

### High-risk MM patients exhibit an over-representation of dysfunctional CD8^+^ T effector populations and interferon-stimulated myeloid cell diminution

Previous studies identified the associations of somatic and germline alterations as well as copy number variations with cytogenetics high-risk disease and poor outcomes in MM^11,18^. The high-risk abnormalities in this sub-cohort were similar to CoMMpass (**Supplemental Table 1**) and other NDMM studies^11,18^, with 39.0% of patients with 1q gain, 13.9% with t(4;14)[WHSC1/MMSET/NSD2], and 24.4% of patients having two or more lesions (**Figure 3a, Supplemental Table 1, 2**). Based on the Skerget *et al.*^11^ risk stratification of CoMMpass patients (**Figure 1a,d**), we performed a comparison of the BMME cells between high and standard-risk patients at baseline and observed significant differences in the CD8^+^ T and myeloid cell compartments.

**Figure 3.**
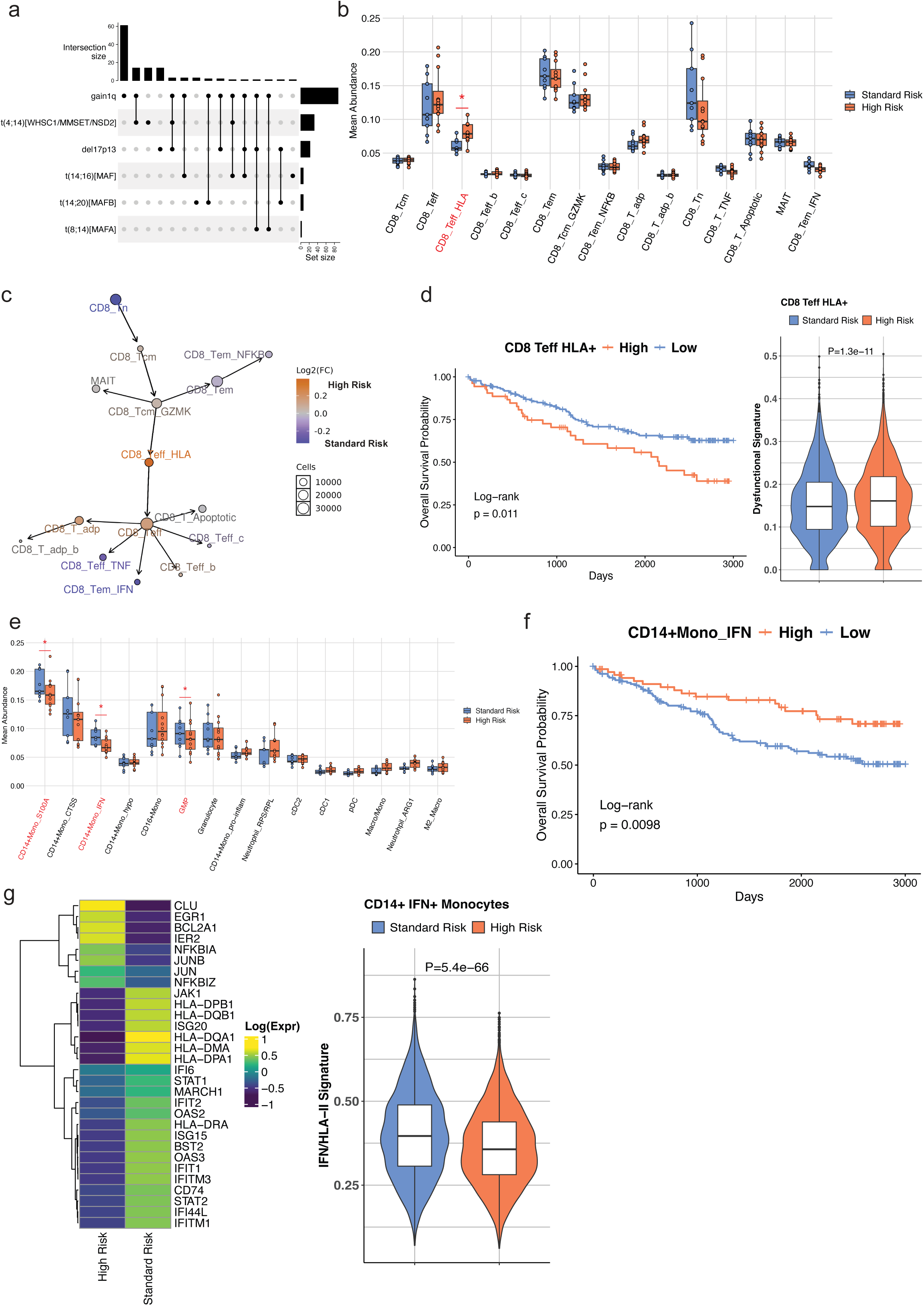
Alterations in the MM bone barrow microenvironment based on cytogenetic risk profile. **(a)** UpSet plot of the genetic lesions comprising the definition of the high-risk group. **(b)** Boxplots of the CD8+ T cell mean abundance estimates across batches for the high (n = 123) and standard (n = 108) risk groups at baseline from the Dirichlet regression model adjusting for batch. Significantly different proportions between high- and standard-risk patients are denoted with a red asterisk (* p-value < 0.05). **(c)** Pseudotime trajectory of the CD8 T cells, coupled with the mean cell composition fold change differences (log2 scale) between high- and standard-risk patients at baseline, presented in a blue-to-orange color scale. The size of the nodes corresponds to the number of cells per cluster. **(d)** *(Left)* Survival curve from regressing the overall survival on the CD8 Teff HLA+ cell abundances. The cell abundances were corrected for batch by taking the Pearson residuals from a Dirichlet regression model with batch as the covariate. The cut-off was determined using maximally selected rank statistics and set at the 77% quantile (177 low, 53 high). *(Right)* Violin plots of the Ucell signature enrichment of CD8 Teff HLA+ cells based on the expression of dysfunctional genes (*CD57, ZEB2, KLRG1, KLRK1, TIGIT, LAG3, PDCD1, CTLA4, TIM-3)* between the high- and standard-risk patients at baseline. The p-value denoting the significantly different enrichment is displayed (Wilcoxon rank sum test, two-sided). **(e)** Boxplots of the monocyte cell mean abundance estimates across batches for the high (n = 123) and standard (n = 108) risk groups at baseline from the Dirichlet regression model adjusting for batch. Significantly different proportions between high- and standard-risk patients are denoted with a red asterisk (* p-value < 0.05). **(f)** Survival curve from regressing the overall survival on the CD14+ IFN+ monocytes cell abundances. The cell abundances were corrected for batch by taking the Pearson residuals from a Dirichlet regression model with batch as the covariate. The cut-off was determined using maximally selected rank statistics and set at the 69% quantile (159 low, 71 high). **(g)** *(Left)* Heatmap displaying the average expression of HLA-II and IFN-related significantly differentially expressed genes in the CD14+ IFN+ monocytes between the high and standard-risk groups at baseline. *(Right)* Violin plots of the Ucell signature enrichment of CD14+ IFN+ monocytes based on the expression of the 23 IFN and HLA-II induced downregulated genes presented in the heatmap between the high- and standard-risk patients at baseline. The p-value denoting the significantly different enrichment is displayed (Wilcoxon rank sum test, two-sided).

Trajectory analysis of the CD8^+^ T cell compartment captured the differentiation trajectory spanning from early-stage naïve T cells to memory and to cytotoxic effector T cells, providing a framework for exploring dynamic changes in gene expression and cellular proportions (**Figure 3b,c**). Specifically, high-risk patients presented significantly elevated proportions (*P* = 0.013, log2 fold-change = 0.462, **Figure 3c**) of dysfunctional CD8^+^ T cell (i.e., CD8_Teff_HLA) and lower abundances of differentiated, highly activated, and pro-inflammatory cells, also indicated by their position in pseudotime lineage trajectories (**Figure 3b,c**). The higher abundances of these putatively dysfunctional CD8^+^HLA^+^ T effector cells (CD8_Teff_HLA) were significantly associated with poor overall survival (OS) (log-rank test *P* = 0.011, **Figure 3d**) and PFS (log-rank test *P* = 0.032, **Supplemental Figure 4a**). The CD8^+^HLA^+^ T effector cells in the high-risk cohort was further associated with a significantly increased CD8 T cell dysfunctional signature score, as captured by the expression of nine key marker genes (*CD57, ZEB2, KLRG1, KLRK1, TIGIT, LAG3, PDCD1, CTLA4, TIM-3*)^21–24^ (*P* = 1.27×10^-11^, **Figure 3d, Supplemental Table 4)**.

Among myeloid cells, we observed three distinct cell populations depleted in the high-risk group (CD14+Mono_S100A, CD14+Mono_IFN, GMP) (**Figure 3e**). In particular, the interferon-stimulated CD14^+^ monocytes with high expression of interferon and HLA-II class genes (CD14+Mono_IFN) exhibited lower abundance in the high-risk group (*P* = 0.05, log_2_FC = -0.224, **Figure 3e**), associated with better OS (log-rank test *P* = 0.0098, **Figure 3f**) and PFS (log-rank test *P* = 0.042, **Supplemental Figure 4b**). Differential gene expression analysis (DGEA) identified a down-regulation of 23 interferon type I and HLA class-II activity-related genes in CD14^+^IFN^+^ monocytes of the high-risk group (**Figure 3g**). Gene set enrichment analysis using this signature depicted a significantly lower score in high-risk patients (*P* = 5.4×10^-66^, **Figure 3g**), suggesting antigen presentation loss and induction of an immunosuppressive microenvironment.

After characterizing the myeloma BMME at baseline, we assessed whether the observed patterns also manifested across time in follow-up samples. In our cohort, 39 patients with baseline cytogenetic information (22 high-risk, 17 standard-risk) had follow-up samples at either relapse or remission, which also had cytogenetic information. From these 39 patients (87 samples), we selected 34 patients (74 samples), consisting of 19 high-risk (42 samples) and 15 standard-risk (32 samples), who progressed/relapsed (**Supplemental Figure 5a**). Interestingly, the proportion of regulatory T cells (T_regs_) increased significantly at relapse timepoints relative to baseline in the high-risk group, indicating expansion of an immunosuppressive phenotype (*P* = 0.026, **Supplemental Figure 5b**). A trend of increasing T_regs_ over time was also observed in relapsed standard-risk patients (R^2^ = 0.077, BH-FDR = 2.6e^-26^, **Supplemental Figure 5c**), with a less prominent abundance than in the high-risk patients (R^2^ =0.029, BH-FDR = 3.1e-15,). Among CD8^+^ T cells, the dysfunctional CD8_Teff_HLA increased over time in the high-risk group (R^2^ = 0.035, BH-FDR = 3.7e^-22^, **Supplemental Figure 5d**). On the other end, classical (CD14+Mono_S100A) and non-classical monocytes (CD16+Mono) were significantly depleted in the high-risk group over time (CD14+Mono_S100A: (R^2^ = 0.052, BH-FDR = 3.2e^-44^), CD16+Mono: (R^2^ =0.014, BH-FDR=7.1e^-08^, **Supplemental Figure 5f,g**) with the most pronounced decrease observed in the proinflammatory CD14^+^IFN^+^ monocytes relevant to our observations at baseline (R^2^ =0.155, BH-FDR =3.6e-60, **Supplemental Figure 5e**). This analysis provides preliminary insights after the treatment changes in BMME for comparison with baseline NDMM profiles that will be explored deeply in future studies.

### Rapid progressors display accumulation of effector and depletion of naïve T cell populations

We subsequently investigated alterations in the BMME based on disease progression, comparing patients who rapidly progressed within <18 months (RP, n = 67) following initial therapy to those who achieved sustained remission or non-progression for at least 4 years (NP, n = 83) (**Supplemental Figure 1a**). Most of these patients received standard triplet therapy, consisting of a PI, IMiD, and a glucocorticoid as their first line of therapy (**Supplemental Table 3**). Broadly, rapid progressors had lower abundances of CD4^+^ T cells and B cells, and higher levels of myeloid, plasma, and erythroid cells relative to non-progressors (**Figure 4a**). Samples from rapid progressor patients also exhibited significant enrichment of myeloid cells (P<0.05) and lower levels of B cell populations (P<0.05, **Figure 4b)**, including the immature, transitional, naive and memory B-cell clusters (P<0.05, **Supplemental Figure 6**). This suggests a shift towards myelopoiesis in patients with rapid progression of the disease, an indicator of stressed BMME^25^.

**Figure 4.**
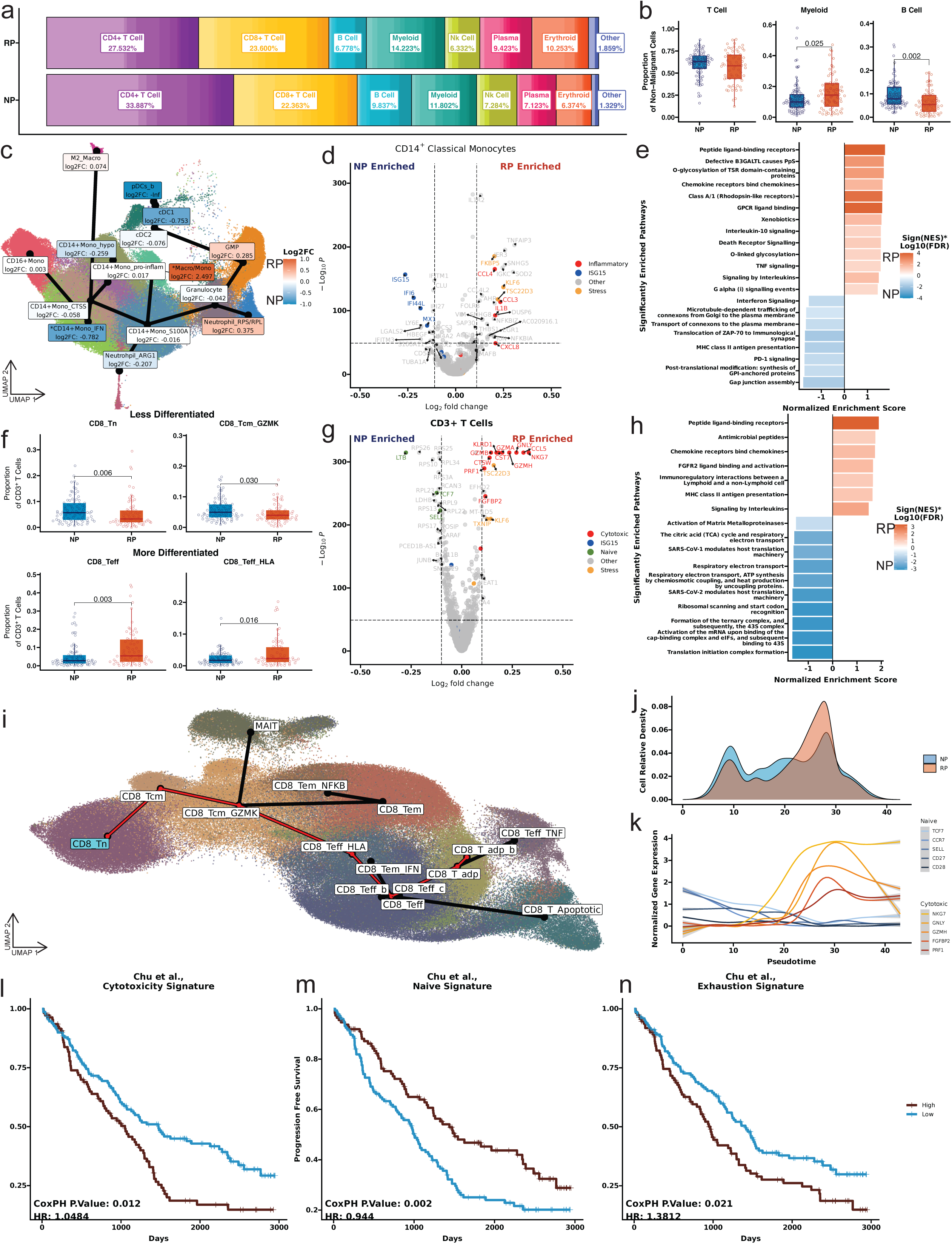
Single-cell level alterations in the bone marrow microenvironment of rapidly progressing MM patients. **(a)** Stacked bar chart displaying the mean per-patient cell type proportions at baseline across the full dataset, split between RP and NP patients. Clusters are colored by their major cell type and shaded by individual clusters. The average proportion of major cell types has been shown on the graph. **(b)** Box plots displaying the distribution of per-patient proportions for T cells, B cells, and myeloid cells as a percentage of non-malignant cells, split by progression categories. Doublet populations are excluded. Open circles represent individual patients. The significance of the difference in proportions between Rapid and Non-progressors has been calculated using the Wilcoxon rank sum test. The non-significant p-values >0.05 are not shown. **(c)** Trajectory of the CD14^+^ monocyte population along with differential abundance results. Lines represent transitions along the trajectories, with circles representing each cluster, and branches representing different lineages. Cluster labels along with fold change (log2) in cluster proportion between NP and RP samples have been shown along the trajectory. Labels are also shaded by fold change (log2) and clusters with significant differences in proportion were marked using an asterisk (* *P* < 0.05; ** *P* < 0.01; Wilcoxon rank sum test). **(d)** A volcano plot displaying the differentially expressed genes between NP and RP patients across the CD14+ monocyte compartment. The x-axis displays the log 2-fold change and the y-axis -log_10_ of the BH-adjusted p-value. The significantly differentially expressed genes associated with inflammation and ISG15 antiviral pathways are shown with red and blue colors respectively. **(e)** A bar plot displaying gene set enrichment analysis results for the differentially expressed genes shown in **(d)**. The x-axis displays the normalized enrichment score for each pathway, where positive values indicate the pathway associated with the NP-enriched markers, and negative values indicate the pathway associated with the RP-enriched markers. Pathways are shaded by -log_10_FDR and. The pathways with BH-adjusted p values < 0.1 were considered significant and shown in the plot. **(f)** Box plots displaying the distribution of per-patient proportions of selected significantly enriched CD3+ T Cell populations. Doublet populations are excluded. Open circles represent individual patients. The significance of the difference in proportions between Rapid and Non-progressors has been calculated using the Wilcoxon rank sum test. The non-significant p-values >0.05 are not shown. **(g)** A volcano plot displaying the differentially expressed genes between NP and RP patients across all CD3^+^ T cells. The x-axis displays the log 2-fold change and the y-axis -log_10_ of the BH-adjusted p-value. Select genes are highlighted and colored based on their associated function. **(h)** A bar plot displaying gene set enrichment analysis results for the differentially expressed genes shown in **(g)**. The x-axis displays the normalized enrichment score for each pathway, where positive and negative values indicate the pathways associated with the NP-enriched and RP markers respectively. The pathways are shaded by signed FDR. The pathways with BH-adjusted p values < 0.1 were considered significant and shown in the plot. **(I)** Trajectory analysis of CD8^+^ T cells. Cells are colored by clusters with trajectories connecting clusters drawn in black, with the trajectory strongly associated with cytotoxic populations highlighted in red. The cluster set as the origin for the resulting trajectory is highlighted in cyan color. **(j)** A density plot showing the distribution of cells concerning their pseudotime along the cytotoxicity lineage. Low pseudotime corresponds to cells closer to the origin cluster (CD8_Tn), while later pseudotime corresponds to differentiated cytotoxic clusters. **(k)** The smoothed normalized gene expression along the cytotoxicity lineage’s pseudotime for five Naïve associated genes (blue) and five Cytotoxicity associated genes (red). Gene expression for individual cells is weighted by slingshot’s lineage assignment weight. **(l-n)** Survival plot displaying the relationship between progression-free survival and the patient’s average cytotoxicity signature **(l)**, naïve signature **(m)**, or exhaustion signature **(n)** scores across all CD3^+^ T Cells. The red curve corresponds to patients with a signature score greater than the median, while the blue curve corresponds to patients with a signature score less than the median patient. The p-value for the CoxPH model fitted on the continuous signature score, correcting for processing site and batch, is displayed in the bottom left corner of each panel.

To delve deeper into this shift towards myeloid versus lymphoid populations in rapid progressors, we performed a trajectory analysis on the myeloid compartment revealing the expected progression from immature CD14^+^ to mature CD16^+^ monocytes transitioning along the path from non-inflammatory to activated CD14^+^ monocytes (**Figure 4c**). The rapid progressors presented significantly elevated proportions of CD14^+^CD163^+^ monocytes (Macro/Mono, p=0.004) and lower abundance of proinflammatory CD14^+^ monocytes (CD14+Mono_IFN, p=0.04) (**Figure 4c, Supplemental Figure 7**), similar to observations in high-risk patients (**Figure 3e**). DGEA across the CD14^+^ monocyte populations identified significant upregulation of proinflammatory markers in rapid progressors, such as *CCL3*, *CCL4*, *IL1B*, and *CXCL8*, whereas interferon signaling-related genes were increased in non-progressors (i.e., *ISG15*, *IFI6*, *IFI44*, *MX1*) (**Figure 4d**). Pathway analysis further highlighted the enrichment of proinflammatory pathways, including interleukin, TNF, IL-10, and chemokine signaling in the rapid progressors, pointing to a polarization towards an immunosuppressive phenotype. Conversely, significant enrichment of MHC-II antigen presentation and interferon signaling pathways was observed in non-progressors suggesting classical antigen processing and presentation (**Figure 4e**).

The focused analysis of the T cell compartment identified a significantly higher proportion (P=0.01) of CD8^+^ T cells (45.5%) in rapid progressors in comparison to non-progressors (39.9%) (**Supplemental Figure 8**). A significantly higher proportion of early-stage CD8^+^ naïve (T_n,_ P=0.006) and CD8^+^ GZMK^+^ central memory (CD8_Tcm_GZMK, P=0.03) cells were identified in non-progressors, while rapid-progressor patients exhibited higher abundance of differentiated CD8^+^ cytotoxic effector (CD8_Teff, P=0.003) and HLA^+^ effector (CD8_Teff_HLA, P=0.01) populations (**Figure 4f**, **Supplemental Figure 9**). DGEA further confirmed this by identifying a significant upregulation of cytotoxicity markers (*NKG7, GNLY, PRF1, FGFBP2, KLRD1, GZMB, GZMA,*) in the T cells of rapid progressors. The T cells of non-progressors demonstrated significant upregulation of markers genes for early-stage, naïve populations (*LTB, TCF7, SELL*) (**Figure 4g**). Furthermore, rapid progressors presented a significant enrichment of interleukin and chemokine signaling pathways, while non-progressors were primarily enriched in ribosomal and translational pathways, likely reflecting the high rRNA expression associated with more naïve T cell populations^26^ (**Figure 4h**). We assessed the differences in cellular abundance and gene expression across the lineage consisting of cytotoxic populations in our CD8^+^ trajectory **(Figure 4i).** The analysis of cellular proportions along the pseudotime of the cytotoxic cell lineage from trajectory analysis (**Figure 4j**) revealed a higher density of cells from rapid-progressors at later pseudotime points, corresponding with the CD8 T effector populations (i.e., CD8_Teff_HLA, CD8_Teff). In contrast, cells from non-progressor patients exhibited relatively higher densities at earlier pseudotime points, corresponding to naive and memory T cells (i.e., CD8_Tn, CD8_Tcm_GZMK). Further evaluation of gene expression along the trajectory depicted that markers related to cytotoxicity (*NKG7*, *GZMH*, *FGFBP2*) achieved the highest expression toward the end region of the trajectory (later pseudotime) and were majorly enriched in cells from rapidly progressing patients. This end region of the trajectory presented minimal or near zero expression of markers corresponding to early-stage naïve T cell populations (*CCR7*, *SELL*, *TCF7*, *CD27*, *CD28)* that were enriched in non-progressors (**Figure 4k**). This suggests accumulation of terminally differentiated, cytotoxic, CD27^neg^CD28^neg^ CD8+ T effector cells in rapid-progressing patients, accompanied by a corresponding reduction in the healthy naïve and central memory pool necessary for mounting an immunological memory, which might be associated with poor outcomes. To further explore this hypothesis, we evaluated associations of independent cytotoxic and naïve CD8^+^ T cell gene signatures from the pan-cancer T cell atlas^27^ to predict outcomes in our dataset. The higher abundance of signature positive cytotoxic CD8^+^ T cells was associated with worse PFS (CoxPH *P*<0.012) (**Figure 4l, Supplemental Table 4**). Conversely, patients enriched in a naïve-like signature across their T cell compartment displayed better PFS (CoxPH *P* < 0.002) (**Figure 4m, Supplemental Table 4**). Additionally, patients with a higher CD3^+^ T cell exhaustion signature scores also presented poor PFS (CoxPH *P* < 0.021) (**Figure 4n, Supplemental Table 4**). However, expression of exhaustion did not correspond to the RP-enriched CD8_Teff population, and seems to primarily originate from CD8_Tem, CD8_Teff_HLA, CD8_Tem_IFN, and CD8_T_adp populations (**Supplemental Figure 10)**.

We also repeated the above T cell compartment analysis using only samples from patients treated with triplet therapy. While therapy itself cannot impact the baseline immune composition, it can influence outcomes. These analyses also showed similar results with a significant enrichment of naïve CD8 T cells and B cells in the non-progressors, as well of the enrichment of more differentiated T cell populations, such as CD8_Teff and CD8_Teff_HLA, in rapid-progressors on triplet therapy (**Supplemental Figure 11**).

### Enrichment of CD4^+^ cytotoxic T cells in the immune microenvironment distinguishes rapid progressors with standard-risk cytogenetics

Considering the cellular alterations observed across cytogenetic risk- and progression-based groups, we next sought to determine whether there are common changes in the immune microenvironment associated with high-risk and rapid disease progression. Additionally, we sought to gain insights into why patients assigned to the standard-risk group may progress rapidly despite their classification. Both high-risk and rapid progressors exhibited significant enrichment of the CD8_Teff_HLA population and depletion of IFN-α stimulated CD4^+^ and CD8^+^ naïve, memory, and effector populations (i.e., CD8_Tem_IFN, CD4_Tem_IFN, CD4_Tcm_IFN, CD14_Mono_IFN), as well as classical monocytes and immature and naïve B cells (**Figure 5a**). Notably, standard-risk and non-progression-associated T cell populations displayed common enrichment of IFN-stimulation (ISG15) related genes, including *ISG15, MX1, OAS1, IFI6,* and *IFI44L*.

**Figure 5.**
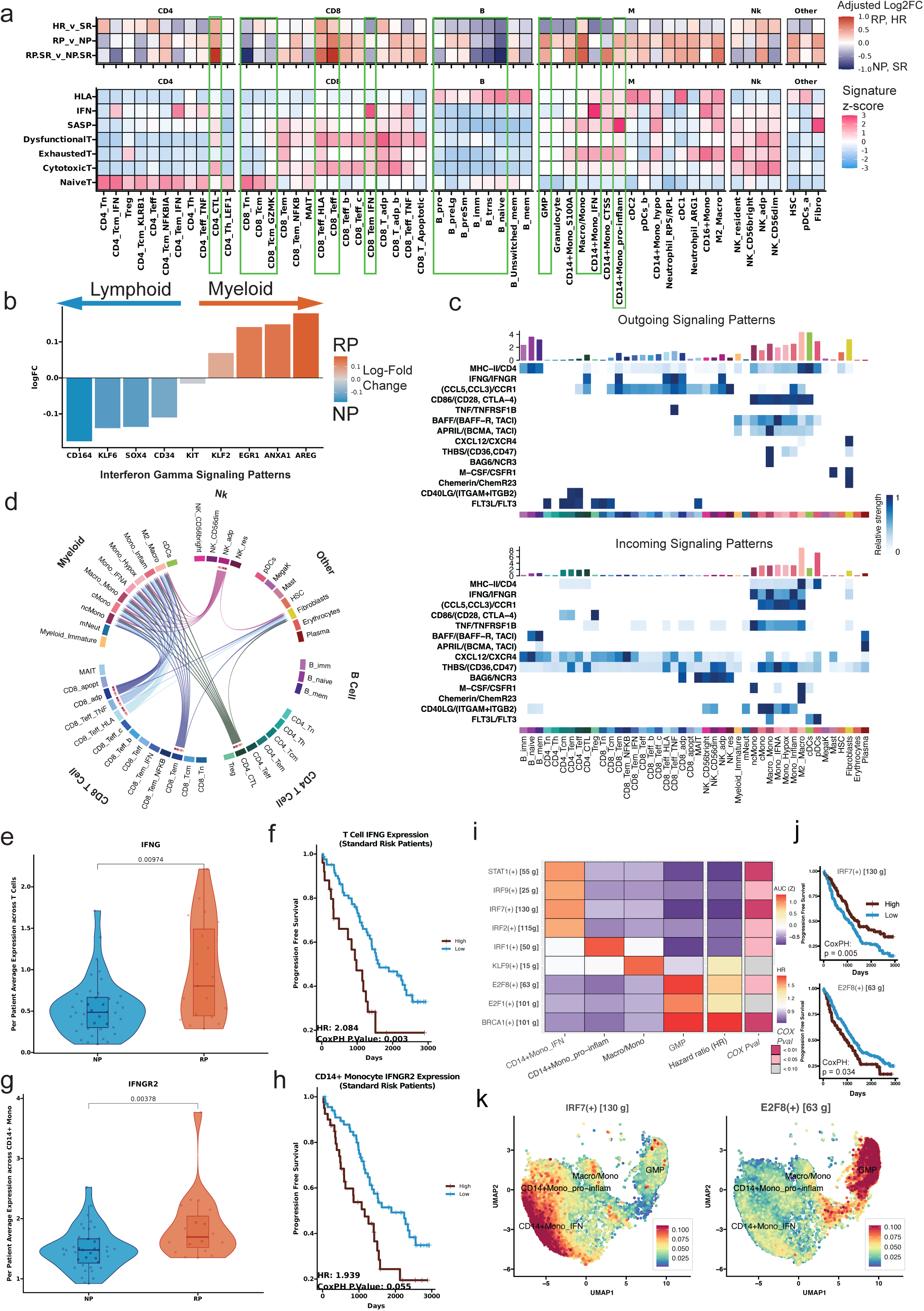
Pathway and systems biology analysis to decipher mechanisms of poor outcomes in MM. **(a)** *(Top)* Comparison of differential abundance results by cytogenetic risk (HR_v_SR), progression (RP_v_NP), and progression within standard-risk patients (RP.SR_v_NP.SR). Each box represents the estimated log2-fold change between different risk and progression based comparisons (HR v SR, RP v NP, RP.SR v NP.SR) after adjusting for batch using a Dirichlet model. Rows represent different comparisons; columns represent different cell populations. Orange shades indicate RP, HR, or RP.SR upregulated cell populations, while blue shades indicate SR, NP, or NP.SR upregulated populations. *(Bottom)* Average normalized signature scores for select immune signatures (Supplemental Table 4) across the various cell populations. **(b)** Bar graph displaying differentially enriched markers between Non and Rapid progressors within the CD34^+^ HSC population. Bars represent the log2-fold change, with positive indicating the gene is enriched in non-progressors, while negative blue bars indicate enrichment in Rapid progressors. **(c)** Heatmap of intercellular communication depicting key patterns of outgoing *(top)* and incoming *(bottom)* signaling communication between cell types. Each row of the heatmap represents a ligand-receptor pair, where the relative strength of the outgoing signal (ligand expression) by each cell type is shown on the top heatmap, and the relative strength of the corresponding incoming signal (receptor expression) by each cell type is shown on the bottom heatmap. Key signaling pairs between T cell and myeloid populations are depicted, including cytokine and IFN-g signaling, in addition to BAFF and APRIL signaling patterns implicated in MM. **(d)** Chord diagram indicating the IFN-g signaling network in our dataset. Chords are colored by the ‘sender’ cell type (population expressing the ligand) and point towards the ‘receiver’ cell type (population expressing the receptor), illustrating strong outgoing IFN-g signaling from T and NK cell populations, corresponding to increased incoming IFN-g signaling to myeloid and fibroblast populations. **(e-h)** Comparing the average expression of IFN-g across T cells **(e-f)** and IFN-gR2 across CD14+ Monocytes **(g-h)** and their associations with outcome across standard-risk patients. Box and violin plots comparing the per-patient average expression of IFN-g in T cells **(e)** or IFN-gR2 in CD14+ Monocytes **(g)** across standard-risk NP and standard-risk RP patients. Each dot represents individual patient. **(f,h)** Kaplan Meier curves displaying the association between expression of IFN-g in T cells **(f)** or expression of IFN-gR2 in CD14+ Monocytes and progression-free survival within standard-risk patients. “High” group corresponds to patients with an expression of IFN-g above the median, while “Low” corresponds to patients below the median. The hazard ratio and P value of a Cox regression model fitted to the IFN-g expression are also calculated for each analysis. **(i)** Heatmap displaying the normalized average AUC score for various transcriptional regulons on selected myeloid populations. Additional columns include the hazard ratio, along with the p-value, estimated from a Cox proportionality hazard model fit on average patient AUC scores (categorized into high and low activity). **(j)** Survival plots display the corresponding survival associations with the expression of the corresponding ligand within the myeloid compartment of patients, where high E2F8 regulon expression *(bottom)* and low IRF7 *(top)* regulon expression are associated with poor outcomes. **(k)** Feature plots showing the per cell AUC values for the IRF7 *(left)* and E2F8 *(right)* transcription factor regulons derived from SCENIC analysis across key myeloid populations. Blue color indicates low activity (or AUC), while red color indicates high activity.

Further analysis of standard-risk patients who experienced rapid disease progression (SR-RP, n = 19) compared to those with non-progression (SR-NP, n = 40) showed significant enrichment of myeloid populations paired with depletion of B lymphoid populations and a decreased proportion of CD8^+^ T cells relative to CD4^+^ T cells (**Figure 5a, Supplemental Figure 12**). Differentiated T cell populations (CD8_Teff, CD8_Teff_HLA) showed a significant association with SR-RP patients (p < 0.05, **Figure 5a, Supplemental Figure 13**). Uniquely, the SR-RP cohort showed significant enrichment of rare CD4^+^ cytotoxic T cells (*P* = 0.029), suggesting a putative role in rapid disease progression in standard-risk patients (**Figure 5a**). Additionally, we also observed significant reductions in B cell progenitors in the SR-RP group (P=0.044, **Supplemental Figure 13**). Paired with the significant shift from B lymphoid populations enriched in the NP cohort to myeloid populations in the RP cohort (**Figure 4b**), this suggested that changes in the immune composition may be traced back to altered hematopoiesis within the BMME. Therefore, we investigated the differentially expressed genes within the HSC cluster, revealing that the HSC cluster indeed reflected a shift toward myelopoiesis in the rapid progressors, with over-expression of myeloid lineage commitment markers, while the non-progressors exhibited a slight over-expression of lymphoid lineage commitment markers, such as SOX4 (**Figure 5b**).

### Cellular communication analysis depicts IFN-**y** driven proinflammatory and immunosuppressive changes in patients with poor outcomes

To explore potential BMME signaling changes associated with cytogenetic risk and disease progression, we investigated intercellular communication patterns, revealing several key pathways in outcome-associated subpopulations (**Figure 5c**). MHC-II expression was enriched in antigen-presenting cells (B cells, M2_Macro, cDCs), which was associated with non-progression (**Figure 4e**) and standard-risk (**Figure 3h**) cohorts, pointing towards an improved adaptive immune response in these groups. We also observed increased expression of interferon-gamma (IFN-γ) in CD8^+^ T effector populations (CD8_Teff_TNF, CD8_Teff_HLA, CD8_Tem, **Figure 5c-d**, **Figure 2f-g**), in CD4^+^ cytotoxic populations (CD4_CTL), and in CMV adaptive-like NK cells (NK_adp). Higher IFN-γ receptor expression was also found in the rapid progressor-associated classical and inflammatory monocyte clusters (Macro/Mono, CD14+Mono_ProInflam). Markedly, CD8_Teff_HLA cells were significantly associated with rapid progression and high risk (**Figure 4f**, **Figure 3d**), suggesting that IFN-γ signaling in the BME may contribute to the inflammatory alterations in monocytes of rapid progressors. Notably, the RP-associated cluster (Macro/Mono, **Figure 4c**, **Supplemental Figure 7**) was also found to express *BAG6*, an inhibitor of NK-mediated cytotoxicity in its soluble form^28^, and well-documented molecules in MM oncogenesis and progression, thrombospondin (THBS), and APRIL. In contrast, the interferon-associated monocyte cluster, associated with SR and NP patients (**Figure 3f**, **Figure 4c**), was found to highly express BAFF, an essential promoter of B cell survival and terminal differentiation^29^. BAFF can bind to TACI expressed on plasma cells, though it has a much higher affinity to BAFF-R expressed on the mature B cell populations, which are more abundant in non-progressors (**Figure 4b**).

Given that IFN-γ overexpression in the T cell compartment correlates with rapid disease progression, we further investigated IFN-γ expression in standard-risk patients and its relationship to outcomes. Standard-risk rapid progressors displayed significantly higher average IFN-γ expression across their T cell compartment (**Figure 5e**), which was associated with poor outcomes (**Figure 5f**). Furthermore, CD14^+^ monocytes of standard-risk rapid progressors had significantly higher IFN-γ receptor (i.e., *IFNGR2*) expression (**Figure 5g**), which was associated with poor outcomes in standard-risk patients (**Figure 5h**). These findings appeared to indicate that heightened IFN-γ expression before therapy may be a prognostic indicator of poor outcomes.

In a systems biology analysis, we further investigated gene regulatory networks, particularly focusing on myeloid subpopulations associated with high-risk and rapid disease progression (e.g., CD14+Mono_IFN) and identified enrichment of regulatory networks for IRF2, IRF7, IRF9, and STAT1 transcription factors (TFs) (**Figure 5i-k, Supplemental Figure 14, Supplemental Table 5**). These TFs are regulated by type I interferon (IFN-α) and promote the transcription of IFN-α stimulated genes, including *ISG15*^30,31^. Examining the survival association of IRF7-regulon activity within the myeloid compartment, we observed that patients with increased IRF7-regulon activity exhibited better outcomes (CoxPH *P* < 0.01) (**Figure 5j**). Additionally, regulatory networks of cell proliferation related *E2F1* and *E2F8* TFs were enriched in the GMP population elevated in patients with rapid progression (**Figure 5i,k**). Increased E2F8-regulon activity was linked with poor survival outcomes within the myeloid compartment, aligning with our previous observation of increased myelopoiesis in patients with rapid progression (CoxPH *P* < 0.05) (**Figure 5j**).

### Integrating baseline immune signatures with cytogenetic risk improves our ability to predict outcomes

Finally, we assessed the ability of immune cell clusters to predict disease progression in a univariate and multivariable framework (age, sex, stage, and cytogenetic risk) by employing a bootstrapping approach and three different statistical methods (**Figure 6a**). The performance of these predictive models was assessed using the area under the receiver operating characteristic curve (AUC).

**Figure 6.**
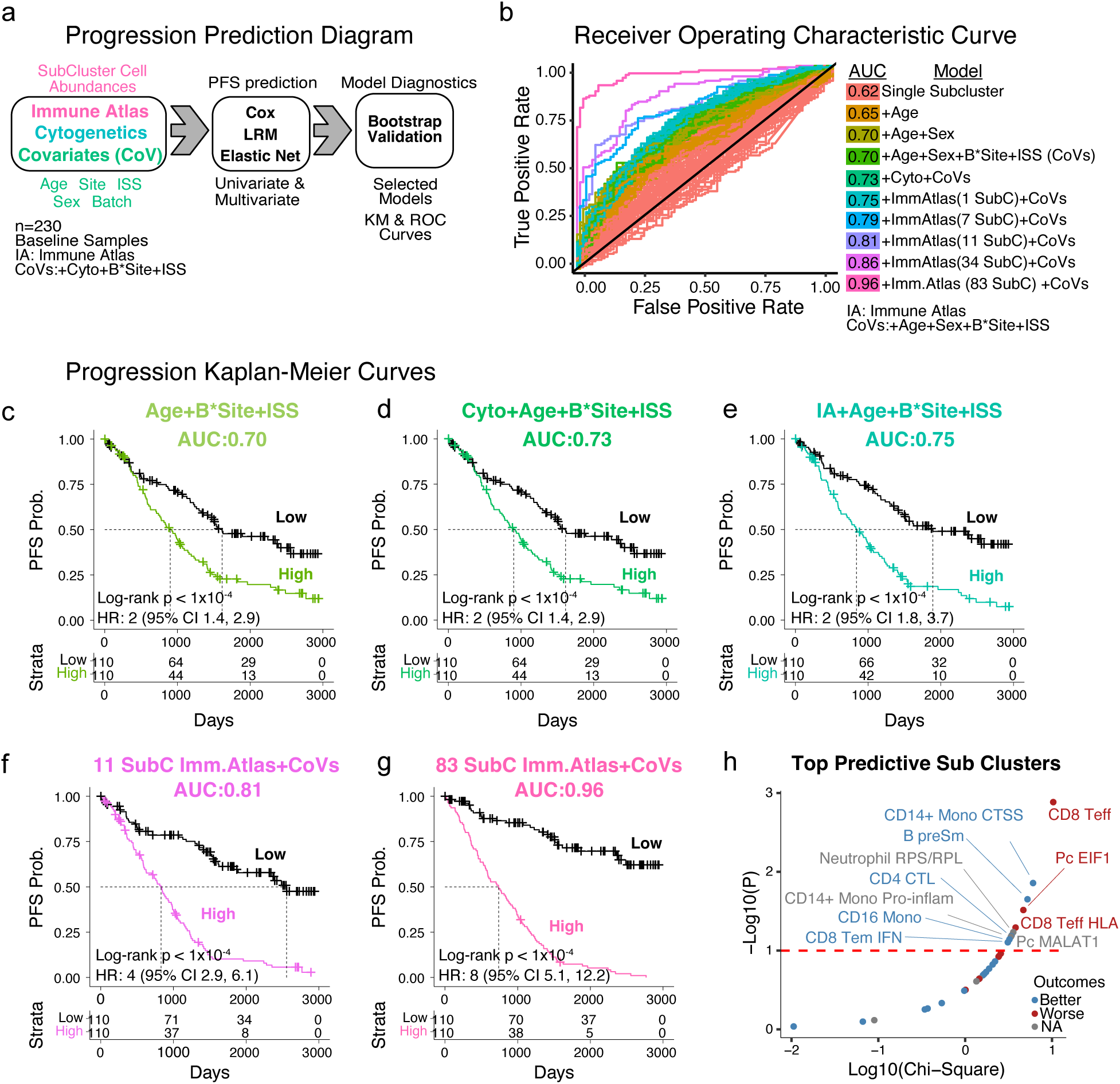
Prediction of MM progression by integration of cytogenetics risk along with immune signatures. **(a)** Shows a diagram with the type of variables that were tested (immune signatures, cytogenetics, and clinical variables (covariates)) followed by the three regression strategies used (elastic net, logistic regression, and Cox). Finally, bootstrap validation was used for model selection **(b)** Receiver operating characteristic (ROC) curves for progression prediction models based on single clusters, clinical variables, and cytogenetics or immune atlas variables alone and in combination are shown colored based on the specific group of models. The labels indicate SubC = SubClusters, CoV = Covariates and these include age, Batch, Site, ISS and Cytogenetic. Kaplan-Meier curves showing the separation of patients with high or low scores for prediction of PFS are shown for **(c)** Age, ISS stage, and batch **(d)** cytogenetics, age, ISS stage, and batch, **(e)** Immune Atlas Signatures, Age, ISS stage, and batch. Kaplan-Meier curves show the separation of patients when cytogenetic risk scores are combined with the **(f)** best 11 predictive immune atlas subclusters or **(g)** with all 83 subclusters **(h)** importance of immune subclusters for predicting the progression. The clusters with better and poor MM outcomes are shown with blue and red colors respectively.

Clinical data alone yielded an AUC value of 0.7 in predicting PFS (**Figure 6b, c**). Incorporating cytogenetic risk with clinical variables (stage and demographics) increased the PFS prediction to 0.73 **(Figure 6b, d)**. Similarly, the predictive power of any single immune subcluster (SubC) combined with clinical variables only marginally improved prediction (AUC = 0.75, **Figure 6b,e**). Iterative feature selection combining subsets of BMME immune clusters with clinical covariates and cytogenetics improved the average AUC for predicting disease progression (**Figure 6b, f-g**). Specifically, combining immune atlas clusters, clinical variables, and cytogenetics resulted in AUC values ranging from 0.75 (any given subcluster) to 0.96 (all subclusters), contingent on the number of subclusters considered during modeling (**Figure 6b,g**). This substantial increase in AUC by combining clinical and immune features (**Supplemental Figure 15a, b**) further highlights the importance of the BMME. Although marginal differences in AUC were observed among individual compartments alone, integrative models showcased a significant advantage over simpler models (**Supplemental Figure 15a-d**). Finally, we identified the most relevant 11 subclusters selected using an elbow test on predictive power vs number of clusters, resulting in a high precision/recall model (AUC:0.81, **Figure 6b,f,h**). This model for stratifying patients into progressors and non-progressors (**Figure 6b,f,h**) included differentiated cytotoxic populations (CD8_Teff, CD8_Teff_HLA) along with inflammatory myeloid populations (CD14+Mono_ProInflam) (**Figure 6h**). This exploratory analysis suggests that the integration of BMME information with clinical and genetic variables may enhance risk stratification and outcome prediction, yet it still requires independent validation.

## DISCUSSION

In this study, we have produced a large comprehensive multiple myeloma single-cell Immune Atlas focused on mapping the myeloma BMME. By standardizing and harmonizing a workflow across multiple centers, we successfully profiled 361 myeloma samples, resulting in >1.1 million cells from the BMME, capturing numerous biologically relevant cell states, including rare subtypes harder to detect in smaller studies, such as cytotoxic CD4^+^ T cells, mast cells, HSCs, and fibroblasts. The atlas also enabled deciphering BMME variations among patients with diverse cytogenetic risk profiles and clinical outcomes; notably these patients were not treated with recently approved T-cell immunotherapies, suggesting that immune function has a broad, treatment-independent role in suppressing tumor growth.

The T cell compartment of rapid progressors and high-risk patients displayed an accumulation of terminally differentiated CD8^+^ effector T cells, specifically dysfunctional cytotoxic cells, with reduced naïve populations (**Figure 7**). This state is sometimes referred to as immunosenescence, as the expanded effector population expresses low levels of cytotoxicity-related genes along with the inhibitory KIR KLRG1, and lacks costimulatory receptors CD27 and CD28, resulting in poor antigen-mediated proliferation capabilities^32–36^. Additionally, some studies have indicated that the immunomodulatory effects of drugs such as IMiDs may act through the costimulatory CD28^+^ pathway^37^, and therefore, this population may show a diminished response to standard first-line therapies, potentially leading to poorer outcomes. Depletion of the naïve pool can be driven both by thymic involution^38,39^ or by antigenic pressure driven from either the myeloma cells or by other chronic infections such as CMV or EBV^40–42^. Impairment of naïve T cells reduces the TCR repertoire clonality^36,43^, which typically is associated with worse outcomes in various malignancies^44,45^. Therapies that stress the immune environment, including many of the immunotherapies used for MM, such as IMiDs and ASCT, can also drive similar alterations^46,47^, which in the elderly may be slow to recover due to impairment of new naïve T cell reconstitution^36,38,39^. Cytotoxic cells are critical for clearing the malignant populations; however, the accumulation of these differentiated T cell populations may reflect a less reactive T cell compartment. Determining if this influences response to immunotherapies will require profiling that specific patient cohort. Additionally, these populations contribute to the inflammatory microenvironment through the production of cytokines such as IFN-γ, which we observed highly expressed in the HLADR^+^, CD28^neg^ population associated with both high-risk and poor outcomes. Unlike exhaustion, it is not well understood if this senescent state can be reversed, though some studies have indicated it may be possible^48^. Given that immune therapies could aggravate the T cell compartment imbalance, it may be better to utilize more targeted therapies, such as bispecific antibodies, and CAR-T cells in the first line of therapy, rather than only for relapsed disease.

**Figure 7:**
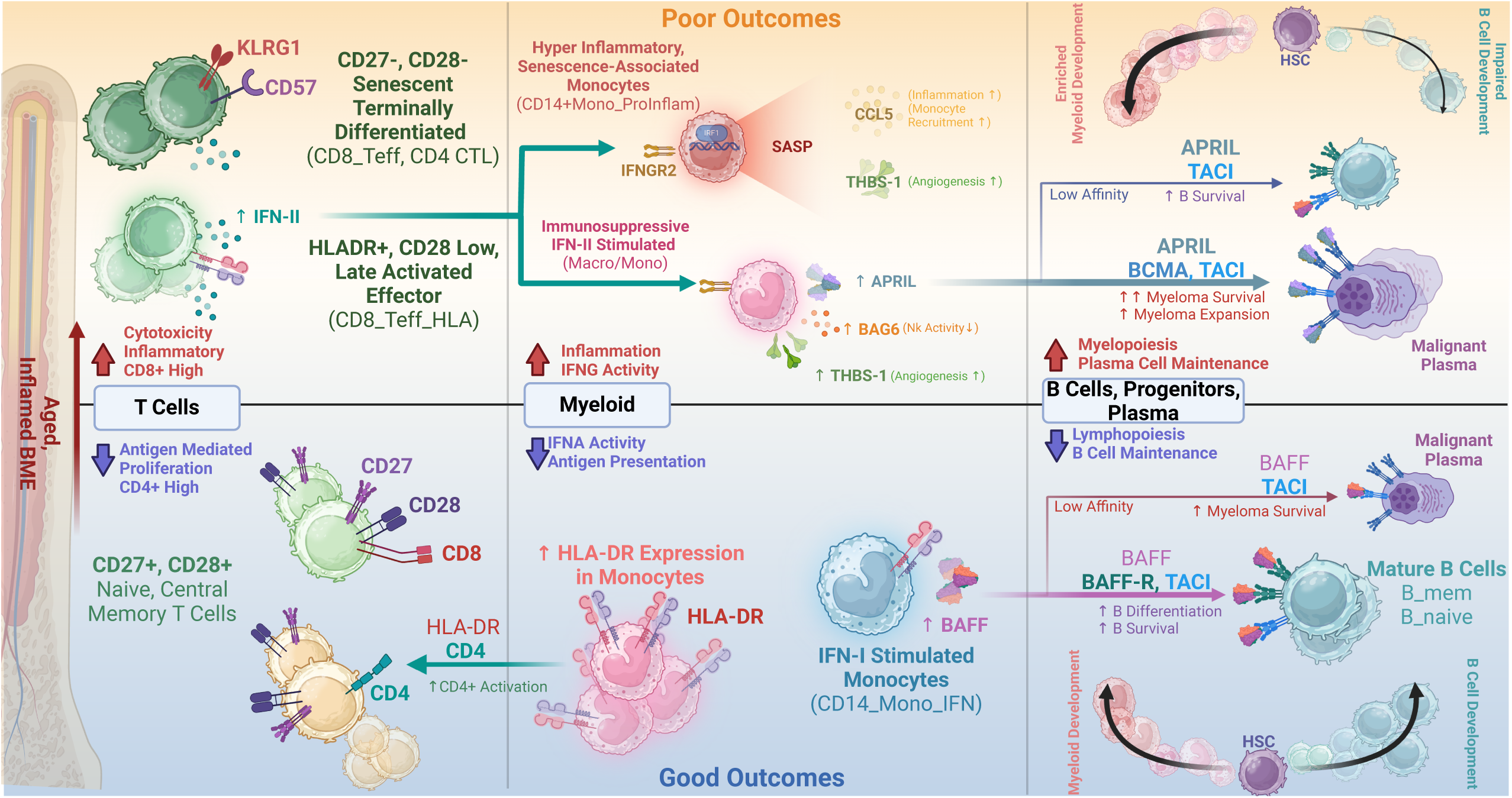
Inflammatory remodeling defines the BMME in rapidly progressing MM. Summary of the key cellular subtypes and signaling pathways comprising the MM BMME and their association with patient outcomes. Within the aging bone marrow, a state of chronic inflammation, known as ‘inflammaging’, results in altered lymphoid and myeloid cell populations, enabling immune escape of malignant plasma cells. Within the T cell compartment, MM patients with poor outcomes exhibit a shift toward immunosenescent and late-activated CD8+ T cells, producing IFN-II that drives senescence-associated and immunosuppressive phenotypes in myeloid compartment. In contrast, MM patients with better outcomes display highly proliferative naïve and central memory CD8+ T cell subsets, in addition to enriched T helper populations driven by increased MHC-II antigen presentation among myeloid cells. T cell and myeloid populations in these patients appear to be stimulated by type I interferons, in contrast to patients with poor outcomes exhibiting enrichment of IFN-II signaling. This difference in interferon stimulation appears to be linked to patient outcomes, in part, through the differential expression of BAFF by IFN-I-stimulated monocytes and APRIL by IFN-II-stimulated monocytes. Notably, BAFF preferentially binds to mature B cells to promote survival, potentially enhancing B cell-mediated immunity and leading to improved outcomes. Conversely, APRIL has been shown to inhibit memory B cell function and promote malignant plasma cell survival. This dysregulation is further highlighted in the shift from B cell development towards increased myelopoiesis in patients with poor outcomes. Created with Biorender.

In addition to T cell alterations, rapid progressors displayed a shift towards myelopoiesis in the bone marrow, reflected by general depletion of the B cell compartment, including the bone marrow native immature populations, compensated by the increased myeloid compartment (**Figure 7**). Myelopoiesis in the bone marrow can be driven by stress or inflammation that drives HSCs to differentiate toward myeloid lineages at a higher frequency^49^. Myeloid cells can be beneficial as non-progressors, and standard risk patients showed enrichment of MHC-II signaling in their CD14+ monocytes, which are known to promote antigen-driven activation of T cells^50^. However, these cells can also be a source of inflammatory cytokines promoting tumor survival, immunosuppression, or angiogenesis, as observed in rapid progressors, displaying enrichment of senescent-associated secretory profile factors, including IL-8, CCL3, or IL-1B^51^. The enrichment of these inflammatory factors may be related to IFN-γ produced by the expanded CD8^+^ effector T cell populations, as the inflamed myeloid cells both express the receptor *IFNGR2* and the transcription factor *IRF1*, which is associated with IFN-γ activity^52^. Additionally, other studies have demonstrated that IFN-γ can promote an inflammatory myeloid phenotype by inhibiting IL-10^53^, which would normally act as a negative feedback regulatory mechanism. This is distinct from the ISG15-related interferon activity observed in non-progressors and standard-risk patients, typically driven by type-I IFNs^54^.

Cell-cell communication analysis identified both BAFF (*TNFSF13B*) and APRIL *(TNFSF13*) expression in the myeloid compartment. BAFF expression was primarily associated with myeloid populations enriched in non-progressors such as interferon-associated monocytes. BAFF can bind to TACI expressed on plasma cells, though it has a much higher affinity to BAFF-R expressed in mature B cell populations to promote their differentiation and survival. Conversely, APRIL was most strongly associated with the Macro/Mono population enriched in rapid progressors. APRIL is known to bind to TACI (*TNFRSF13B*) on malignant plasma cells, promoting their survival and MM progression^55,56^.

Cytogenetics alone demonstrated positive predictive capabilities, yet integrating information from the BMME could enhance stratification and guide optimal therapeutic selection. We observed that the prevalence of differentiated BMME immune cell populations can predict outcomes with good accuracy in our cohort regardless of cytogenetics. Importantly, combining tumor cytogenetics, and clinical variables with the BMME cellular immune signatures significantly improved the accuracy in stratifying myeloma outcomes. Patients with immunosenescent and inflamed bone marrow microenvironment might have poor overall or event-free survival even with a favorable genetic makeup. Therefore, we posit that future treatments targeting the immune microenvironment could improve outcomes of myeloma. This observation can elevate the significance of capturing the BMME as a prognostic marker for MM. Increasing the studies capturing such information at the cohort scale could enable us to establish a new generation of comprehensive risk scores or the derivation of simplified lower-cost assays that focus only on the most informative populations. Furthermore, these data may identify ancillary therapeutic targets that improve the efficacy of current treatment strategies and may contribute to rationally designed, personalized treatment regimens based on both the tumor and the immune microenvironment.

The study creates an extensive and comprehensive resource to map the granular cellular landscape of myeloma from baseline samples. This lays the foundation for future studies to explore how the BMME is altered by various immunostimulatory and targeted treatments. However, the study has multiple limitations, including only studying gene expression, and does not include any proteomic or functional profiling. Additionally, our study covers immune cells comprehensively but with a sparse representation of non-immune BM stromal populations, limiting our capability to evaluate their associations with outcomes. More targeted studies may still be required to understand the role of BM stromal populations in MM outcomes.

This single-cell dataset extends upon wealth of patient information in the MMRF CoMMpass study including tumor gene expression, DNA copy number abnormalities translocations and mutations as well as long-term clinical data^6^. This study is ongoing and will continue to generate additional BMME immune data. As we demonstrated in this investigation, combining orthogonal information, in this case genomic and cell abundance profiles can uncover foundational information for the disease with direct translational implications and applicability. This paradigm can enable us to better understand the combinations of factors that influence outcomes in multiple myeloma and move closer to the goal of optimizing therapy for each patient to ensure the best outcomes.

## Methods

### Materials Availability

This study did not generate new unique reagents.

### Data availability

All the single-cell raw data is available at MMRF’s VLAB shared resource. Requests to access these data will be reviewed by data access committee at MMRF and any data shared will be released under a data transfer agreement that will protect the identities of patients involved in the study.

### Code availability

All the code used for data analysis and generation of figures will be available on the MMRF Immune Atlas Consortium GitHub (https://github.com/theMMRF/MMRF_ImmuneAtlas)

### Ethics approval and participant consent

All samples for the study were obtained from the MMRF CoMMpass clinical trial (NCT01454297). Procedures involving human participants as part of this trial were performed by the ethical standards of the MMRF research committee. Written informed consent was obtained from patients for the collection and analysis of samples and clinical information by the MMRF. The Institutional Review Board at each participating medical center approved the study protocol. The list of participating institutes that have approved the study protocol is available at ClinicalTrials.gov (NCT01454297).

### Experimental model and human subject details

A total of 361 CD138^neg^ MM bone marrow mononuclear cells (BMMC) samples were collected from multiple myeloma patients (n = 263) enrolled in the MMRF CoMMpass study (NCT01454297). Patients enrolled in the study were monitored via quarterly check-ins for up to eight years following initial disease diagnosis. All patients were required to be eligible for either standard triplet therapy (immunomodulatory drug, proteasome inhibitor, glucocorticoid) or doublet therapy. Most patients received triplet therapy in their first line of therapy. The details of patients’ information are available in **Supplemental Table 1**. Samples were acquired pre-therapy (baseline) and post-therapy (relapse or remission), and then processed at four institutions: Emory University, Mayo Clinic Rochester, Mount Sinai School of Medicine, and Washington University.

### CD138^neg^ cells isolation and cryopreservation of cell samples

Bone marrow aspirates from the Multiple Myeloma Research Consortium (MMRC) tissue bank were separated into CD138^pos^ (myeloma cells) and CD138^neg^ (immune, bone marrow cells) fractions using immunomagnetic cells selection targeting CD138 surface expression (automated RoboSep and manual EasySep from StemCell Technologies Inc.). Following magnetic separation, the CD138^neg^ cell fractions were viably frozen. Briefly, the CD138^neg^ cells were centrifuged at 400 × g for 5 min. The resulting cell pellet was resuspended in freezing media consisting of 90% FCS and 10% DSMO at a concentration of 5–30 million cells per ml in multiple aliquots. Cell concentrations and aliquot locations were documented, before storing in liquid nitrogen for future studies.

### Single-cell RNA-seq sample preparation, library construction, and sequencing

To generate high-quality and comparable single-cell data, we developed a highly detailed single-cell protocol based on our pilot studies^12,19,20^ for implementation across centers and performed profiling using Single Cell 3’ profiling (10X Genomics Inc). Briefly for single-cell RNA-seq, aliquots of the CD138^neg^ BMME samples were thawed quickly in 37°C water bath. Cells were washed with a warm medium and pelleted by spinning at 370g for 5 minutes at 4°C. The cell pellet was resuspended in ice-cold phosphate buffer saline (PBS) with 1% BSA and cell viability was measured. If cell viability was < 90%, dead cell removal was performed using the Dead Cell Removal Kit (Miltenyi Biotec Inc). The cell pellet was resuspended in 100 µL of dead cell removal microbeads solution and incubated at room temperature for 15 minutes. Magnetic removal of labeled dead cells was performed using the MS column or autoMACS^®^ Pro Separator. The eluted supernatant containing the live cells was pelleted by centrifugation at 370g for 5 minutes at 4°C. Cells were finally resuspended in ice-cold PBS containing 1.0% BSA. In some samples, 100-150 cells from murine sarcoma lines (NIH/3T3 – CRL-1658, ATCC) were spiked into the final human single-cell suspension to assess batch effects across centers. The cells were loaded onto the 10x Genomics Chromium Controller according to the manufacturer’s instructions, followed by RT-PCR, cDNA amplification, and library preparation using the Chromium Next GEM Single Cell 3′ GEM, Library & Gel Bead Kit v2.1. Briefly, approximately 8,000 cells were partitioned into nanoliter droplets to achieve single-cell resolution for a maximum of 5,000 individual cells/sample. The resulting cDNA was tagged with a common 16nt cell barcode and 10nt unique molecular identifier (UMI) during the RT reaction. Full-length cDNA from poly-A mRNA transcripts was enzymatically fragmented and size selected to optimize the cDNA amplicon size (∼400 bp) for library construction as per recommendations from 10X Genomics Inc. The concentration of the single-cell library was accurately determined through qPCR (Kapa Biosystems) to produce cluster counts appropriate for the paired-end sequencing using NovaSeq 6000 platforms (Illumina). The sequencing data was generated by targeting between 25-50K reads/cell, which provided gene expression profiles of 1,000-4,000 transcripts per cell.

### Single-cell RNA-seq genome alignment and quality control

For analysis of single-cell RNA-seq samples, Cell Ranger (v6.0.1, 10x Genomics Inc) was used for demultiplexing sequence data into FASTQ files, aligning reads to the human genome (GRCh38), and generating gene-by-cell UMI count matrices. Empty droplets were removed using DropletUtils^57^ (v1.14.2) (FDR <0.001).

Ambient RNA was removed using CellBender^58^ (v0.3.0) (FPR = 0.01). For quality control, cells with <1000 UMI reads, <200 unique genes, or > 20% of UMIs mapped to mitochondrial genes were filtered out using Seurat^59^ (v4.3). Harmony^60^ (v0.1) was implemented to mitigate batch effects from processing sites and shipment batches in the resulting cell clusters and embeddings. For a small subset of downstream analyses that directly operate on the count matrix and do not support controlling for a batch covariate, such as CellChat or SCENIC, a corrected count matrix was generated as described in the next section.

### Batch-corrected count matrices for gene regulatory and cellular communication analysis

#### Batch effect estimation

First, the Poisson Pearson residuals were computed for each gene across all cells^61^. Genes with zero UMI counts across all cells were excluded from further steps. For the remaining genes, the proportion of variance explained by batch in the Pearson residuals was estimated using the R-squared from a linear regression model^62^. Genes where the batch explained less than 1% of the variance were removed to avoid overcorrection.

#### Batch corrected counts

The reference count distribution for each gene affected by batch was modeled as either Poisson (when the mean was equal to the variance) or negative binomial. The Poisson parameter was estimated using the maximum likelihood estimator, while the negative binomial mean and dispersion parameters were estimated using a Gamma-Poisson generalized linear model^63^. The batch correction was performed in two steps.

1) Scaling and centering the Pearson residuals using the batch-level means and standard deviations to account for the differences between batches. 2) Transforming the standardized Pearson residuals onto the probability scale using the empirical-distribution function^64^ and then to the batch corrected counts using the quantile function of the reference Poisson or negative binomial distribution. A pseudo-count of 1 and the original zeros observed in the uncorrected UMI counts were restored to preserve the observed sparsity^65^.

### Mouse cell removal

To identify mouse cells, we additionally mapped the raw data to the human (GRCh38) and mouse (mm10) combined reference genome. We removed clusters with more than 80% of cells having less than 95% reads mapped to the human reference genome. Two samples with over 65% of cells being identified as mouse cells were removed from further analysis. Cell barcodes corresponding to mouse cells were removed from the all-sample merged object.

### Clustering and cell annotation

Following the removal of mouse cells, raw counts were log normalized (scale factor = 10000) using Seurat^59^ (v4.3). The first 25 principal components derived from principal component analysis (PCA) were computed from the top 3,000 variable genes to reduce data dimensionality. Harmony was applied to these principal components to generate batch-corrected embeddings, where each combination of processing center and shipment batch was considered an independent variable. To cluster cells of similar transcriptome profile, Louvain clustering was performed on the batch-corrected harmony embeddings using Seurat’s ‘FindClusters’ function^59^. Clusters were visualized using Uniform Manifold Approximation and Projection (UMAP). Clusters were aggregated into five major connected components called compartments based on their separability on the UMAP. To annotate these compartments, a combination of SingleR^66^ and cell type/subtype specific marker expression was used. The identified compartments included ‘T/NK’ (T cells and Natural Killer cells), ‘B-Ery’ (B cells, CD34^+^ populations, erythroblasts), ‘Myeloid’ (Monocytes, Neutrophils, Dendritic cells), ‘Plasma’ (Plasma cells’), and ‘Ery’ (Erythrocytes). A small independent cluster of fibroblasts (946 cells) was observed, in the initial UMAP, and was not included in any compartment.

More precise annotation of individual cell compartments was performed separately by repeating the above process on each compartment, leveraging variable genes specific to each compartment. Due to the highly patient-specific nature of myeloma populations, batch correction in the plasma compartment was done per aliquot instead of per batch. Each cluster was manually annotated based on the expression of canonical markers or top genes of the clusters. While annotating cells, if a possible subset was identified within a given cluster based on marker expression, further sub clustering was performed specific to that cluster using the same procedure. Multiple resolutions were assessed, with the final sub clustering used being the result that isolated the subpopulation of interest, while minimizing the formation of minor or patient-specific clusters.

### Single-cell mutation mapping and CNV inference

To better understand tumor heterogeneity and malignancy of plasma cell populations, we profiled mutations and copy number changes of plasma cells. First, we utilized a mutation mapping strategy to detect mutations within each cell by looking for reads supporting the reference or variant alleles at variant sites in mapped reads from scRNA-seq BAM files. This is achieved by leveraging high-confidence somatic mutations derived from whole-exome sequencing (WES) data from the same patient. The code for mutation mapping is available on GitHub (https://github.com/ding-lab/10Xmapping). Furthermore, we used inferCNV (v.0.8.2, https://github.com/broadinstitute/inferCNV) (with default parameters) to identify sample-level chromosomal copy number variations (CNVs) of plasma cells, using the immune cells as reference normal set.

### Doublet detection

Doublets were identified by flagging clusters with high doublet proportions as predicted via DoubletFinder^67^, Scrublet^68^ (v0.2.3), and Pegasus (https://github.com/lilab-bcb/pegasus) (v1.8.1). Scrublet was used to detect doublets with the expected doublet rate set at 0.06 and thresholded at 0.2. Doublet-enriched clusters, characterized by at least two methods (FDR < 0.05, Fisher’s exact test), were manually reviewed and marked as doublets accordingly. Characteristics considered when reviewing doublet-enriched clusters include the simultaneous, high expression of canonical markers from unrelated lineages (e.g., T cell markers *CD3*, *CD8A*, *GZMK* and myeloid markers *LYZ*, *CST3*, *CD14*), or UMI counts disproportionately high relative to similar cell types. Seventeen cell clusters (Cells n=74,282) were flagged as doublets and were omitted from downstream differential expression, abundance, and trajectory analysis.

### Differential expression among cell types and clinical groups

Differential expression analysis was performed using linear (as implemented in limma^69^) and mixed effect models (as implemented in lme4^70^) R packages to identify markers enriched in each population, or between clinical groups of interest. Our models were adjusted for covariates including clinically relevant age, sex, stage, and technical factors like sequencing site and batch. Significance was determined using moderated t-test statistics on the log-normalized expression. P values are adjusted for multiple comparisons using Benjamini-Hochberg correction. Log fold change was computed by performing a logarithmic transformation on the ratio of the arithmetic mean expression for cells between groups being compared.

### Differential abundance of cell types and subtypes

Differential abundance was performed by computing the per-patient cell-type proportion across all cells and within a specified compartment using a Dirichlet regression model^71^ and Wilcoxon Rank Sum tests. Dirichlet regression models were used when the observed cell abundances were different across batches with respect to a covariate of interest (e.g. Risk) whereas Wilcoxon Rank Sum tests were used when the association between batch and the covariate of interest was not significant. Dirichlet regression model was implemented using DirichletReg R package^71^. Wilcoxon Rank Sum tests were implemented with the rstatix package (https://cran.r-project.org/web//packages//rstatix/), with the rank-biserial correlation coefficient estimated using the effectsize package^72^. P values less than 0.05 was considered statistically significant.

### Patient stratification based on time interval to disease progression

Patients in the CoMMpass study had regular three-month check-ins in which clinical parameters were evaluated following therapy. The day of disease progression was identified using standard IMWG criteria. Progression data used in this study was derived from the IA22 CoMMpass clinical metadata release. Patients were categorized into discrete progression groups based on their progression-free survival, and the duration of time the patient was enrolled in the study. The extreme categories of rapid progressing (RP) and non-progressing (NP) use cut-offs matching those of our pilot study^12^. Rapid-progressing patients are those with a progression event within 18 months of therapy (PFS < 18 months). Non-progressing patients include those who had no progression event for at least four years following therapy (PFS > 4 years). Progressing patients are those who have a documented progression event between 18 months and 4 years (PFS > 18 months, PFS < 4 years). Progressing patients are those who experience a documented progression event from 18 months to 4 years. Incomplete patients are individuals who exited the study before four years of disease diagnosis without experiencing a progression event.

### Cytogenetic risk-based stratification of patient samples

The cytogenetic risk categorization was defined using translocation data or copy number data derived from CD138^pos^ whole genome sequencing results included in the IA21 CoMMpass metadata release. Thresholds for calling mutation events in the MMRF CoMMpass data are based off of the work by Skerget et al.^11^ Patients at high risk were defined with one of the six following mutation events: del17p13, t(14;16)[MAF], t(8;14)[MAFA], t(14;20)[MAFB], t(4;14)[WHSC1/MMSET/NSD2], 1q gain. This extends the definition proposed by Skerget et al., by incorporating 1q gain. Patients with none of these six events were considered standard risk. Patients with partial mutation data, such as having only translocation or only copy number data, can be classified as high-risk if a high-risk mutation is present in the available data but otherwise were excluded from downstream analyses involving risk-based stratification.

### Prediction of patient progression based on immune cell abundances, cytogenetics, and demographics

To assess the progression prediction capability of the immune signatures alone as well as in combination with clinical variables, we developed and evaluated multiple classifiers. We have evaluated the predictive power of clusters containing cells from at least 50% of samples, resulting in the usage of 83 subclusters. Subsequently, the cell frequencies of these subclusters were utilized to construct both univariate and multivariate models employing three distinct methods: Cox regression, logistic linear regression (LRM), and elastic net regression. Internal validation using bootstrap was used to verify the robustness of results.

The survival curves based on the CD8 Teff HLA+ and CD14+ IFN+ Monocyte abundances are based on a Cox proportional hazards model regressing the overall survival on the Pearson residuals from a Dirichlet regression model with batch as the covariate. The optimal cut-off to separate the patients at baseline in low and high groups was determined using maximally selected rank statistics^73,74^ implemented in the surv_cutpoint function from the survminer package [https://CRAN.R-project.org/package=survminer]. For the elastic net regression models, a sensitivity analysis of the coefficients was performed to facilitate feature selection and the identification of pertinent features. Likewise, features were selected from LRM and Cox models using p-value filtering. Our modeling approach was designed to assess individual or multiple subclusters, integrated with additional variables such as age, sex, disease stage (ISS), and the cytogenetic risk descriptor mentioned earlier.

To ensure the robustness of our models, a bootstrap validation approach was implemented, yielding bias-corrected indexes specific to each model type. Model performance metrics, including Somers’ index, Dxy, and diagnostic statistics were computed using Harrel’s rms R package. The following R packages glmnet, surv, coxph, and rms packages were employed for identifying and testing predictions of disease progression. Visualization was carried out using R packages such as ggplot, tidyverse, pheatmap, survminer, and gtsummary.

### Cell transition trajectory analysis

Pseudotemporal ordering of cells was performed using the Slingshot R package^75^. Cells with a known biological lineage were isolated from all other populations (e.g., CD8 T Cells), and doublet and mitochondrial-enriched populations were excluded. New variable features and batch-corrected embedding components were computed as described above in the “Cluster and cell annotations” section. Slingshot was performed on the first 25 batch-corrected harmony embeddings. If an identified progenitor, or less-differentiated population was detected through annotation, this cluster was designated as the ‘start cluster’ for trajectory analysis. Pseudotimes for all clusters, representing the distance along the trajectory from the starting cluster were calculated.

### Survival Analysis

The patients were categorized based on clinical characteristics and risk groups. To analyze survival outcomes, we calculated the probabilities of progression-free survival (PFS) and overall survival (OS), and generated Kaplan-Meier curves for both groups, using the survival^76^ (v.3.2.7) and survminer (v.0.4.9) packages in R. The PFS and OS data were derived from the IA22 CoMMpass clinical metadata release. Patients who left the study prior to any follow-up appointments, or patients with large delays in the start of therapy, were excluded from survival analysis. Cox proportional hazard models were used to determine the clinical characteristics that had the most significant and independent impact on patient survival. The significant clinical features identified from the Kaplan-Meier curves were then included in the models to assess their individual contribution to survival while adjusting for factors such as age, sex, and race.

### Longitudinal single-cell analysis in relapse patients

Longitudinal analysis of single-cell data was performed on patients with samples taken at relapse or progression of the disease. Fold change differences in cell proportions were calculated by dividing the cell proportions at the last time point by the cell proportions at the baseline or first time point. The significance of fold change (log2 scale) was calculated using the paired Wilcoxon-signed rank test. To better understand the temporal changes in cellular composition between cytogenetic groups, linear models were used to estimate the change in cellular proportion over time. Visit intervals were converted to the number of months from diagnosis. A variance-stabilizing arcsin square root transformation was applied to individual cell type proportions to account for the naturally-occurring mean-variance relationship which arises in proportion data bounded between 0 and 1^77,78^. The lm function in R was used to fit the linear models between the arcsin-transformed proportion and visit month for each cell type split by patients categorized as standard- or high-risk. The slope of the linear model indicates the direction and magnitude of the temporal relationship between cellular abundance and time. P-values regarding the significance of the association across all cell types were adjusted using the BH procedure.

### Pathway activity analysis and signature scoring

Gene-set enrichment analysis (GSEA) was performed to identify the pathways enriched across cell types or clinical groups. The GSEA analysis function gsePathway from ReactomePA^79^ was used to compute normalized enrichment scores derived from an ordered gene list. Gene lists were ordered by log fold-change computed via limma trend, correcting for processing site and batch among cell types or clinical groups (See, Differential Expression). To evaluate the enrichment of a gene-signature in individual cells, we computed a signature score the AddModuleScore_UCell function from the UCell package^80^. A list of genes from each biological signature was provided as an input. Higher scores were given to cells that consistently showed higher expression of genes in the marker list relative to a randomly selected set of background genes. To derive a per-patient signature score, the mean signature score of all cells in the relevant compartment was computed. Wherever necessary, the P values were adjusted using the Benjamini-Hochberg approach.

### Cell-cell communication analysis

CellChat^81^ (v2.1.0) was used to identify possible cell-cell interactions across the bone marrow microenvironment. The normalized, batch corrected count matrix (See: Batch-corrected count matrices) was used as the input to CellChat. Doublet clusters were excluded from this analysis. For downstream analysis, some clusters representing biologically similar subtypes were aggregated. A table displaying the mapping between the original clusters and their CellChat clusters is provided in Supplemental Table 6.

### Gene regulatory network analysis

The gene regulatory networks (GRNs) for selected clusters within our dataset were estimated using pySCENIC^82^, an implementation of SCENIC (Single-Cell Regulatory Network Inference and Clustering). The analysis was focused on selected clusters of interest from the myeloid compartment (CD14+Mono_IFN, CD14+Mono pro-in-lam, Macro/Mono, GMP). The batch corrected count matrix served as the input to run GRNBoost2^83^ and generate co-expression modules. GRNs were further inferred using the hg38_refseq-r80 (mc_v10_clust) motif database, hgnc motif annotation (v9) and and pySCENIC’s default settings. Due to the stochastic nature of the GRNBoost2 algorithm, slightly varying regulons are detected in each run. Hence, high confidence regulons were filtered out if they were present in >80% of runs, while their target genes were considered if they were detected in >90% of runs. Using AUCell from pySCENIC, each cell was assigned a gene signature score (AUC) indicating the degree of transcription factor activity. The AUC values were normalized across each regulon, and their mean was calculated for each cluster to identify regulons that were strongly associated with a specific cluster. AUC values for each cell in the clusters of interest were averaged to get a per patient per regulon score. The cut-point algorithm was used for grouping samples into ‘high’ and ‘low’ regulon activity categories^84^. Survival analysis was performed using the Kaplan-Meier method and Cox proportional hazards regression model on the ‘high’ and ‘low’ activity variables. As AUC values were derived from batch-corrected count matrices, shipment batch was not adjusted for in the Cox model.

## Author Contributions

The following authors helped conceived the manuscript: WP, EGK, CA, MH, STO, MPR, NK, IAC, ZC, SL, SSB, RV, GM, LD, SG, MB.

The following authors performed data analysis:

WP, LY, EGK, YPJ, DK, MEM, SN, SS, JS, DDS, MAF, JDVS, IC, IFDS, YL,, IAC, ZC, AL, SKS, SSB, SG, MB.

The following authors contributed to cell annotation:

WP, LY, EGK, YPJ, DK, CA, MEM, SS, YM, AC, IAC, SL, SSB, MB.

The following authors provided domain expertise:

WP, EGK, YPJ, DK, CA, MEM, MH, SS, JS, GC, MBa, NP, IS, RA, EL, BET, AC, MPR, YL, SD, NK, IAC, ZC, AL, JR, SKS, SL, SK, SSB, TK, DA, HJC, GM, LD, SG, ISV, MB.

The following authors provided clinical expertise:

EL, MAF, IC, MS, IAC, MVD, SK, TK, DA.

The following authors performed experiments:

YS, KS, MBa, RA, RC, BET, JF, EA, MS, VS, IAC, AHR, SL, MB.

The following authors interpreted the data:

WP, LY, EGK, YPJ, DK, CA, MEM, MH, SN, YS, JTW, SS, RGJ, JFS, AC, IC, YL, SD, NK,, IAC, ZC, AL, SKS, SSB, SG, MB.

The following authors contributed to the writing of the manuscript:

WP, LY, EGK, YPJ, DK, CA, MEM, MH, SN, SS, MBa, EA, YL, SD, MS, VS, NK,, IAC, SL, SK, SSB, TK, GM, MB.

The following authors provided oversight of the study:

EGK, CA, MH, NK, IAC, ZC, AL, SKS, SL, SK, SSB, RV, DA, GM, LD, SG, ISV, MB.

## Acknowledgments

SG was partially supported by the National Institutes of Health grants CA224319, DK124165, and CA196521, as well as the Multiple Myeloma Research Foundation (MMRF).

JFD: Paula C. and Rodger O. Riney Blood Cancer Research Fund, NCI R35 CA210084;

DK acknowledges support from the MMRF Fellowship Program; SN acknowledges support from MMRF;

MPR was supported by NCI R50CA211466;

JFD was supported by Paula C. and Rodger O. Riney Blood Cancer Research Fund, NCI R35 CA210084; TK was supported by grants 5K12CA090628;

RV was supported by Paula C. and Rodger O. Riney Blood Cancer Research Fund;

DA acknowledges support from MMRF

DL Paula C. and Rodger O. Riney Blood Cancer Research Fund, NCI U24CA211006, U2CCA233303, and PJ000021702;

ISV acknowledges support from MMRF. Data analyses were performed on the Ithaca High Performance Computing Cluster (HPC), Spatial Technologies Unit/Precision RNA Medicine Core (RRID:SCR_024905), and the Harvard Medical School O2 HPC Cluster.;

MB acknowledges support from the MMRF and support from Emory University

This work was supported in part through the computational and data resources and staff expertise provided by Scientific Computing and Data at the Icahn School of Medicine at Mount Sinai and supported by the Clinical and Translational Science Award (CTSA) grant UL1TR004419 from the National Center for Advancing Translational Sciences.

## Competing Interests

S.G. reports other research funding from Boehringer-Ingelheim, Bristol-Myers Squibb, Celgene, Genentech, Regeneron, and Takeda, and consulting from Taiho Pharmaceuticals, not related to this study. JFD is on the consulting/advisory committee for Rivervest, Bioline, Amphivena, Bluebird, Celegene, Incyte, NeoImuneTech, and Macrogenics and has ownership investment in Magenta and Wugen; JR declares consulting with Attivare, Parexel, Clario/Bioclinica, Imaging Endpoint, and Wolters Kluwer Health, Inc. Serves on a DSMB with Karyopharm. Grants and nonfinancial support from Celgene, BMS and Sanofi.; JR has a patent for PCT/US2021059199 pending.; SK declares Research funding for clinical trials to the institution: Abbvie, Amgen, Allogene, BMS, Carsgen, GSK, Janssen, Roche-Genentech, Takeda, Regeneron Consulting/Advisory Board participation: (with no personal payments) Abbvie, BMS, Janssen, Roche-Genentech, Takeda, Pfizer, Loxo Oncology, K36, Sanofi, ArcellX, Beigene; TK declares research funding from Novartis, Pfizer. Advisory Board: BMS DA declares grants from MMRF, CTN (NIHLBI), Celgene, Pharmacyclics and Kite Pharma. Other support from Juno, Partners TX, Karyopharm, BMS, Aviv MedTech Ltd., Takeda, Legend Bio Tech, Chugai, Caribou Biosciences, Janssen, Parexel, Sanofi, and Kowa.; DA has a patent for PCT/US2021/059199 pending.; ISV reports grants from NCI, NHLBI, NIDDK, Harvard Stem Cell Institute, and consulting for Mosaic LLC, AlphaSights, NextRNA, and Guidepoint Global outside of the submitted work; JAF is a consultant for CPRIT and Wugen on work unrelated to the manuscript. Unrelated to this work, J.A.F. has a monoclonal antibody licensed to EMD Millipore and is an inventor on patent/patent applications (WO 2019/152387, US 63/018,108) licensed to Kiadis Inc. and held/submitted by Nationwide Children’s Hospital on TGF-β resistant, expanded NK cells. Other authors declare no competing financial or non-financial interests

***\* Immune Atlas Consortium***:

Nikolaos Kalavros, Jennifer Rogers, Travis Dawson, Brian H. Lee, Geoffrey Kelly, Laura Walker, Nicolas F. Fernandez, John Leech, Jarod Morgenroth-Rebin, Krista Angeliadis, Matthew A. Wyczalkowski, Song Cao, Omar Ibrahim, Roderick Lin, Todd A. Fehniger, Andrew Houston

