## Supplemental Document 1 for "A single-cell atlas characterizes dysregulation of the bone marrow immune microenvironment associated with outcomes in multiple myeloma"

### Cell Population Annotation Dictionary

### Author List Footnotes:

#: Co-First authors

%: Co-Second authors

$: Senior and Co-corresponding authors

### Cell Population Annotation Dictionary

The immune cell atlas is composed of 1,149,344 cells, partitioned into 5 major compartments: NK and T cells (k=629,877, 54.80% of all cells), B cells and erythroblasts (k=232,056, 20.19% of all cells), Erythrocytes (k=20,300, 1.77% of all cells), Myeloid cells (k=168,874, 14.69% of all cells), Fibroblasts (k=946, 0.08% of all cells), and Plasma cells (k=97,291, 8.46% of all cells).

The following list will provide information for on the 106 clusters, including plasma cell and doublet populations. Percentages refer to the fraction of all cells collectively, including baseline and follow-up timepoints, without weighing by patient. For each cluster, the internal cluster name will be listed in the format of [Compartment].[Cluster].[Subcluster], along with the shorthand cell cell-type ID referred to in the rest of the main text. For non-doublet populations, a short description of the population is provided, along with markers. Refer to **Figure 2** and **Supplemental Figure 2** in the text for UMAPs and DotPlots for these populations.

#### NK and T cells

NK and T cells formed the largest compartment (k=629.877, 54.80% of all cells) and were divided into 32 cell types, including CD4+ and CD8+, NK T cells and NK cells. The T cell compartment was mainly characterized based on the expression of canonical T cell markers indicating distinct T cell states from naïve/central memory to early effector memory T cells that give rise to highly activated and cytotoxic effector memory T cells, NK T cells and pro-inflammatory phenotypes. The NK compartment, identified through lack of CD3 expression combined with canonical NK markers, was mainly characterized based on the expression of *CD56 (NCAM1)* canonical marker, ranging from immature CD56-high cells to mature CD56-dim cells.

**CD4 T cells:** The CD4 cell compartment (k=306,883, 55.55% of T cells) comprised 11 cell states.

**NkT.0 (CD4_Tn):** A CD4 naïve T cell population (k=144,092, 26.08% of T cells) with intermediate to low expression of activation markers *(CD44, CD69)* and strong expression of naïve/central memory-related markers (*SELL*, *CD7*, *LEF1*, *IL7R, TCF7, CCR7).*

**NkT.1.0 (CD4_Tcm_KLRB1):** A CD4 central memory T cell population (k=49,246, 8.95% of T cells) with intermediate expression of *CD69* and *CD44* activation markers. Strong expression of central memory-related markers *IL7R, TCF7* and of killer cell lectin-like receptor B1 (*KLRB1*).

**NkT.1.1 (CD4_Teff):** A CD4 effector population (k=21,296, 3.86% of T cells), in an intermediate state highly expressing activation markers (*CD44*, *CD69, DUSP),* as well as naïve/central memory markers including *IL7R*. Strong expression of *KLRB1* and *GZMK* was also observed.

**NkT.1.2 (CD4_Tcm_NFKBIA):** A CD4 central memory T cell population (k=15,462, 2.80% of T cells) with intermediate expression of *CD69* and *CD44* activation markers. Strong expression of central memory-related markers *CXCR4* and *FOXO1.* A co-stimulatory signal was detected with the high expression of *ICOS* and an NF-κB signaling through the high expression of *NFKBIA*.

**NkT.1.4 (CD4_Tem_IFN):** A CD4 effector memory T cell population (k=4,412, 0.80% of T cells), with high expression of memory markers including *IL7R, FAS, TCF7* and intermediate levels of *SELL*. Intermediate *CD44* expression indicating semi-activation. The cluster was highly characterized by an interferon-induced phenotype with the high expression of interferon genes, specifically ones related to ISG15 antiviral pathways, including *IFI6*, *IFI44*, *IFI44L*, *IFIT1*, *IFITM1*, and *IFITM2*.

**NkT.1.5 (CD4_Th):** A CD4 effector helper population (k=2,596, 0.47% of T cells), highly expressing activation markers (*CD44*, *CD69, DUSP, DUSP4),* as well as classical T helper markers including *GATA3* and *STAT1.* Intermediate to high expression of interferon genes, including *IFNG* and *IFITM2*.

**NkT.2.2 (CD4_CTL):** A CD4 cytotoxic T lymphocyte population (k= 13,731, 2.49% of T cells) following a lineage differentiation from CD4 Th1 cells. Characterized by high expression of both CD4-related and cytotoxic markers, including granzymes (*GZMA*, *GZMB*, *GZMK*, *GZMH*) and *GNLY*.

**NkT.7 (CD4_Th_LEF1):** A CD4 T cell population (k= 19,603, 3.55% of T cells) presenting an intermediate phenotype between naïve/memory and effector helper T cells. High expression of the early T cell activation marker *CD44* and of naïve and central memory-related markers (*LEF1*, *TCF7*, *CCR7*, *FOXP1, FOXO1*). Upregulation of markers related to a T helper phenotype, including *RORA, STAT3, STAT4,* and *STAT6,* as well as a co-stimulatory activity with high expression of *CD28.*

**NkT.8 (Treg):** A T regulatory cell population (k=18,469, 3.34% of T cells) strongly expressing canonical T reg markers, including *FOXP3*, *IL2RA*, *IL2RB*, and the dysfunctional markers *CTLA-4*, *TOX*, and *TIGIT*.

**NkT.10.1 (CD4_Teff_TNF):** A CD4 effector T cell population (k=5,108, 0.92% of T cells), with high expression of memory markers including *CD69* and *CD44*. The cell population exhibited an interferon-stimulated/induced pro-inflammatory phenotype by highly expressing *TNF*, *IFNG, IFI44*, *IFIT1*, *IFIT2*, *IFIT3*, and *NFKBIA*. NkT.10.0 (CD8_Teff_TNF) is a CD8+ equivalent population which displays many similar characteristics to this cluster.

**NkT.12 (CD4_Tcm_IFN):** A CD4 central memory T cell population (k=12,688, 2.30% of T cells) with naive/central memory markers, including *SELL*, *CCR7*, *FOXO1*, *CD7,* and *TCF7*. Intermediate levels of activation markers (*CD69*, *CD44*). An interferon-stimulated phenotype, primarily related to ISG15 antiviral pathways, was observed with the high expression of *IFI6*, *IFI44*, *IFI44L*, *IFIT1*, and *IFITM1.*

**<u>CD8 T cells:</u>** The CD8 cell compartment (k= 245,519, 44.45% of T cells) comprised 17 cell states.

**NkT.6 (CD8_Tn):** A CD8 naïve T cell population (k=31,682, 5.74% of T cells) with low expression of the activation markers *CD44* and *CD69* markers. Strong expression of naïve/central memory-related markers (*SELL*, *CD27*, *LEF1*, *IL7R*, and *TCF7*)

**NkT.1.3 (CD8_Tcm):** A CD8 central memory T-cell population (k=5,896, 1.07% of T cells) following a lineage differentiation from CD8 naïve cells (CD8_Tn). Low expression of naïve T cell markers (*TCF7*, *SELL*, *CD27*, *CD28*, *CCR7*) along with memory markers (*TRADD*, *IL7R*, *TIMP1*). Lack of activation (*CD69*, CD44), cytotoxicity or chemokine production related markers.

**NkT.2.0 (CD8_Teff):** A CD8 cytotoxic effector T cell population (k= 50,293, 9.10% of T cells). Strong expression of cytotoxic markers including (*GZMH*, *GNLY*, *PRF1*, *FGFBP2*, *NKG7*) as well as *IFNG.* Lack of *CD27* and *CD28*, indicating an endpoint in CD8 T cell development[1]. No footprints of exhaustion were observed.

**NkT.2.1 (CD8_Teff_HLA):** A CD8 transitioning-effector T cell population (k=20,614, 3.73% of T cells). following a lineage differentiation from GZMK+ Central Memory cells (CD8_Tcm_ GZMK) to GZMB+ Effector cells (CD8_Teff). High expression of specific granzymes and activation markers including, *GZMA*, *GZMM*, *GZMH, GZMK, CTSW* and *KLRK1.* Certain cytotoxicity markers such as *FGFBP2, GNLY,* and *PRF1,* are present, but lower relative to other CD8+ Cytotoxic populations. High expression of *IFNG* along with MHC-I and MHC-II class markers (*CD74*, *HLA-DRA*, *HLA-DRB1*, *HLA-DPB1*), previously discussed as late activation markers in CD8 T cells[2]. This population does not appear to have a direct mouse equivalent[3]. Exhaustion markers, including *LAG3* and *TIGIT* are also expressed.

**NkT.2.3 (CD8_Teff_b):** A CD8 effector T cell population (k=1,245, 0.23% of T cells). Strong expression of cytotoxicity markers (*GZMB*, *GZMH*, *GNLY*, *PRF1*, *FGFBP2*, *NKG7*). Intermediate expression of erythroid marker genes in the background (*HBA*, *HBB*). Population is not patient nor site specific.

**NkT.2.4 (CD8_Teff_c):** A CD8 effector T cell population (k=985, 0.18% of T cells) with intermediate to high levels of cytotoxic genes (*GZMB*, *GZMH*, *GNLY*, *PRF1*, *FGFBP2*, *NKG7*). Patient specific cytotoxic cluster, abundant in plasma cell immunoglobulins (*IGLC1*).

**NkT.3.0 (CD8_Tem):** A CD8 effector memory T cell population (k=36,489, 6.61% of T cells) with a pro-inflammatory phenotype, highly expressing early activation markers (*CD69*, *CD44*), the cytotoxic marker *GZMK* and multiple chemokine and cytokine related genes (*CCL3*, *CCL4*, *CMC1*, *XCL1*, *XCL2*). This cluster appears on a separate trajectory from the other cytotoxic populations, branching off from the CD8 Central Memory populations. Expresses some exhaustion markers, such as TIGIT.

**NkT.3.1 (CD8_Tcm_GZMK):** A CD8 central Memory T cell population (k=26,878, 4.87% of T cells) with high expression of *GZMK* cytotoxic marker. Low expression of naïve markers (*TCF7*, *CD27*, *CD28*), and upregulation of memory markers (*TRADD*, *IL7R*, *TIMP1*). Low expression of activation (*CD69* and *CD44* low), and lack of expression on cytokine/chemokine, and cytotoxicity-related markers. The cell population presented as a branch point between the CD8 naïve T cells and the cytotoxic/GZMB+, the activated GZMK+, and the MAIT lineage.

**NkT.3.2 (CD8_Tem_NFKB)**: A CD8 Effector Memory T cell population (k=10,944, 1.98% of T cells) highly expressing *NFKB* related genes. High expression of early activation markers (*CD69*, *CD44*) and the cytotoxic marker *GZMK*. High expression of markers related to NFKB signaling, including *NFKB1*, *REL*, *NR4A2*, and *TNFAIP3.*

**NkT.5.0 (CD8_T_adp):** A CMV Adaptive, NK-Like, CD8 T Cell Population (k=17,630, 3.19% of T cells). This is a cytotoxic CD8+ population, with high expression of cytotoxicity markers (*GZMB, GZMH, GNLY, PRF1, FGFBP2, NKG7)*. High expression of *FCGR3A* (*CD16*), low expression of NK receptor *NCR3*, the inhibitory NK receptor *KLRC2*, and the regulatory T cell marker *IKZF2* (Helios). High expression of the alpha-beta TCRs, *TRAC* and *TRBC1*, and the gamma-delta TCRs, *TRDC* and *TRGC2*. Appears to be related to previously described 'CMV-Adaptive' $CD8^+$ populations[4]. Similar profiles to the CMV Adaptive NK cells (NK_adp).

**NkT.5.2 (CD8_T_adp_b):** A patient specific CMV Adaptive, NK-Like, CD8 T Cell Population (k=366, 0.07% of T cells). Similar expression profiles to CD8_T_adp, including high expression of cytotoxicity molecules, and expression of the inhibitory NK receptor *KLRC2*. High expression of plasma cell immunoglobulins *IGLC2* and *IGLC3*.

**NkT.10.0 (CD8_Teff_TNF):** A TNF+ CD8 T cell population, with markers related to NFKB pathways (k=10,394, 1.98% of T cells). Intermediate expression of cytotoxicity markers (*GZMB*, *GZMH*, *GNLY*), with high expression of *TNF* and *IFNG*. High expression of activation markers (*CD69*, *CD44*) and markers related to NFKB activation (*NFKB1*, *REL*, *NR4A2*, *TNFAIP3*). Enriched in multiple interferon-inducible markers, including *IFIT2* and *IFIT3*. NkT.10.1 (CD4_Teff_TNF) is the CD4+ equivalent population and shows many similar characteristics to this cluster.

**NkT.11 (CD8_T_Apoptotic):** A stressed, apoptotic T cell population, with enriched expression of Mitochondrial markers relative to the rest of the T cell compartment (k=14,477, 2.62% of T cells).

**NkT.13 (MAIT):** Mucosal Associated Invariant T cells (k=11,610, 2.10% of T cells), highly expressing canonical markers including *NCR3*, *SLC4A10*, *ZBTB16*, and *KLRB1*. Cluster also appears to contain a subpopulation of double-negative gamma-delta T cells, defined by expression of *TRGC2* and *TRDC*.

**NkT.14 (CD8_Tem_IFN):** A CD8 T effector memory cell population (k=4,729, 0.86% of T cells) highly expressing cytotoxic markers (*GZMA*, *GZMH*, *GZMK*, *GZMM*, *NKG7*) and interferon related genes (*IFI16*, *IF35*, *IFI44*, *IFI44L*, *IFI6*, *IFIT1*, *IFIT5*, *IFITM1*). High expression of some exhaustion markers (*LAG3*, *TIGIT*, *BATF*).

**NkT.15 (Dbl11**)**:** Doublet Population (k=806, 0.15% of T cells).

**NkT.16 (Dbl12**)**:** Doublet Population (k=481, 0.09% of T cells).

<u>**Natural Killer (NK) Cells:**</u> The NK cell compartment (k=77,475, 6.74% of all cells) is comprised of four different populations. NK cells were originally found in the NK/T compartment, and were distinguished by negative expression of T cell lineage markers ($CD3^-$, $CD8A^-$, $CD4^-$), along with

positive expression of canonical NK markers, including *NCAM1* (*CD56*), *CD247*, and *NCR3*. The cell populations were divided into CD56-bright and CD56-dim.

**NkT.4 (NK_CD56dim):** CD56 Dim NK cell population (k=45,913, 59.26% of NK cells). This is a standard cytotoxic CD56 dim NK population. The cluster presented low *NCAM1* expression and high levels of *FCGR3A* (*CD16*) and *GZMB*. A highly cytotoxic NK cluster with high expression of cytotoxic markers such as *GNLY*, *PRF1*, and *FGFBP2.* The population upregulated the activating receptors *KLRB1* and *KLRF1*. Distinguished from other CD56 Dim clusters by the high expression of *FCER1G* and *SH2DB1*[5,6].

**NkT.5.1 (NK_adp): A** CMV Adaptive, CD56-Dim NK cell population (k=13,752, 17.75% of NK cells). High expression of cytotoxicity markers such as *GNLY, PRF1, NKG7, FGFBP2*. This cell population highly expressed markers associated with an 'Adaptive NK' phenotype, downregulating the *KLRB1* and *KLRF1* receptors, lacking the *FCER1G* expression, and upregulating the inhibitor NK receptors *KLRC2* and *KLRC3*[7,8].

**NkT.9.0 (NK_CD56bright):** CD56 Bright NK cell population (k=10,566, 13.64% of NK cells). This is an immature, CD56 bright NK population, distinguished by high expression of *NCAM1*, negative expression of *FCGR3A* (*CD16*), and expression of *GZMK* instead of *GZMB*. The population has lower expression of cytotoxicity molecules (*PRF1*, *NKG7*), and higher expression of chemokines (*CMC1*, *XCL1*, *XCL2*). Cluster is distinguished from other CD56 bright populations by the expression of *TCF7* and *SELL*[5,6].

**NkT.9.1 (NK_resident):** CD56 Bright, Bone Marrow Resident NK cell population (k=7,244, 9.35% of NK cells). This is a variant of a CD56 bright NK population, distinguished by the expression of *CD69*, *CD160*, *TIGIT*, and *IKZF3*, with no expression of naïve markers *TCF7* and *SELL.* This cluster is negative for *FCGR3A* (CD16) and *GZMB* and has a high expression of *GZMK*. Low expression of markers related to cytotoxicity (*PRF1, NKG7)* with a high pro-inflammatory activity by highly expressing chemokine genes such as *CMC1, XCL1, XCL2, CCL3, CCL4*[5,9].

**<u>B cells, Erythroblasts, and Progenitors</u>**

The B-Ery cell compartment (k=232,056, 20.19% of all cells) comprised 19 different cell populations including B cells (k=131,021, 11.40% of all cells) and erythroblasts (k=81,977, 7.13% of all cells). Three of them were found enriched to doublets and excluded from the downstream analysis. B cells were characterized by lineage markers including *CD19*, *CD20, CD79a*, and *CD79b*, while erythroblasts were labeled based on *S100A6*, *PRDX2*, and *BLVRB* expression.

**BEry.7 (HSC):** Hematopoetic Stem Cells (HSC, k=9,507, 0.83% of all cells). Lacking lineage B cell markers and highly expressing canonical HSC markers, including *CD34* and *AVP*.

**BEry.11 (pDCs):** Plasmacytoid Dentric cells (pDCs) (k=6,894, 0.60% of all cells). High expression of canonical pDC markers, including *CLEC4A*, neuropilin-1 (*NRP1*), IL-3R alpha (*IL3RA*), and *IRF8*. High expression of *GZMB* was also observed.

**BEry.5 (Dbl1):** Doublet (k=15615, 1.36% of all cells).

**BEry.8 (Dbl2):** Doublet (k=9178, 0.80% of all cells).

**BEry.13 (Dbl3):** Doublet (k=5046, 0.44% of all cells).

**BEry.14 (Dbl4):** Doublet (k=4427, 0.39%% of all cells).

**<u>B Cells:</u>** B cells (k=112,370, 9.78% of all cells) were split into eight different cell populations spanning the B cell maturation trajectory from B cell progenitors to memory B cells.

**BEry.1 (B_naive):** A naïve B cell population (k=41,610, 37.03% of B cells) highly expressing IgM *(IGHM)* and IgD *(IGHD)* and lacking *CD27* expression, indicative of the naïve phenotype. An elevated *CD79a* expression relative to *CD79b* was also observed, congruent with B cell maturation, in addition to markers associated with B cell co-stimulation and activation, including *CD83, IL4R*, *FCER2* (CD23), and *TNFRSF13C* (*BAFF-R*). Together these markers indicate that this population is a mature naïve B cell population recirculating from the spleen and undergoing stimulation towards an activated state.

**BEry.2 (B_imm):** An immature B cell population (k=18,452, 16.42% of B cells) highly expressing *IgM* (*IGHM*), in addition to traditional immature B cell markers, including *CD10* (*MME*), *CD24*, and *CD38* and *l*owly expressing *VPREB1* and *IGLL1* pre-B cell markers.

**BEry.3 (B_mem):** Switched Memory B cells (k=17,598, 15.66% of B cells) with elevated *CD79a* expression relative to *CD79b*, in addition to *CD83*, *IL4R*, and *TNFRSF13C* (*BAFF-R*), indicating an activated circulating mature cell state. High expression of *CD27* and *IgA (IGHA2)* and low expression of *IgM* (*IGHM*) and *IgD* (*IGHD*) indicating a switched memory phenotype.

**BEry.6 (B_trns):** Transitional B cells (k=9,849, 8.76% of B cells). In contrast to the immature B cell population, this population presented no expression of pre-B markers and upregulated IgM (*IGHM*) and IgD (*IGHD*). Decreasing expression of immature B cells markers, including *CD24* and *CD38*, as well as increasing expression of mature B markers *PTPRC* and *MS4A1* (CD20).

**BEry.9 (B_Unswitched_mem):** Unswitched Memory B cells (k=9,007, 8.02% of B cells) with elevated *CD79A* expression relative to *CD79B*, in addition to *CD83*, *IL4R*, and *TNFRSF13C* (BAFF-R). High expression of *CD27,* IgM (*IGHM*) and IgD (*IGHD*) indicating an unswitched memory phenotype.

**BEry.10 (B_pro):** Pro-B cells (k=7553, 6.72% of B cells) with low expression of *CD34* and *CD19*, recombination markers *RAG1*, *RAG2*, and *DNTT*, and pre-BCR markers *VPREB1* and *IGLL1*. Furthermore, the position of this cluster on the B cell trajectory supports its relative labelling as a pro-B cell cluster.

**BEry.12 (B_preSm):** Large pre-B cells (k=6246, 5.56% of B cells). Large pre-B cells are actively engaged in proliferation before entering V-J recombination as small pre-B cells. Upregulation of proliferative markers including *MKi67*, *TOP2A*, and *CENPF* and decreased expression of VDJ recombination genes, including *RAG1*, *RAG2*, and *DNTT.* High expression of B cell differentiation factor *XBP1*, as well as immature B cell markers, including CD10 (*MME*), *CD24*, and *CD38*.

**BEry.16 (B_preLg):** Small pre-B cells (k=2055, 1.83% of B cells). Small pre-B cells undergoing V-J recombination show decreased proliferation markers relative to large pre-B cells, as well as increased recombination markers, including *RAG1*, *RAG2*, and *DNTT*. High expression of immature B cell markers, including *CD10* (*MME*), *CD24*, and *CD38*.

**Erythroblasts:** Immature erythroid lineage populations (k=69,019, 6.05% of all cells) were split into five different populations, including megakaryocytes, erythroblasts, and a small mast cell population.

**BEry.0 (EB):** Erythroblast cell population (k=49,034, 59.81% of erythroblasts). High expression of canonical markers of erythrocytes, including *HBB*, *HBA1*, and *HBD.*

**BEry.4 (EB_MKi67_1):** Erythroblast cells with proliferative capacity (k=16,832, 20.53% of erythroblasts). High expression of canonical markers including, *HBB*, *HBA1*, and *HBD,* in addition to the proliferation markers *MKi67* and *TOP2A*.

**BEry.15.0 (MegaK):** Megakaryocytes (k=2222, 0.19% of all cells) with high expression of *CD41* (*ITGA2B*), *PF4*, and *PPBP*.

**BEry.15.1 (MastC):** Mast cells (k=435, 0.04% of all cells) with high expression of the canonical marker *c*-Kit *(KIT),* and intermediate expression of *CD34*, *FCER1A* and mast cell transcription factor, *MTIF*.

**BEry.17 (EB_MKi67_2):** Erythrocytes with high proliferative capacity (k=496, 0.61% of erythroblasts). High expression of canonical erythrocyte markers including *HBB*, *HBA1*, and *HBD* and proliferation markers *MKi67* and *TOP2A.*

**Myeloid cells**

The Myeloid compartment (k=168,874, 14.69% of all cells) comprised 18 different cell populations. Three of them were found enriched to doublets and excluded from the downstream analysis. Myeloid cells including monocytes, macrophage, neutrophil, pDCs, and Dendritic cells (DCs) were characterized by their canonical markers, *CD68*, *CD163*, CSF3R*, LAMP3, CLEC9/10A* respectively.

**Myeloid.0 (CD14+Mono_S100):** A CD14+ Monocyte population (k=31,745, 18.80% of myeloid cells). High expression of S100A family genes (*S100A8, S100A9, S100A12)* and of MHC-I and MHC-II class markers, including (*HLA-DRA*, *HLA-DRB1*, *HLA-DPB1, HLA-DPA1, HLA-DQB1, HLA-DMA*)

**Myeloid.1 (CD14+Mono_CTSS):** A CD14+ Monocyte population (k=23,190, 13.73% of myeloid cells). High expression of *CTSS* gene.

**Myeloid.2 (CD14+Mono_IFN):** Interferon stimulated CD14+ Monocyte population (k=16,001, 9.48% of myeloid cells). High expression of interferon-related genes, including *ISG15, IFI44, IFI6, IFI30, IFITM3, IFI44L, IFITM2, IFI16, IFI44, IFIT3, IFIT2, IFIT1, IFIH1, IFITM1, IFIT5*.

**Myeloid.3 (CD14+Mono_hypo):** A Hypoxic CD14+ Monocyte population (k=14,162, 8.39% of myeloid cells). This cluster exhibits substantial hypoxia, with a high expression of *HIF1A* and *MAP3K8* genes.

**Myeloid.4 (CD16+Mono):** A CD16+ non-classical Monocyte population (k=14,032, 8.31% of myeloid cells) characterized by the high *CD16* expression.

**Myeloid.5 (GMP):** A Granulocyte-monocyte progenitor cells (GMP) (k=12,293, 7.28% of myeloid cells) with high expression of the canonical marker *MPO*.

**Myeloid.6 (Granulocyte):** Granulocytes (k=11,034, 6.53% of myeloid cells). High expression of myeloid lineage related markers (*LYZ*) as well as markers for GMP (such as *MPO*), indicating a granulocytic phenotype.

**Myeloid.7 (Dbl8):** Doublet (k=10,002, 5.92% of myeloid cells).

**Myeloid.8 (CD14+Mono pro-inflam):** A pro-inflammatory CD14+ monocyte population (k=9,539, 5.65% of myeloid cells) characterized by the strong expression of pro-inflammatory markers, such as *IL1B, CXCL12, NLRP3, CCL3* and *CCL4*.

**Myeloid.9 (Dbl9):** Doublet (k=7,823, 4.63% of myeloid cells).

**Myeloid.10 (Neutrophil_RPS/RPL):** Neutrophils with high expression of ribosomal protein genes (k=5,089, 3.01% of myeloid cells). High expression of *MPO* and *AZU1* genes, characteristic granule proteins of neutrophils. The high expression of RPS/RPLs, suggested the possibility of a lower quality cluster.

**Myeloid.11.0 (cDC2):** Conventional dendritic cell 2 (cDC2) cells (k=4,468, 2.65% of myeloid cells). High expression of *CD1C*, *FCER1A*, and *CLEC10A*, corresponding to a cDC2 phenotype.

**Myeloid.11.1 (cDC1):** Conventional dendritic cell 1 (cDC1) cells (k=304, 0.18% of myeloid cells). High expression of *CLEC9A*, *BATF3* and *XCR1*, representing a cDC1 phenotype.

**Myeloid.11.2 (pDCs):** Plasmacytoid dendritic cells (pDC) (k=27, 0.02% of myeloid cells) with high expression of *LAMP3* and *FSCN1.*

**Myeloid.12 (Macro/Mono):** A Macrophage/Monocyte population (k=4,433, 2.63% of myeloid cells). High expression of *CD14* and *CD163*, corresponding to a transitional state between a monocyte and a macrophage phenotype.

**Myeloid.13 (Neutrohpil_ARG1): Neutrophils** (k=1,904, 1.13% of myeloid cells) highly expressing *ARG1*. Strong levels of canonical neutrophil markers, including *CSF3R*.

**Myeloid.14 (Dbl10):** Doublet (k=1,788, 1.06% of myeloid cells).

**Myeloid.15 (M2_Macro):** M2 Macrophages (k=1,040, 0.62% of myeloid cells). Elevated expression of *CD163, CD206, CD209, TGFB1, IL10,* and *IL18* genes, representing M2 phagocytic phenotype, as well as high expression of *C1QA, C1QB,* and *APOE* genes*.*

**Fibroblasts**

**Full.23 (Fibroblasts):** Fibroblasts (k=946, 0.08% of all cells). This is a small population of cells which was isolated from all other clusters. Select top markers include *CXCL12, COL1A2, VCAN, DCN*.

**Erythrocytes**

The Erythrocytes (k=20,300, 1.77% of all cells) comprised 11 cell states. All cells formed a separate component on the initial UMAP distinct from all other populations, including the erythroblasts. All strongly express erythroid markers (*HBB*, *HBA*, *HBD*). Population was subclustered and

**Ery.0:** Erythrocyte population. Top markers include *TMCC2*, *XPO7*, *TRIM58* (k=3,663, 18.04% of erythrocytes).

**Ery.1:** Erythrocyte population, mitochondrial enriched. Top markers include *MT-CO1*, *MT-CO2*, *MT-CO3*. Top non-mitochondrial markers include *SPTA1*, *SLC4A1*, *TFRC* (k=3,342, 16.46% of erythrocytes).

**Ery.2:** Erythrocyte population. Top markers include *UBB*, *HBB*, and *ARL4A* (k=3,102, 15.28% of erythrocytes).

**Ery.3:** Erythrocyte population. Top markers (Seurat) include *PRDX2*, *CA1*, *CA2* (k=2,001, 9.86% of erythrocytes).

**Ery.4:** Erythrocyte population. Top markers include *HBD*, *HBB*, and *BLVRB* (k=1,923, 9.47% of erythrocytes).

**Ery.5:** Erythrocyte population. Top markers include *NEAT1*, *UBB*, *EIF1* (k=1,565, 7.71% of erythrocytes).

**Ery.6:** Erythrocyte population, with markers related to ISG15 antiviral pathways. Population has high expression of *IFI27*, *ISG15*, *IFIT3*, *IFI6*, and multiple other interferon inducible genes (k=1,271, 6.26% of erythrocytes).

**Ery.7** (Dlb5): Doublet (k=972, 4.79% of erythrocytes).

**Ery.8:** Erythrocyte population with expression of proliferative markers (*CENPF*, *MKI67*) (k=839, 4.13% of erythrocytes).

**Ery.9**: Erythrocyte population with low number of uniquely expressed genes. Top markers include HBG1 and HBG2 (k=794, 3.91% of erythrocytes).

**Ery.10** (Dlb6): Doublet (k=696, 3.43% of erythrocytes).

**Ery.11** (Dlb7): Doublet (k=132, 0.65% of erythrocytes).

### Plasma cells

A population of plasma cells (k=97,291, 8.46% of all cells) were identified based on the expression of various markers such as *JCHAIN, MZB1, SDC1*. These cells are likely a sampling of malignant plasma cells which remained following CD138 isolation, as has been observed in other studies. InferCNV estimates a high frequency of copy number alterations, consistent with these cells being malignant (**Supplementary Figure 2**). Additionally, CD138 isolation would remove normal plasma cells as well. As expected from myeloma cells, the compartment was highly heterogeneous, and multiple patient-specific clusters were noted without per-patient batch correction. Many patient specific populations still remain. Clusters were annotated based on top markers.

**Plasma.0 (Pc_IGKC):** Plasma cell IGKC expressing population (k=24,266, 24.94% of plasma cells). Top DE markers: IGKC, RPS10, RPLP1[10].

**Plasma.1 (Pc_JUN):** Plasma cell JUN expressing population (k=15,520, 15.95% of plasma cells). Top DE markers: JUN, AC007952.4, JUND[11].

**Plasma.2 (Pc_MALAT1):** Plasma cell MALAT1 expressing population (k=11,165, 11.48% of plasma cells). Top DE markers: MALAT1, NEAT1, MT-CO2[12].

**Plasma.3 (Dbl13):** Doublet (k=7,019, 7.21% of plasma cells).

**Plasma.4 (Pc_EIF1):** Plasma cell EIF1, Vimentin expressing population (k=6,907, 7.10% of plasma cells). Top DE markers: EIF1, VIM, EVI2B[13].

**Plasma.5 (Pc_MT-CO2):** Plasma cell MT-CO2 expressing cell population (k=5,397, 5.55% of plasma cells). Top DE markers: MT-CO2, MT-CYB, MT-CO3. The presence of these markers may represent an apoptotic population.

**Plasma.6 (Pc_ATF5):** Plasma cell ATF5 expressing cell population (k=4,524, 4.65% of plasma cells). Top DE markers: ATF5, SELENOS, SEC61G.

**Plasma.7 (Pc_HBB_HMGB1):** Plasma cell HBB, HMGB1 expressing cell population (k=4,380, 4.50% of plasma cells). Top DE markers: HBB, HMGB1, HIST1H4C. The presence of HBB may correlate with erythrocytic contamination.

**Plasma.8 (Pc_HBB_S100A8):** Plasma cell HBB, S100A8 expressing cell population (k=4,112, 4.23% of plasma cells). Top DE markers: HBB, S100A8, S100A9. The presence of HBB may correlate with erythrocytic contamination.

**Plasma.9 (Dbl14):** Doublet (k=3,571, 3.67% of plasma cells).

**Plasma.10 (Dbl15):** Doublet (k=3,221, 3.31% of plasma cells).

**Plasma.11 (Dbl16):** Doublet (k=2,711, 2.79% of plasma cells).

**Plasma.12 (Pc_IFI27):** Plasma cell IFI27 expressing cell population (k=880, 0.90% of plasma cells). Top DE markers: ISG15, IFI27, IFI6[14]. The presence of these markers may represent an interferon responsive population.

**Plasma.13 (Pc_CD74):** Plasma cell CD74 expressing population (k=860, 0.88% of plasma cells). Top DE markers: CD74, HLA-DRA, RPL30. MHC-II Expression on malignant plasma cells has been observed in other studies.[15]

**Plasma.14 (Pc_PPM1K):** Patient specific plasma cell PPM1K expressing population (k=732, 0.75% of plasma cells). Top DE markers: PPM1K, RNGTT, FCRLA.

**Plasma.15 (Pc_BCMA):** Patient specific plasma cell BCMA expressing population (k=451, 0.46% of plasma cells). Top DE markers: IGHG1, IGLC2, TNFRSF17.

**Plasma.16 (Pc_NOL4):** Patient specific plasma cell NOL4 expressing population (k=399, 0.41% of plasma cells). Top DE markers: RPS5, NOL4, RPS3A.

**Plasma.17 (Pc_PUS3):** Patient specific plasma cell PUS3, TMEM60 expressing population (k=244, 0.25% of plasma cells). Top DE markers: PUS3, TMEM60.

**Plasma.18 (Pc_IGHV3):** Patient specific plasma cell IGHV3 expressing population (k=242, 0.25% of plasma cells). Top DE markers: IGHV3-11, IGLV1-44, BIRC3.

**Plasma.19 (Pc_IGKV4):** Patient specific, low quality plasma cell IGKV4 expressing population (k=185, 0.19% of plasma cells). Top DE markers: IGKV4-1, MED8, RNF34.

**Plasma.20 (Pc_MPO):** Patient specific, low quality plasma cell MPO expressing population (k=170, 0.17% of plasma cells). Top DE markers: MPO, LRRC75A, SPON2. The presence of MPO may represent neutrophilic contamination.

**Plasma.21 (Pc_IGHG3):** Patient specific plasma cell IGHG3 expressing population (k=140, 0.14% of plasma cells). Top DE markers: IGHG3, RPLP1, KRTCAP2.

**Plasma.22 (Pc_CCDC88A):** Patient specific plasma cell CCDC88A expressing population (k=104, 0.11% of plasma cells). Top DE markers: CCDC88A, JSRP1, SPAG4.

**Plasma.23 (Pc_IGHA1):** Patient specific plasma cell IGHA1 expressing population (k=91, 0.09% of plasma cells). Top DE markers: IGHA1, IGKV1-6, PLPP5.

**Cell Population Dictionary References**
