## Supplemental Figures (1-15) for "A single-cell atlas characterizes dysregulation of the bone marrow immune microenvironment associated with outcomes in multiple myeloma"

### Author List Footnotes:

#: Co-First authors

%: Co-Second authors

$: Senior and Co-corresponding authors

### **Running Title:** The Immune Atlas of Multiple Myeloma

**Keywords**: Multiple Myeloma, Bone Marrow Microenvironment, Single-Cell, Transcriptome, Inflammation, Senescence

### $ Senior and Co-corresponding authors:

**Manoj Bhasin, PhD**
Health Sciences Research Building,
Room N320
1760 Haygood drive Atlanta, GA 30322


**Ioannis Vlachos, PhD**
330 Brookline Ave, 519A, Dana Building,
BIDMC, Boston, MA 02115


**Sacha Gnjatic, PhD**
1470 Madison Avenue, Hess s5-105,
Box 1044A, New York NY 10029


**Li Ding, PhD**
4444 Forest Park Avenue
St. Louis, MO 63108


**George Mulligan, PhD**
Multiple Myeloma Research Foundation
383 Main Avenue, 5th Floor.
Norwalk, CT 06851


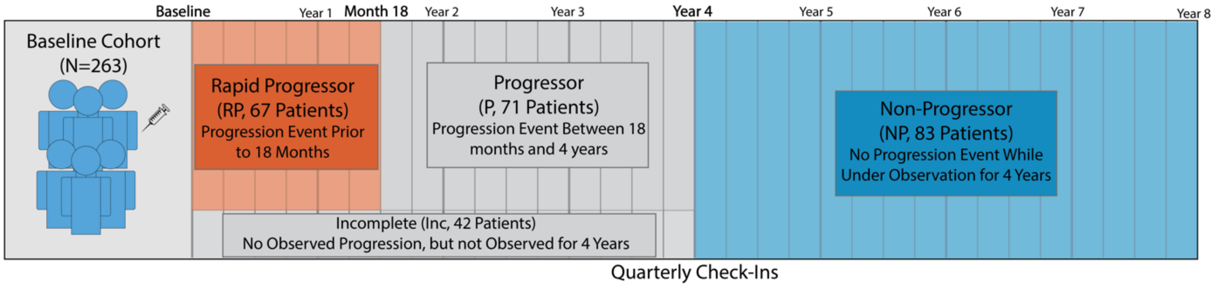


**Supplemental Figure 1: Overview of Progression Group Categorization and Study Design.** All patients in this cohort (n=263) had a sample collected at baseline at the time of multiple myeloma diagnosis prior to any treatment. Patients enrolled in the trial received their first line of therapy and had periodic follow-ups every 3 months up to eight years to evaluate drug response. At each visit, a clinical assessment was performed to classify the patient as 'SCR', 'CR', 'SD', 'Or PD'. If significant changes in clinical parameters were observed, patients may undergo bone marrow biopsy to confirm relapse or remission. For this study, patients who have a progression event within the first 18 months following therapy are classified as 'Rapid Progressors' (RP, n=67). Patients with durable remission or no observed progression for at least four years are classified as 'Non-Progressors' (NP, n=83). Patients with a progression event between 18 months and 4 years are classified as 'Progressors' (P, n=71). The patients who exited the study before four years of disease diagnosis without experiencing a progression event are classified as' Incomplete' (Inc, n=42).

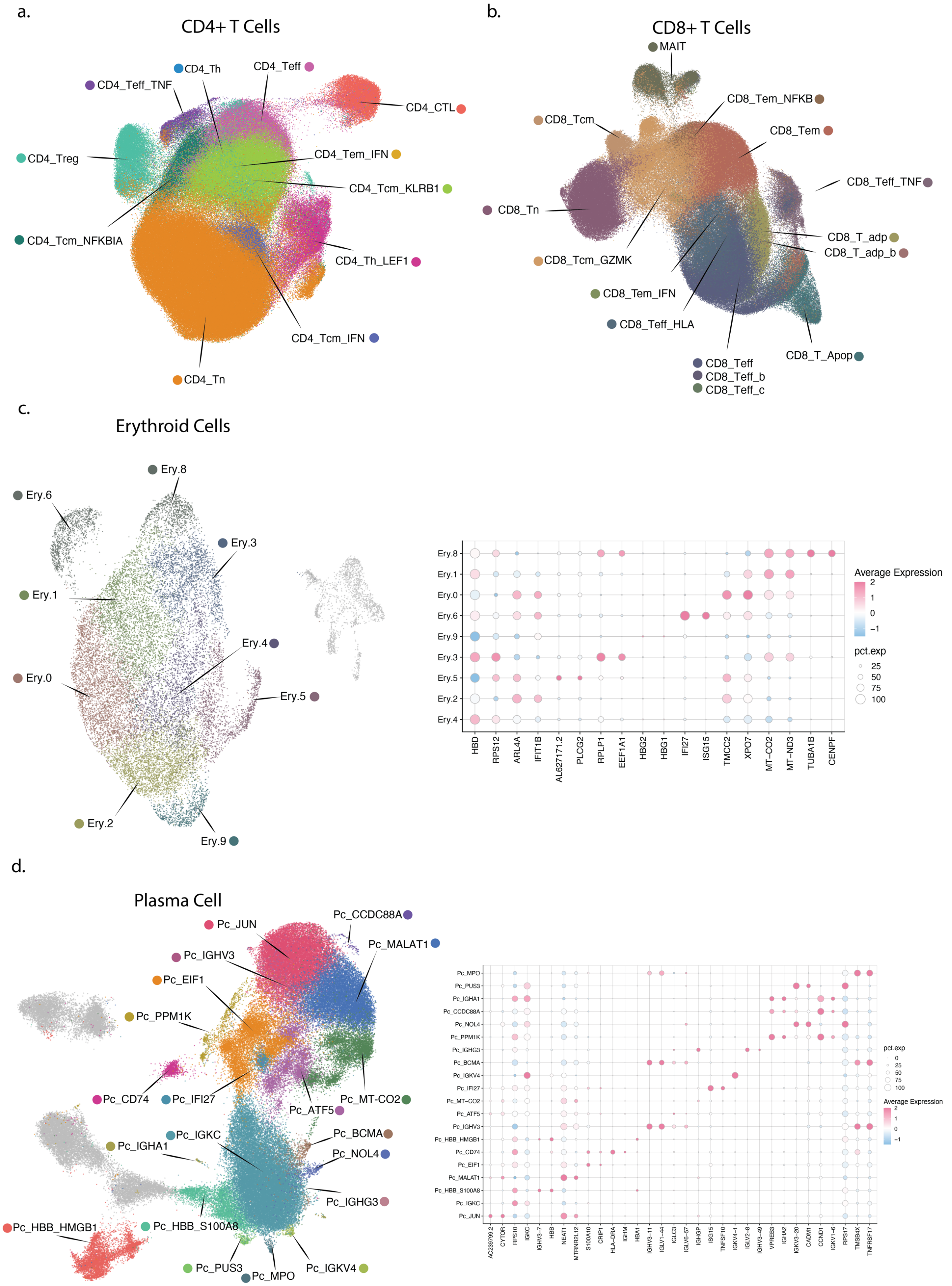
a.
CD4+ T Cells
CD4_Th
CD4_Teff
CD4_Teff_TNF
CD4_CTL
CD4_Treg
CD4_Tem_IFN
CD4_Tcm_KLRB1
CD4_Tcm_NFKBIA
CD4_Th_LEF1
CD4_Tcm_IFN
CD4_Tn
b.
CD8+ T Cells
MAIT
CD8_Tem_NFKB
CD8_Tcm
CD8_Tem
CD8_Teff_TNF
CD8_Tn
CD8_T_adp
CD8_T_adp_b
CD8_Tcm_GZMK
CD8_Tem_IFN
CD8_Teff_HLA
CD8_T_Apop
CD8_Teff
CD8_Teff_b
CD8_Teff_c
c.
Erythroid Cells
Ery.8
Ery.6
Ery.3
Ery.1
Ery.4
Ery.0
Ery.5
Ery.2
Ery.9
Average Expression
pct.exp
HBD
RPS12
ARL4A
IFIT1B
AL627171.2
PLCG2
RPLP1
EEF1A1
HBG2
HBG1
IFI27
ISG15
TMCC2
XPO7
MT−CO2
MT−ND3
TUBA1B
CENPF
d.
Plasma Cell
Pc_JUN
Pc_CCDC88A
Pc_MALAT1
Pc_IGHV3
Pc_EIF1
Pc_PPM1K
Pc_CD74
Pc_IFI27
Pc_MT-CO2
Pc_ATF5
Pc_IGKC
Pc_BCMA
Pc_IGHA1
Pc_NOL4
Pc_IGHG3
Pc_HBB_HMGB1
Pc_HBB_S100A8
Pc_PUS3
Pc_MPO
Pc_IGKV4
pct.exp
Average Expression

**Supplemental Figure 2: T cells, Plasma, and erythrocyte clusters identified in the study.** UMAPs displaying **(a.**) CD4$^+$ T cell clusters and **(b.**) CD8$^+$ T clusters. UMAPs and cluster marker Dot plots for **(c.)** Erythrocytes and **(d.)** Plasma cells. Populations identified as doublets are colored in grey. The dots in the dot plots are color-coded according to average expression levels, where red represents high expression and blue represents low expression. Additionally, the size of each dot reflects the percentage expression.

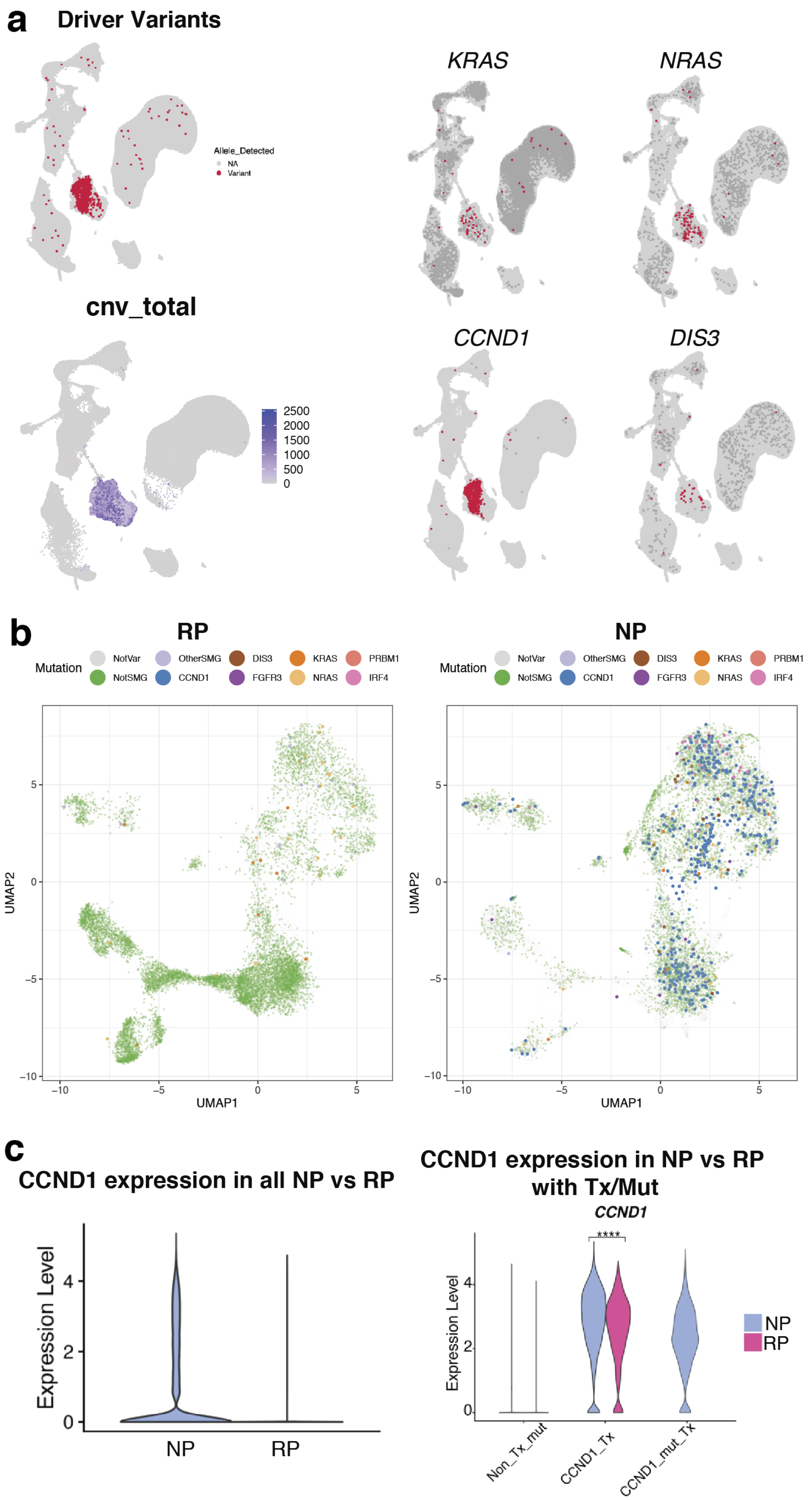
a
Driver Variants
Allele_Detected
NA
Variant
KRAS
NRAS
cnv_total
2500
2000
1500
1000
500
0
CCND1
DIS3
b
RP
NP
Mutation
NotVar
NotSMG
OtherSMG
CCND1
DIS3
FGFR3
KRAS
NRAS
PRBM1
IRF4
UMAP1
UMAP2
c
CCND1 expression in all NP vs RP
Expression Level
NP
RP
CCND1 expression in NP vs RP with Tx/Mut
CCND1
****
Non_Tx_mut
CCND1_Tx
CCND1_mut_Tx

**Supplemental Figure 3: Analysis of Plasma cells identified in the CD138$^{neg}$ fraction of BM.** **(a)** *Top:* UMAP of all cell types, as shown in Figure 2A. Cells with mutations in any of the driver genes (listed in panel b) are in red, cells with reference alleles are in dark grey, and cells with no WES data for mutation mapping are in light grey. Bottom: UMAP of plasma cells with cells colored by inferred copy number changes (red). **(b)** UMAP of cells from baseline samples of rapid progressors *(left)* or non-progressors *(right).* Cells are colored by various mutations. **(c)** *CCND1* expression of cells in the RP vs NP patients. *CCND1* expression of cells from patients with mutations (mut) and/or translocations (Tx) based on analysis of bulk sequencing genomic data.

*Supplementary Figure 4*

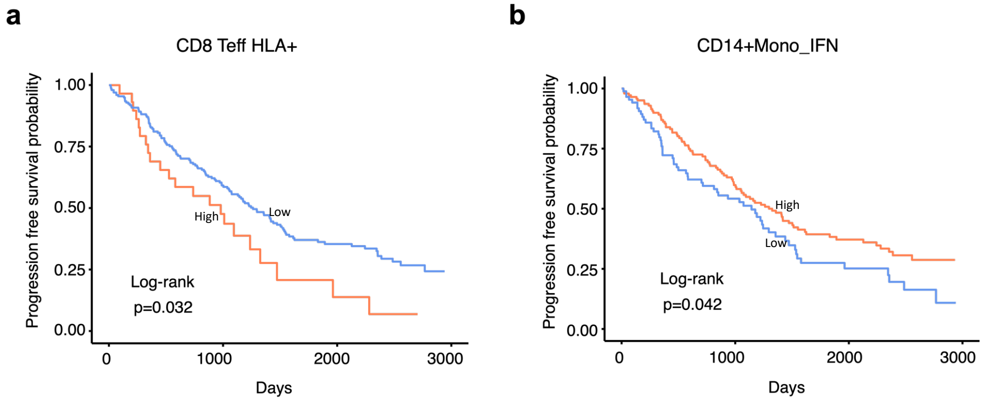


**Supplemental Figure 4: Progression-free survival curves for differentially abundant high and standard risk cell populations. (a)** Survival curve from regressing the progression-free survival on the putatively dysfunctional CD8+ Teff HLA+ cell abundances. The cell abundances were corrected for batch by taking the Pearson residuals from a Dirichlet regression model with batch as the covariate. The cut-off was determined using maximally selected rank statistics and set at the 87% quantile (201 low, 29 high). The enrichment of CD8+ Teff HLA+ is associated with significantly poor survival. **(b)** Survival curve regressing the progression-free survival on the CD14+ IFN+ Monocytes cell abundances. The cell abundances were corrected for batch by taking the Pearson residuals from a Dirichlet regression model with batch as the covariate. The cut-off was determined using maximally selected rank statistics and set at the 37% quantile (86 low, 144 high). The enrichment of CD14+ IFN+ Monocytes is associated with significantly better survival.

*Supplementary Figure 5*

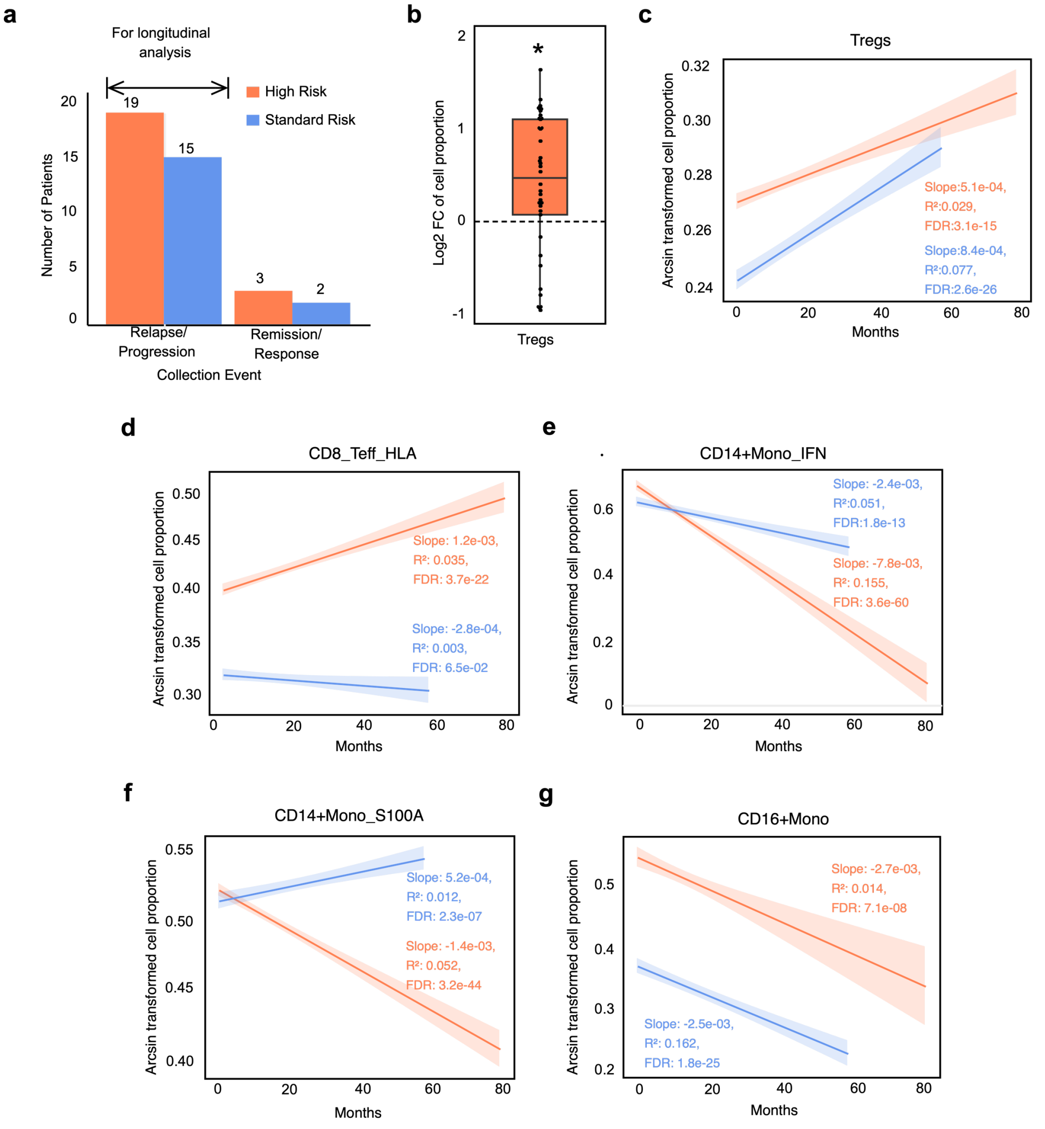


**Supplemental Figure 5: Longitudinal Analysis of relapsed patient groups with follow-up samples. (a)** Bar plots portraying the number of patients showing relapse/ progression and remission/ response classified by risk groups (i.e., HR, SR). Patients showing relapse are chosen for longitudinal analysis. **(b)** Box plots displaying the fold change (log2 scale) of T reg cell proportions between first and last time points in high-risk patients. **Significant difference is denoted with * (~~-~~p-value < 0.01; Wilcoxon paired signed rank test) (c~~C~~-g~~G~~)** Linear model representation with trend line and ± standard deviation of **(c)** Treg, **(d)** CD8_Teff_HLA, **(e)** CD14+Mono_IFN **(f)** CD14+Mono_S100A **(g~~G~~)** CD16+Mono cell proportions over time across high- and standard-risk groups. False Discovery Rate (FDR)- Benjamini Hochberg correction, $R^2$, and slope values calculated based on a linear regression model are depicted.

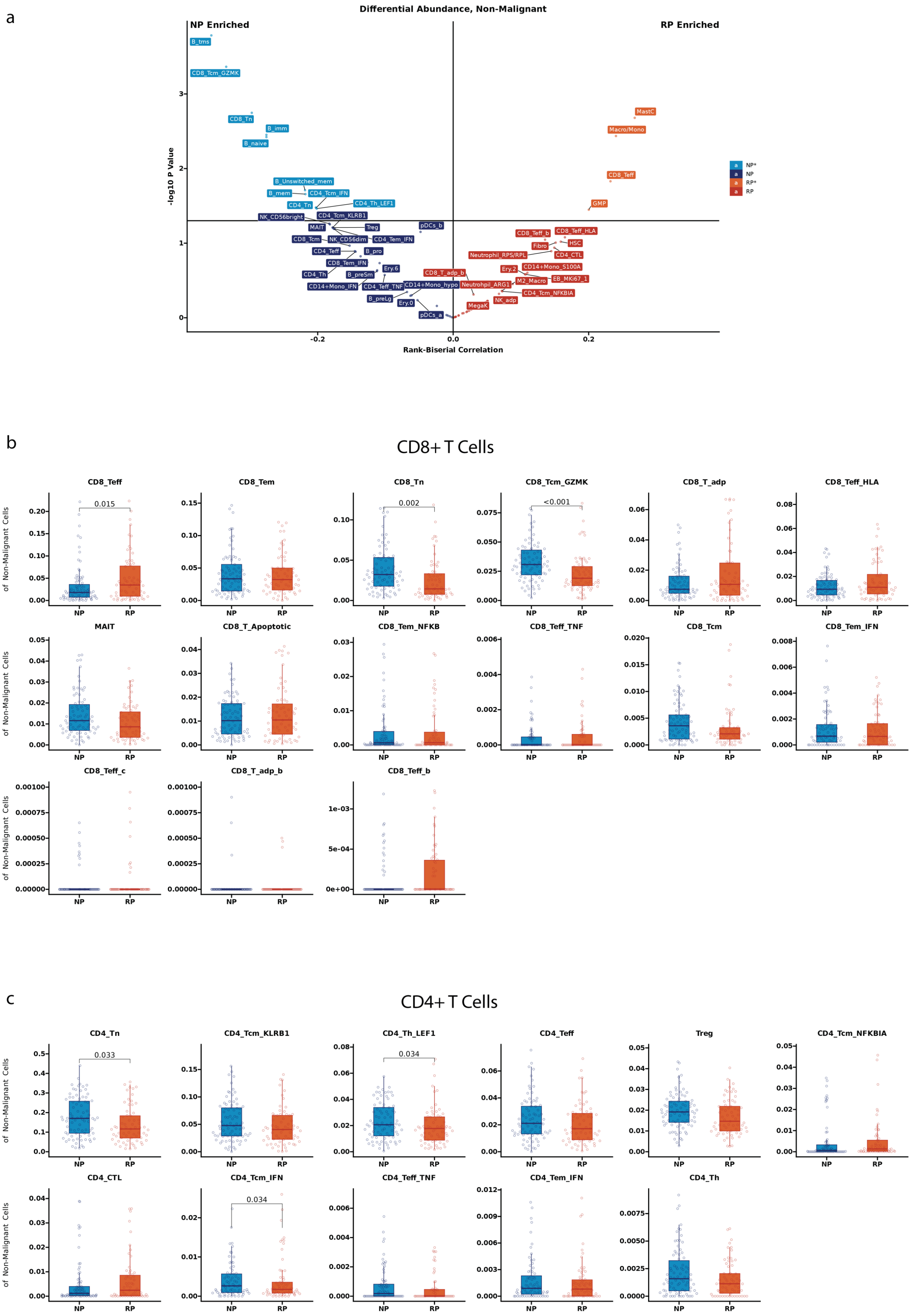
a
Differential Abundance, Non-Malignant
NP Enriched
RP Enriched
-log10 P Value
Rank-Biserial Correlation
NP*
NP
RP*
RP
B_tms
CD8_Tcm_GZMK
CD8_Tn
B_imm
B_naive
B_Unswitched_mem
B_mem
CD4_Tcm_IFN
CD4_Tn
CD4_Th_LEF1
NK_CD56bright
CD4_Tcm_KLRB1
MAIT
Treg
pDCs_b
CD8_Tcm
NK_CD56dim
CD4_Tem_IFN
CD4_Teff
B_pro
CD8_Tem_IFN
Ery.6
CD4_Th
B_preSm
CD14+Mono_IFN
CD4_Teff_TNF
CD14+Mono_hypo
B_preLg
Ery.0
pDCs_a
MastC
Macro/Mono
CD8_Teff
GMP
CD8_Teff_b
CD8_Teff_HLA
Fibro
HSC
Neutrophil_RPS/RPL
CD4_CTL
CD8_T_adp_b
Ery.2
CD14+Mono_S100A
EB_MKI67_1
Neutrohpil_ARG1
M2_Macro
CD4_Tcm_NFKBIA
MegaK
NK_adp
b
CD8+ T Cells
Proportion of Non-Malignant Cells
CD8_Teff
0.015
CD8_Tem
CD8_Tn
0.002
CD8_Tcm_GZMK
<0.001
CD8_T_adp
CD8_Teff_HLA
MAIT
CD8_T_Apoptotic
CD8_Tem_NFKB
CD8_Teff_TNF
CD8_Tcm
CD8_Tem_IFN
CD8_Teff_c
CD8_T_adp_b
CD8_Teff_b
NP
RP
c
CD4+ T Cells
CD4_Tn
0.033
CD4_Tcm_KLRB1
CD4_Th_LEF1
0.034
CD4_Teff
Treg
CD4_Tcm_NFKBIA
CD4_CTL
CD4_Tcm_IFN
0.034
CD4_Teff_TNF
CD4_Tem_IFN
CD4_Th

d

### Myeloid Cells

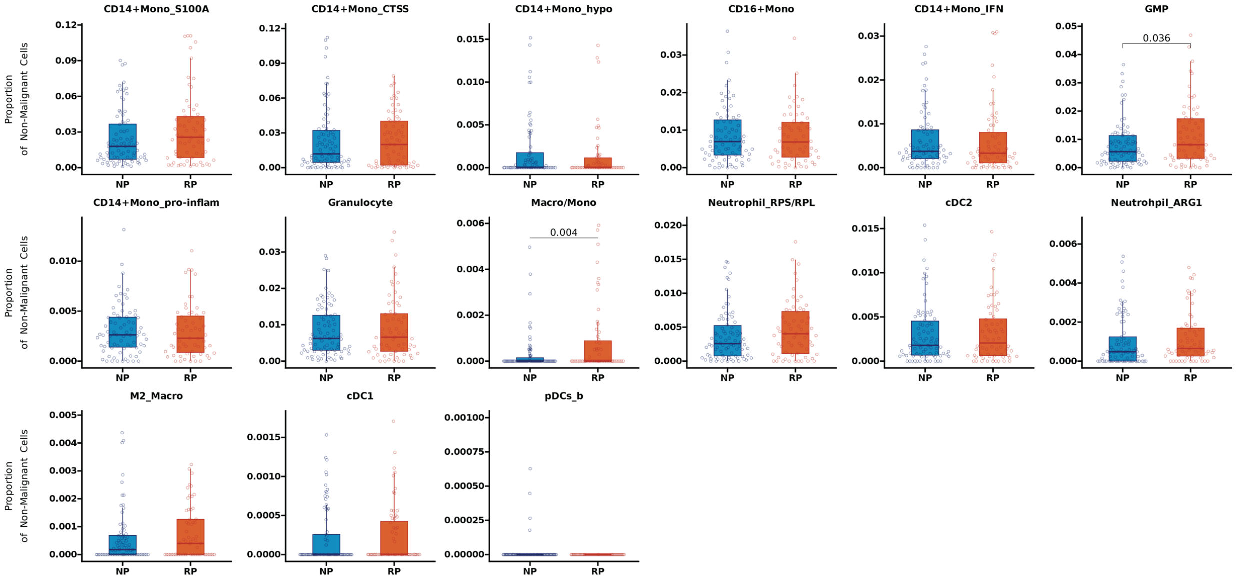


e

### B Cells

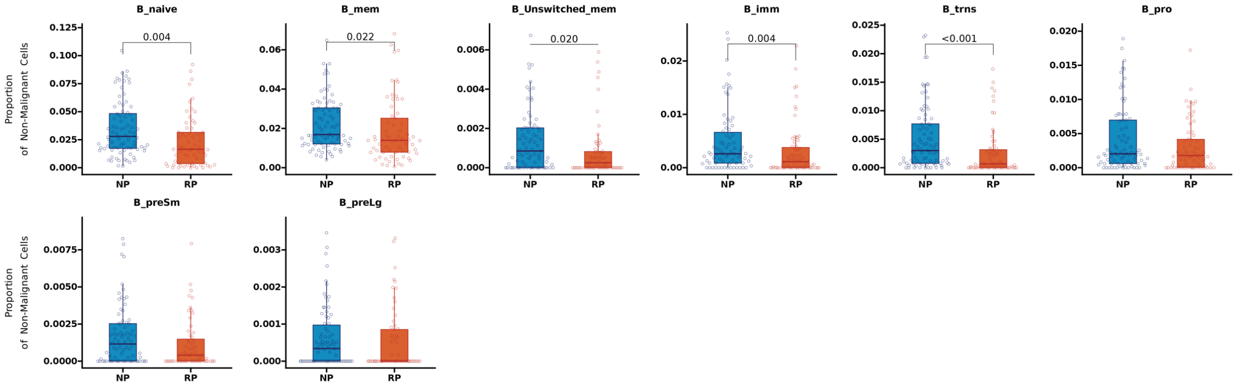


f

### Other Cells

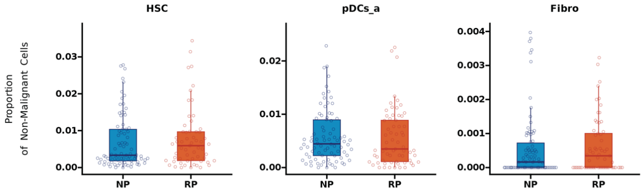

g

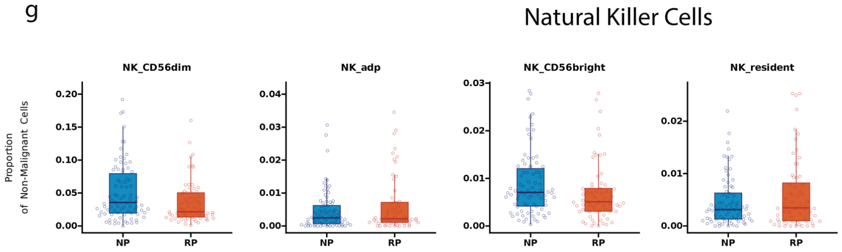


h

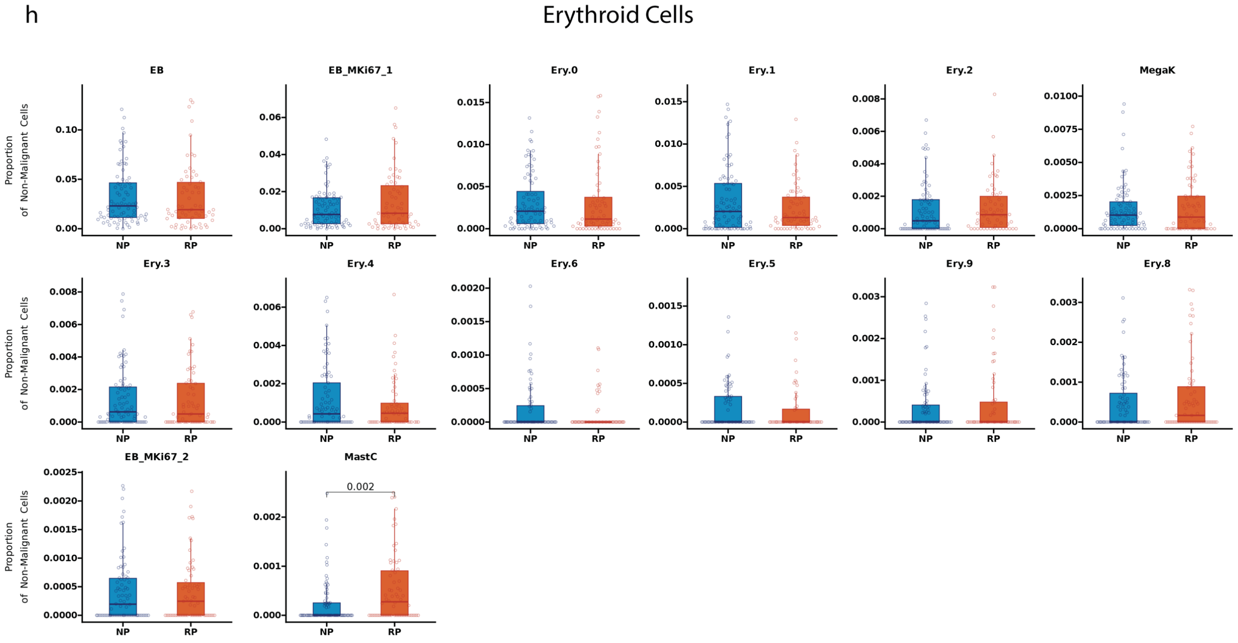


**Supplemental Figure 6: Differential abundance analysis for non-malignant cells based on progression stratification (Rapid Progressor (RP) and Non-Progressors (NP).** Differential abundance results comparing NP and RP cellular proportions for each cluster. The cellular proportions for each patient were determined based on the total non-malignant cells. **a**. Scatter plot summarizing the differential abundance results. The x-axis displays the rank-biserial correlation coefficient, while the y-axis displays the -log10 of the p-value from the Wilcoxon rank-sum test. Labels with lighter shades represent clusters with significant changes in proportions between rapid and non-progressing patients ($p < 0.05$). Box plots displaying the proportion of cells of a given cluster as a fraction of non-malignant cells, with clusters grouped by $CD8^{+}$ T cells **(b),** $CD4^{+}$ T cells (**c)**, Myeloid cells (**d)**, B Cells (**e**), Other/Independent cell populations (**f**), and Natural Killer cells (**g**), and Erythroid lineage cells (**h)**. The significance of the differences in cell proportions was calculated using the Wilcoxon rank sum test and added to panels if significant ($p < 0.05$).

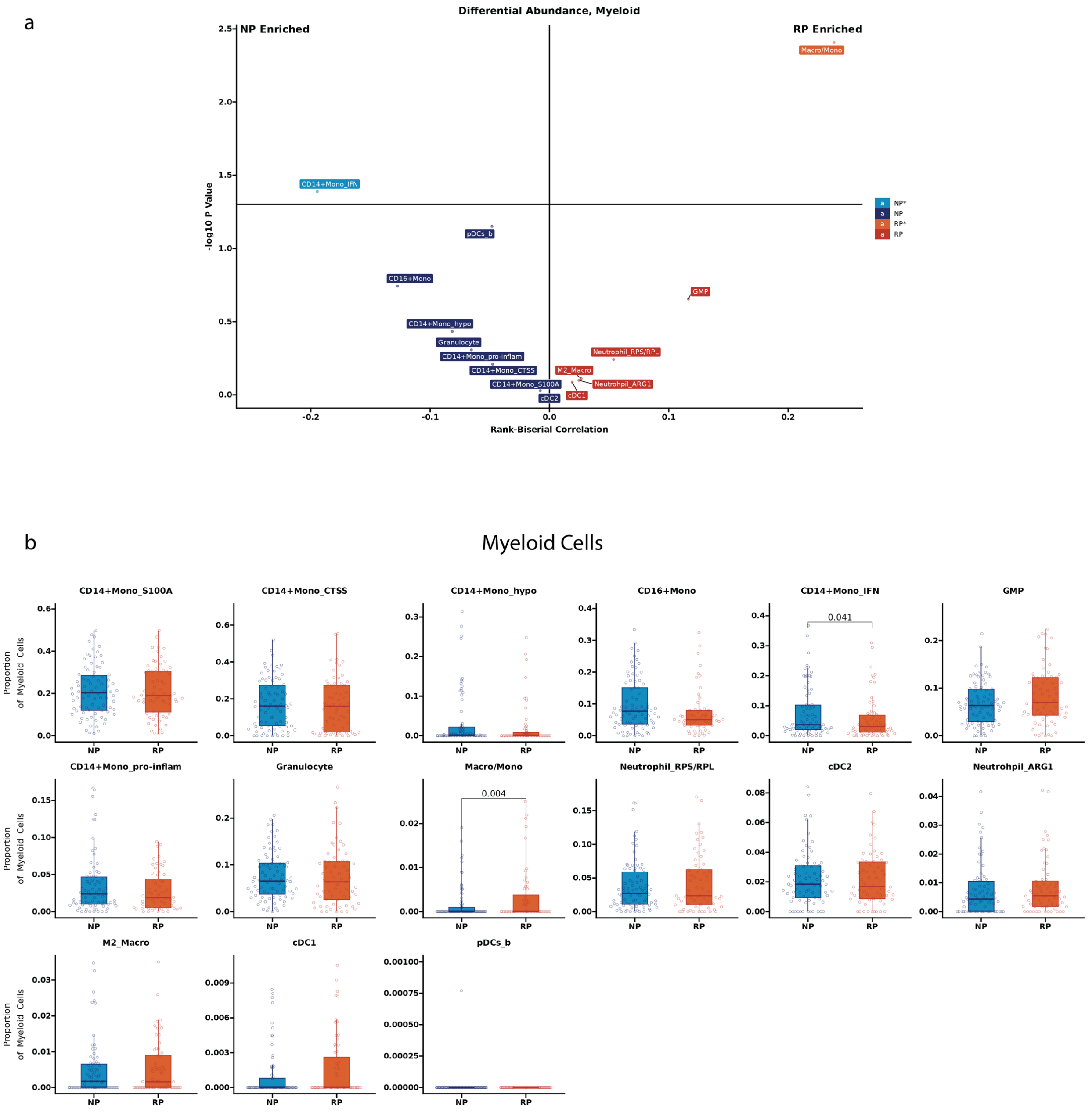


**Supplemental Figure 7: Focused Differential Abundance Analysis on the Myeloid Cell compartment. a.** Scatter plot summarizing the differential abundance results for the myeloid cell clusters by calculation proportion based on total myeloid cells in the study. The x-axis displays the rank-biserial correlation coefficient, while the y-axis displays the -log10 of the p-value of the Wilcoxon rank-sum test. Labels with lighter shades represent clusters with significant differences between rapid and non-progressors ($p < 0.05$). **b.** Box plots displaying the proportion of cells within a given cluster of myeloid cells stratified based on progression. The significance of the difference in cell proportions was calculated using the Wilcoxon rank sum test and added to the panel if significant ($p < 0.05$).

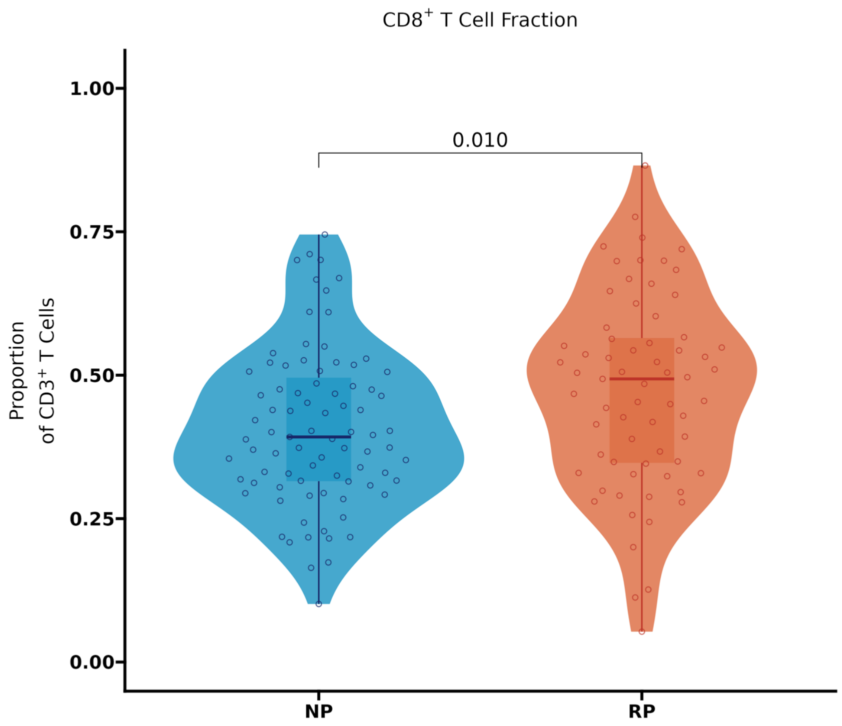


**Supplemental Figure 8**: **Comparison of CD8+ T Cell proportion of total CD3+ T cells.** Box and violin plot displaying the average patient proportion of CD8+ T cells determined based on total CD3+ T cells. Circles represent individual patients. The significance of the difference in cell proportions between rapid and non-progressors was calculated using the Wilcoxon rank sum test.

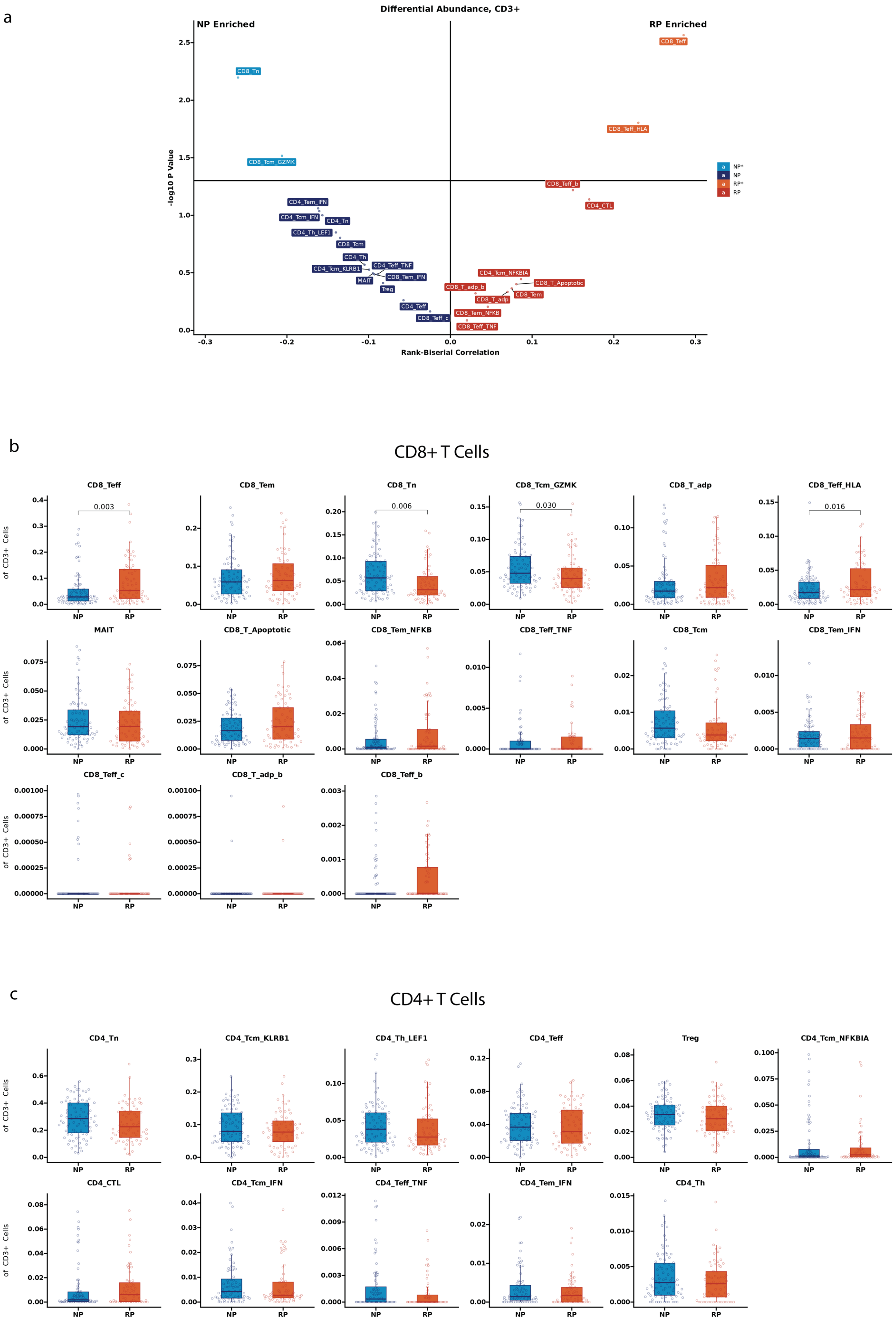

a
Differential Abundance, CD3+
NP Enriched
RP Enriched
-log10 P Value
Rank-Biserial Correlation
b
CD8+ T Cells
CD8_Teff
CD8_Tem
CD8_Tn
CD8_Tcm_GZMK
CD8_T_adp
CD8_Teff_HLA
MAIT
CD8_T_Apoptotic
CD8_Tem_NFKB
CD8_Teff_TNF
CD8_Tcm
CD8_Tem_IFN
CD8_Teff_c
CD8_T_adp_b
CD8_Teff_b
Proportion of CD3+ Cells
c
CD4+ T Cells
CD4_Tn
CD4_Tcm_KLRB1
CD4_Th_LEF1
CD4_Teff
Treg
CD4_Tcm_NFKBIA
CD4_CTL
CD4_Tcm_IFN
CD4_Teff_TNF
CD4_Tem_IFN
CD4_Th

**Supplemental Figure 9: Differential abundance analysis on the T cell compartment between patients with rapid and no progression of MM. a)** Scatter plot summarizing the differential abundance results of the clusters in the T cell compartment as a proportion of total T cells captured for each patient. The x-axis displays the rank-biserial correlation coefficient, while the y-axis displays the -log10 of the p-value from the Wilcoxon rank-sum test. Labels with lighter shades represent populations with significant differences ($p < 0.05$). Box plots displaying the distribution of patient proportions within a given cluster of $CD8^+$ **(b)** and $CD4^+$ **(c)** T cells. Circles represent individual patients. The significance of the difference in cell proportions was calculated using the Wilcoxon rank sum test and added to the panel if significant ($p < 0.05$).

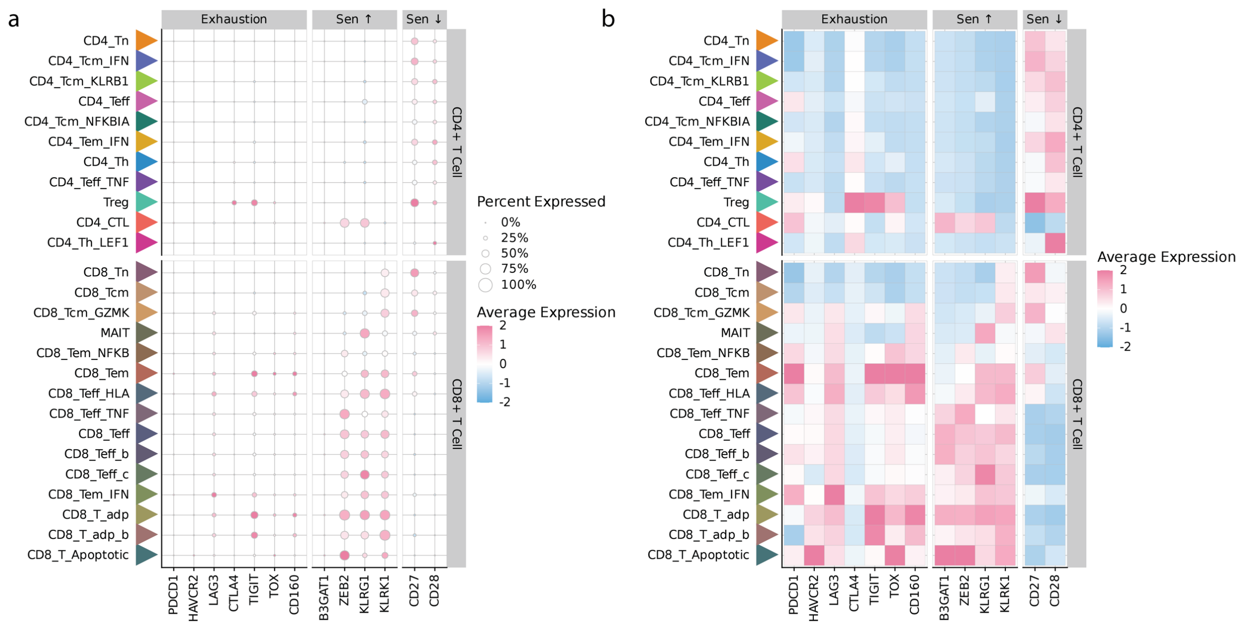


**Supplemental Figure 10: Exhaustion and Senescence marker expression across the T Cell compartment. (a)** Dot plot and **(b)** Tile plot displaying the scaled normalized average expression within each cluster of various exhaustion or senescence markers. Rows represent individual clusters, columns represent individual genes. Genes are grouped by whether they represent exhaustion, markers associated with senescence, or markers downregulated in senescence. In **(a)**, dot color represents average expression, with red representing higher average expression, and blue indicating lower expression. The size of the dot represents the percentage of cells expressing each gene in individual cell types. In **(b),** tile color represents the average gene expression with shades of red and blue representing high and low expression respectively.

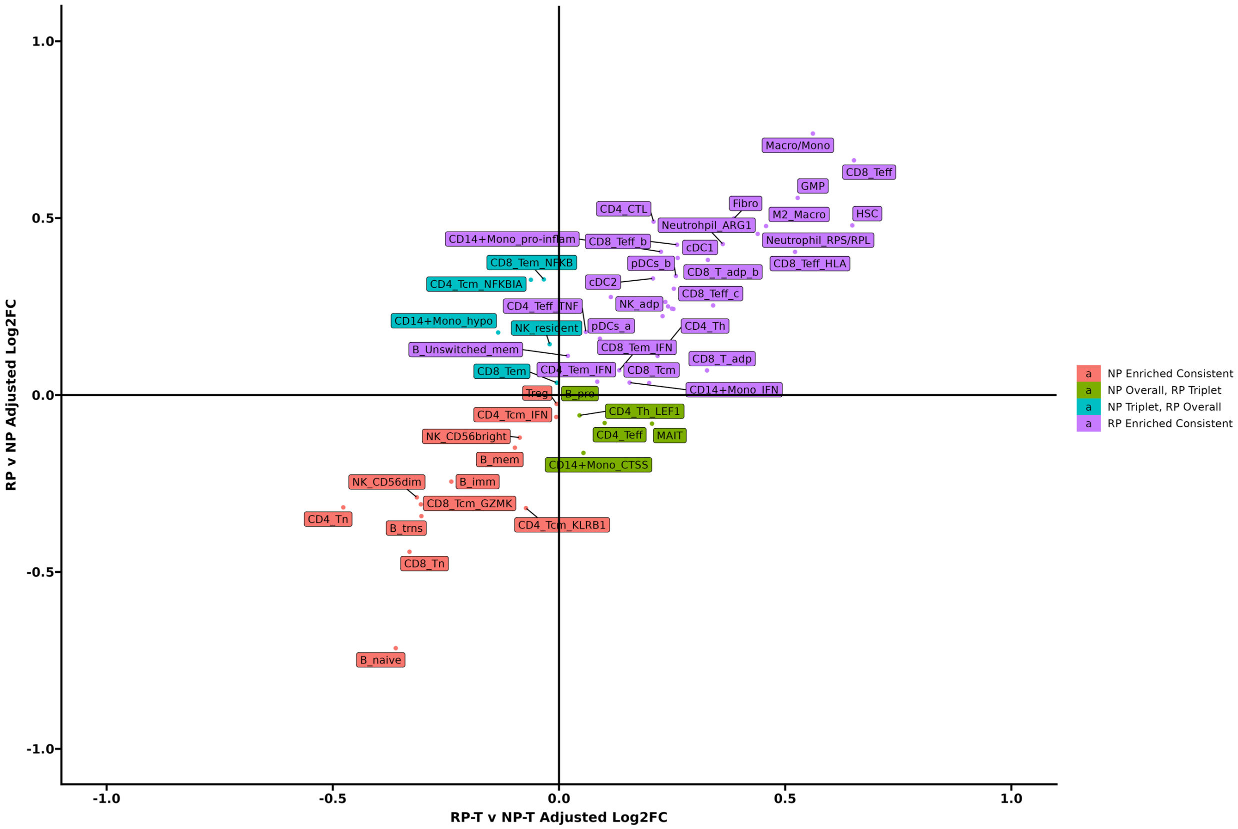


**Supplemental Figure 11:** Comparing the differential abundance profiles of Triplet therapy patients to the overall cohort. The X and Y axis displays the Dirichlet-estimated log2 fold change between RP (positive) and NP (negative) patients across triplet therapy patients (X axis) or all patients (Y axis) as a fraction of non-malignant cells. The majority of clusters show strong similarities between the triplet therapy sub-cohort and the overall population. Key RP populations such as GMPs, CD8_Teff, and CD8_Teff_HLA show consistent enrichment in RP in both groups (top right quadrant), while NP populations such as B_naive, CD8_Tn, and CD8_Tcm_GZMK show consistent enrichment in NP (bottom left quadrant).

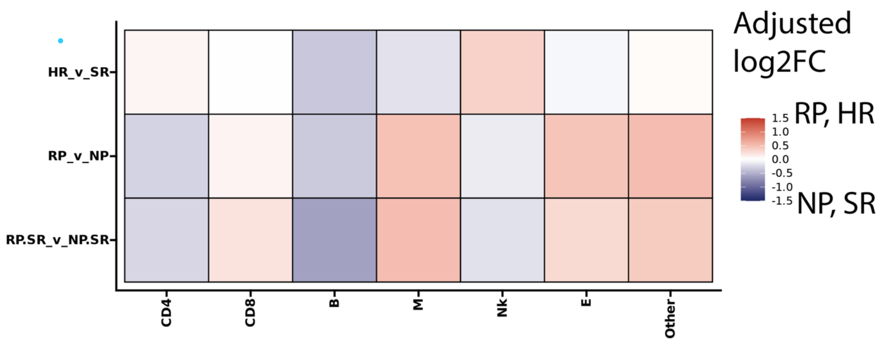


**Supplemental Figure 12:** Comparison of differential abundance results by cytogenetic risk (HR_v_SR), progression (RP_v_NP), and progression within standard-risk patients (RP.SR_v_NP.SR). Each box represents the estimated log2-fold change between different risk and progression based comparisons (HR v SR, RP v NP, RP.SR v NP.SR) after adjusting for batch using a Dirichlet model. Rows represent different comparisons, columns represent different major cell populations. Orange shades indicate RP, HR, or RP.SR upregulated major cell populations, while blue shades indicate SR, NP, or NP.SR upregulated populations.

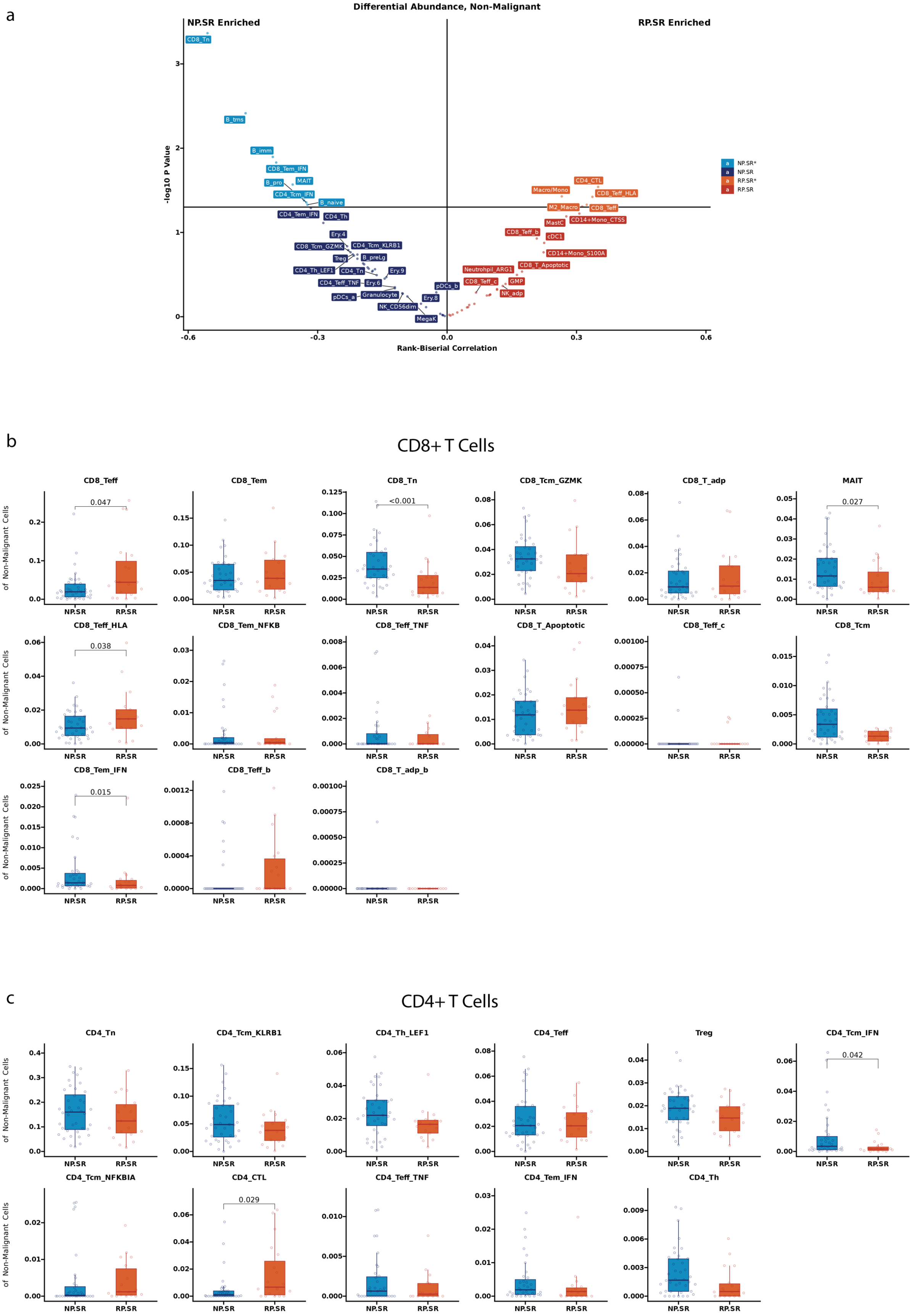
a
Differential Abundance, Non-Malignant
NP.SR Enriched
RP.SR Enriched
-log10 P Value
Rank-Biserial Correlation
NP.SR*
NP.SR
RP.SR*
RP.SR
CD8_Tn
B_trns
B_imm
CD8_Tem_IFN
B_pro
MAIT
CD4_Tcm_IFN
B_naive
CD4_Tem_IFN
CD4_Th
Ery.4
CD8_Tcm_GZMK
CD4_Tcm_KLRB1
Treg
B_preLg
CD4_Th_LEF1
CD4_Tn
Ery.9
CD4_Teff_TNF
Ery.6
pDCs_b
pDCs_a
Granulocyte
Ery.8
NK_CD56dim
MegaK
CD4_CTL
Macro/Mono
CD8_Teff_HLA
M2_Macro
CD8_Teff
CD14+Mono_CTSS
MastC
CD8_Teff_b
cDC1
CD14+Mono_S100A
CD8_T_Apoptotic
Neutrohpil_ARG1
CD8_Teff_c
GMP
NK_adp
b
CD8+ T Cells
Proportion of Non-Malignant Cells
CD8_Teff
0.047
CD8_Tem
CD8_Tn
<0.001
CD8_Tcm_GZMK
CD8_T_adp
MAIT
0.027
CD8_Teff_HLA
0.038
CD8_Tem_NFKB
CD8_Teff_TNF
CD8_T_Apoptotic
CD8_Teff_c
CD8_Tcm
CD8_Tem_IFN
0.015
CD8_Teff_b
CD8_T_adp_b
NP.SR
RP.SR
c
CD4+ T Cells
CD4_Tn
CD4_Tcm_KLRB1
CD4_Th_LEF1
CD4_Teff
Treg
CD4_Tcm_IFN
0.042
CD4_Tcm_NFKBIA
CD4_CTL
0.029
CD4_Teff_TNF
CD4_Tem_IFN
CD4_Th

d

### Myeloid Cells

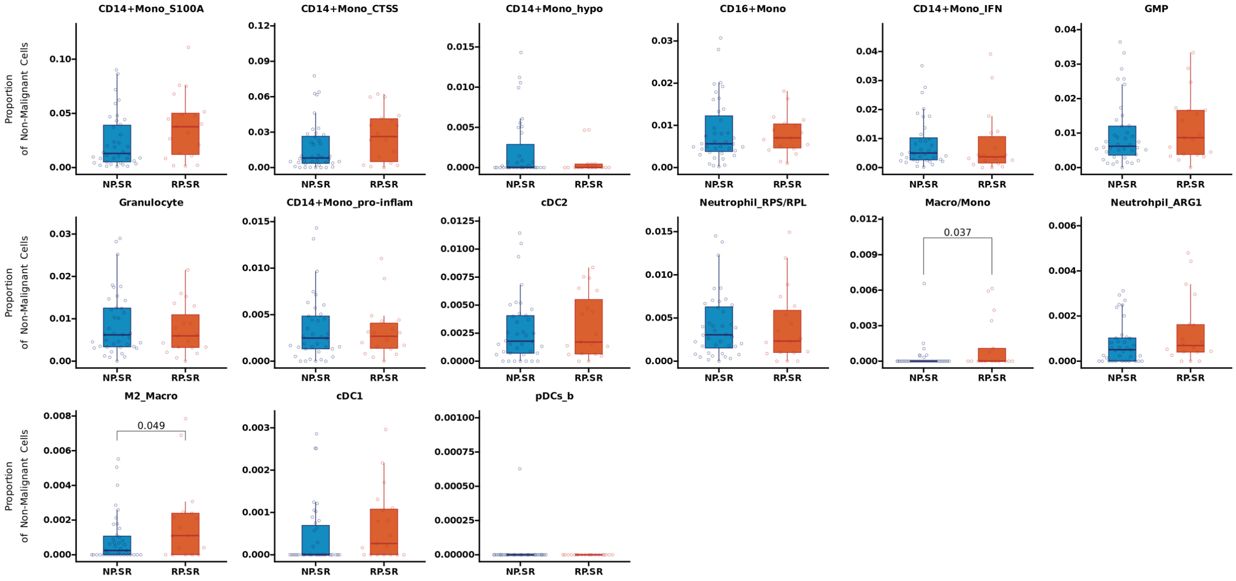


e

### B Cells

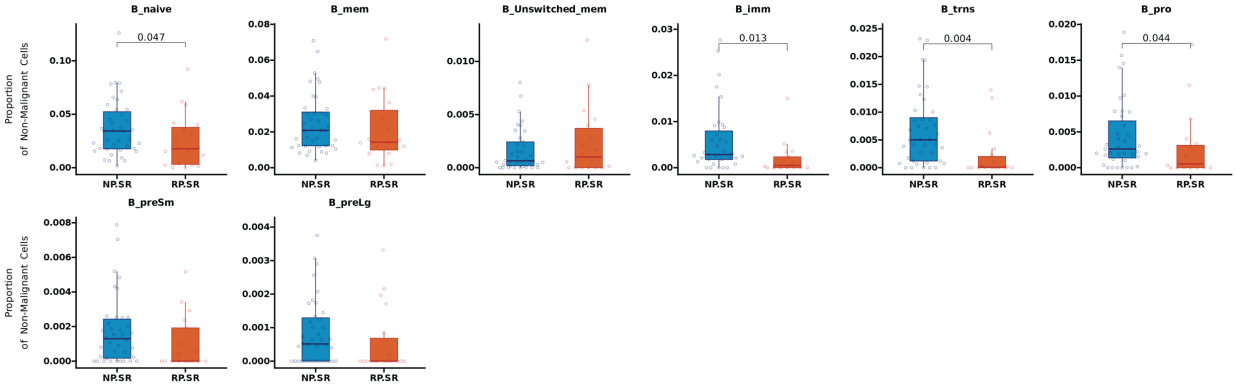


f

### Other Cells

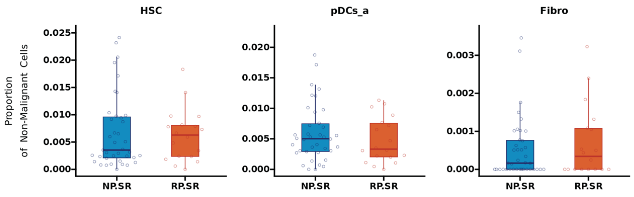

g

Natural Killer Cells

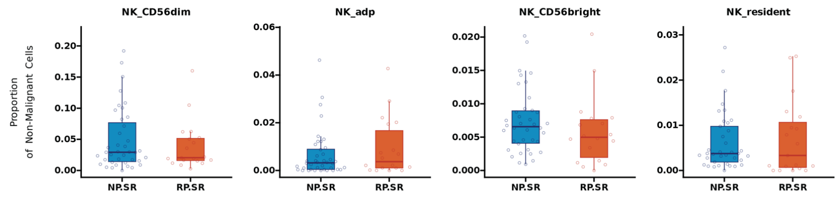


h

Erythroid Cells

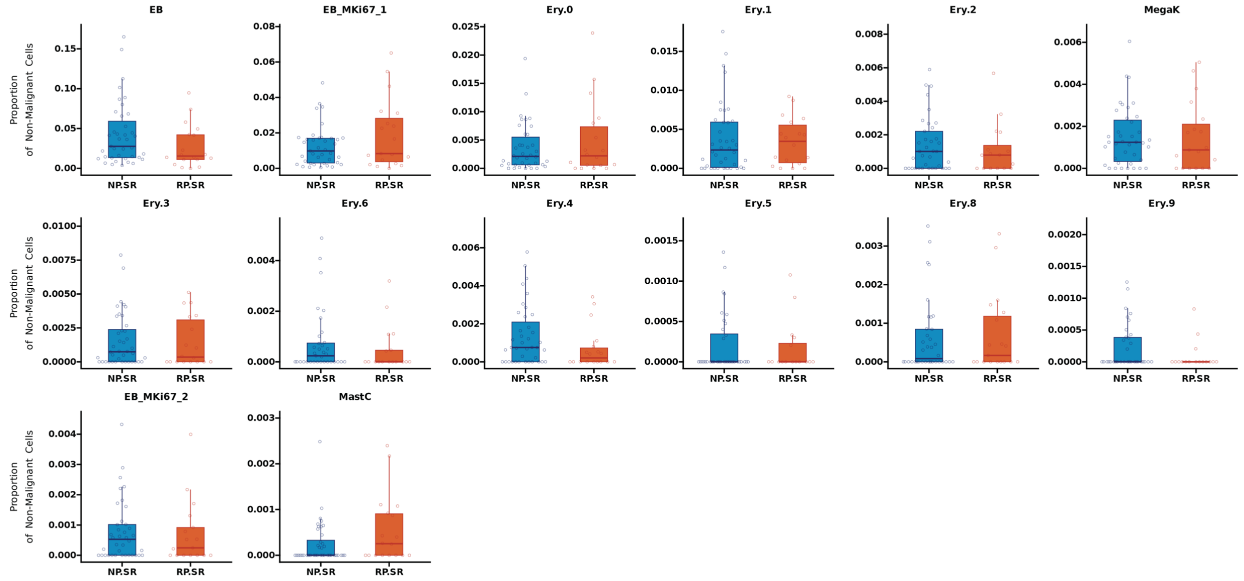


**Supplemental Figure 13: Differential Abundance Analysis for Non-Malignant Cells based on progression stratification of standard risk patients (Standard Risk Rapid Progressor (RP.SR) and Standard Risk Non-Progressors (NP.SR).** Differential abundance results comparing **NP.SR** and **ç** cellular proportions for each cluster. The cellular proportions for each patient were determined based on the total non-malignant cells. **a**. Scatter plot summarizing the differential abundance results. The x-axis displays the rank-biserial correlation coefficient, while the y-axis displays the -log10 of the p-value from the Wilcoxon rank-sum test. Labels with lighter shades represent clusters with significant changes in proportions between RP.SR and NP.SR non-progressing patients ($p < 0.05$). Box plots displaying the proportion of cells of a given cluster as a fraction of non-malignant cells, with clusters grouped by $CD8^{+}$ T cells **(b),** $CD4^{+}$ T cells (**c)**, Myeloid cells (**d)**, B Cells (**e)**, Other/Independent cell populations (**f)**, and Natural Killer cells (**g**), and Erythroid lineage cells (**h)**. The significance of the difference in cell proportions was calculated using the Wilcoxon rank sum test and added to the panel if significant ($p < 0.05$).

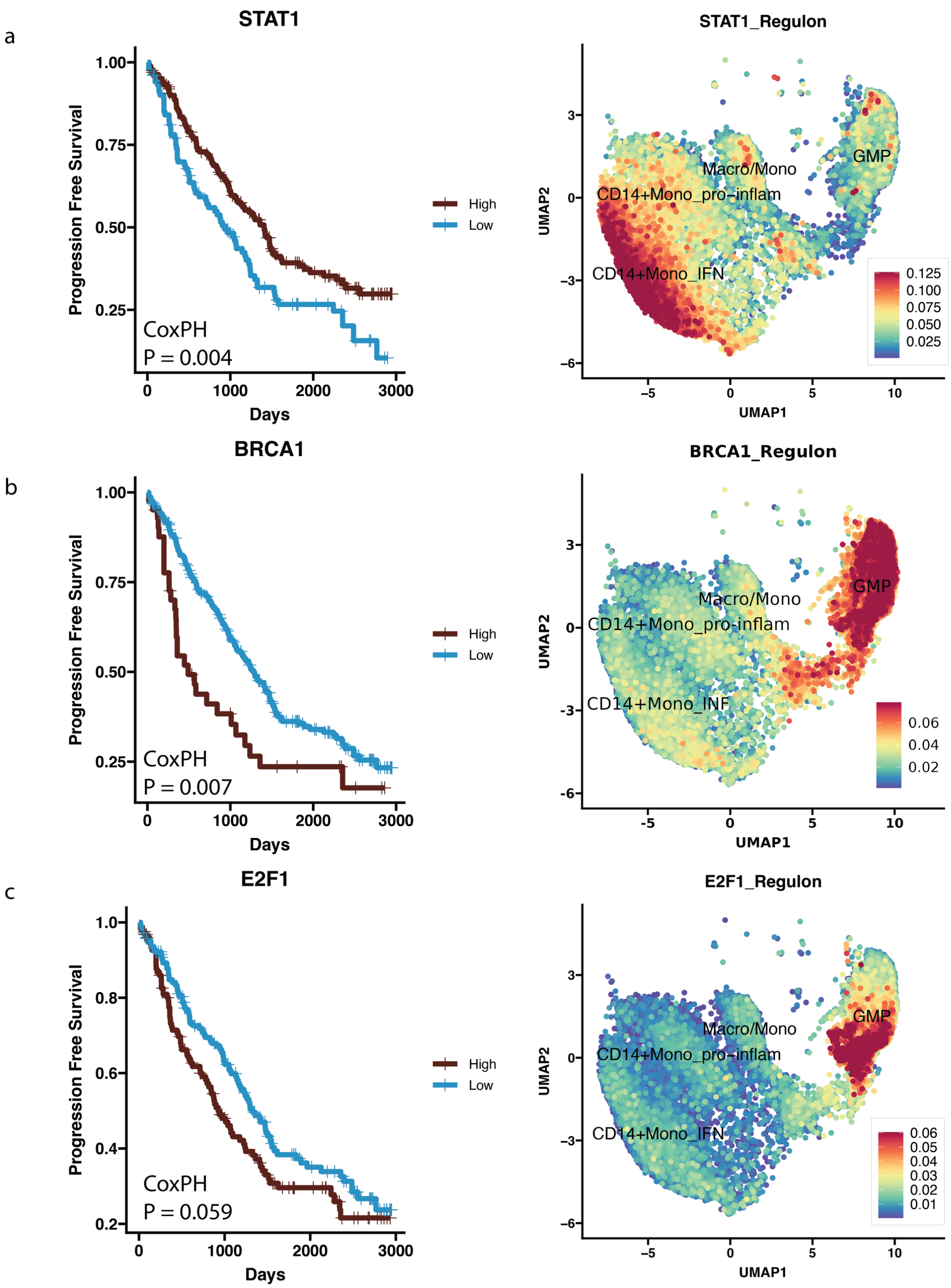
a
STAT1
Progression Free Survival
1.00
0.75
0.50
0.25
CoxPH
P = 0.004
0
1000
2000
3000
Days
High
Low
STAT1_Regulon
UMAP2
UMAP1
GMP
Macro/Mono
CD14+Mono_pro-inflam
CD14+Mono_IFN
0.125
0.100
0.075
0.050
0.025
b
BRCA1
Progression Free Survival
CoxPH
P = 0.007
Days
High
Low
BRCA1_Regulon
GMP
Macro/Mono
CD14+Mono_pro-inflam
CD14+Mono_INF
0.06
0.04
0.02
c
E2F1
Progression Free Survival
1.0
0.8
0.6
0.4
0.2
CoxPH
P = 0.059
Days
High
Low
E2F1_Regulon
GMP
Macro/Mono
CD14+Mono_pro-inflam
CD14+Mono_IFN
0.06
0.05
0.04
0.03
0.02
0.01

d

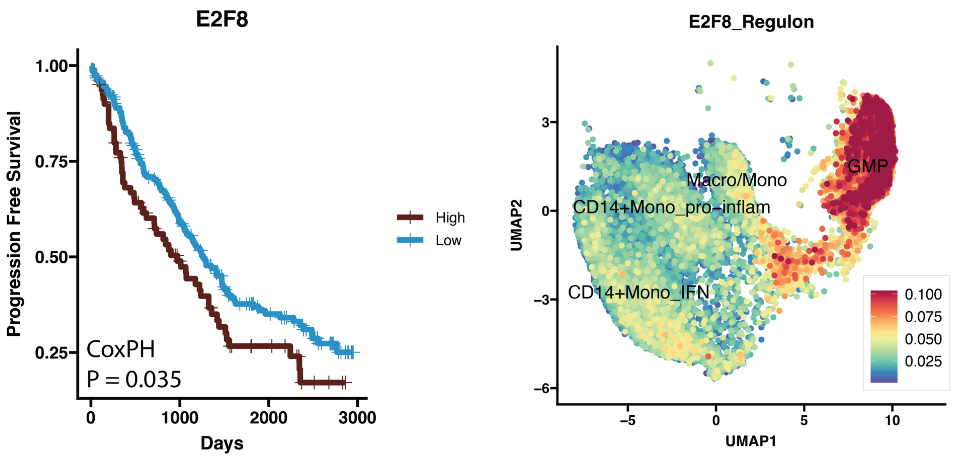

E2F8
1.00
0.75
0.50
0.25
Progression Free Survival
CoxPH
P = 0.035
0
1000
2000
3000
Days
High
Low
E2F8_Regulon
Macro/Mono
GMP
CD14+Mono_pro−inflam
CD14+Mono_IFN
0.100
0.075
0.050
0.025
3
0
−3
−6
UMAP2
−5
0
5
10
UMAP1


e

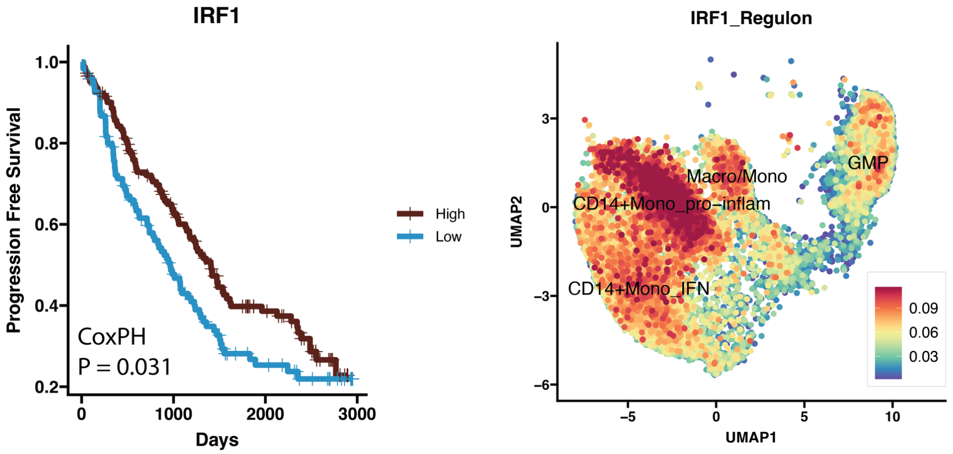

IRF1
1.0
0.8
0.6
0.4
0.2
Progression Free Survival
CoxPH
P = 0.031
0
1000
2000
3000
Days
High
Low
IRF1_Regulon
Macro/Mono
GMP
CD14+Mono_pro−inflam
CD14+Mono_IFN
0.09
0.06
0.03
3
0
−3
−6
UMAP2
−5
0
5
10
UMAP1


f

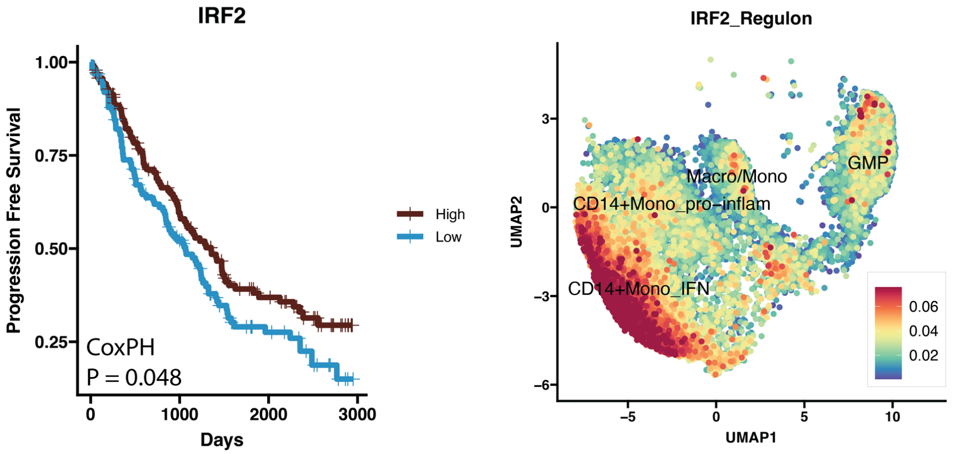

IRF2
1.00
0.75
0.50
0.25
Progression Free Survival
CoxPH
P = 0.048
0
1000
2000
3000
Days
High
Low
IRF2_Regulon
Macro/Mono
GMP
CD14+Mono_pro−inflam
CD14+Mono_IFN
0.06
0.04
0.02
3
0
−3
−6
UMAP2
−5
0
5
10
UMAP1

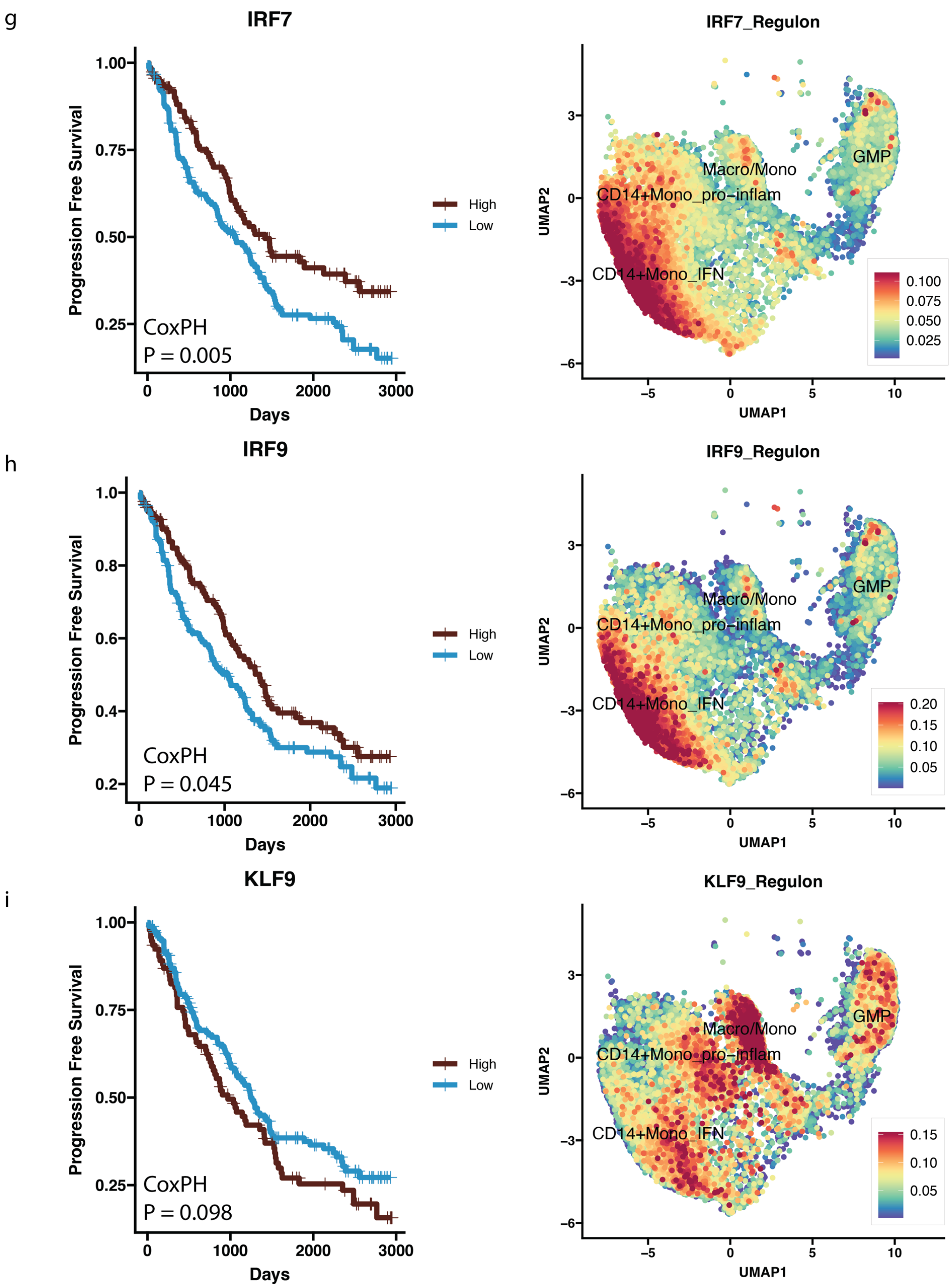


**Supplemental Figure 14: Survival analysis and gene regulator scores for the top gene regulators of select myeloid populations.** Gene regulatory networks were created using pySCENIC on select myeloid clusters. Kaplan-Meier curves (left) and feature plots (right) for the select myeloid populations are displayed for STAT1 **(a),** BRCA1 **(b)**, E2F1 **(c),** E2F8 **(d),** IRF1 **(e)**, IRF2 **(f)**, IRF7 **(g)**, IRF9 **(h)**, KLF9 **(i)**. (left) Kaplan-Meier curves based on average AUC value across the selected myeloid populations. The cut-point approach was implemented to identify split patients into high and low groups for survival analysis. P value indicates the significance of a

CoxPH model fitted to the regulon AUC score across select myeloid clusters. (right) A feature plot displaying the per-cell AUC score across the select myeloid clusters.

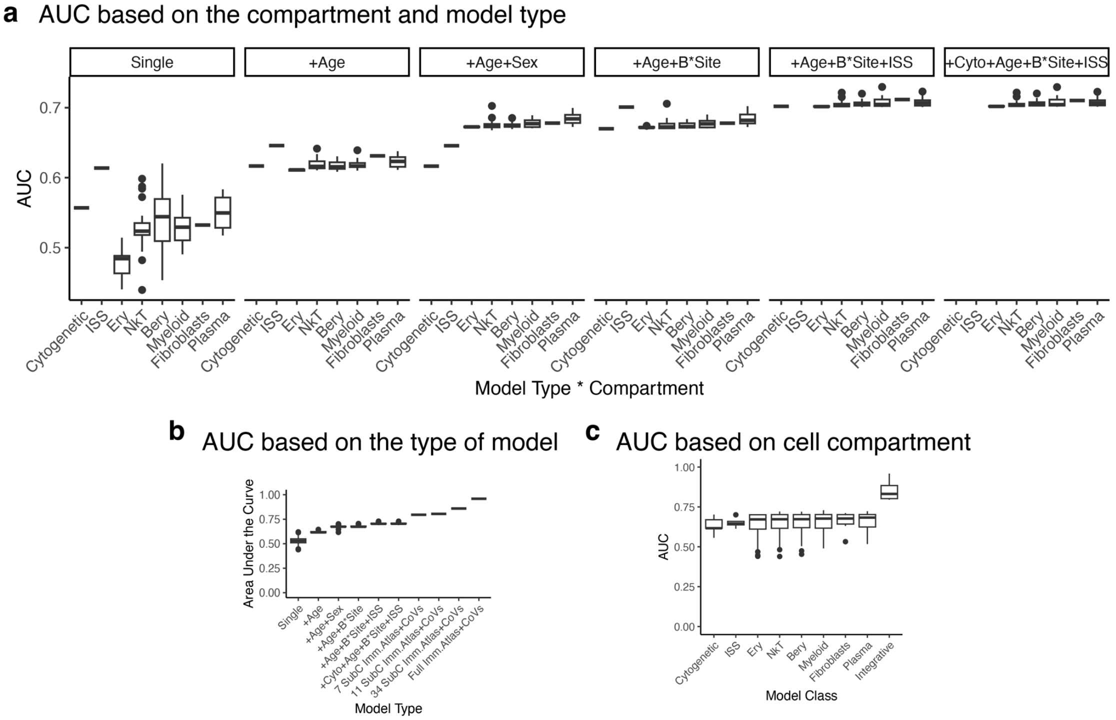


**d**

### LRM Bootstrap Validation Diagnostic Tables

#### ISS+Age+Sex+Batch+Site

| | index.orig | training | test | optimism | index.corrected | n | auc |
|---|---|---|---|---|---|---|---|
| *Dxy* | 0.4 | 0.47 | 0.36 | 0.11 | 0.29 | 1000 | 0.7 |
| *R2* | 0.16 | 0.21 | 0.12 | 0.09 | 0.07 | 1000 | 0.7 |
| *Intercept* | 0 | 0 | 0.12 | −0.12 | 0.12 | 1000 | 0.7 |
| *Slope* | 1 | 1 | 0.72 | 0.28 | 0.72 | 1000 | 0.7 |
| *Emax* | 0 | 0 | 0.09 | 0.09 | 0.09 | 1000 | 0.7 |
| *D* | 0.12 | 0.17 | 0.09 | 0.07 | 0.05 | 1000 | 0.7 |
| *U* | −0.01 | −0.01 | 0.02 | −0.02 | 0.02 | 1000 | 0.7 |
| *Q* | 0.13 | 0.17 | 0.08 | 0.1 | 0.03 | 1000 | 0.7 |
| *B* | 0.21 | 0.2 | 0.22 | −0.02 | 0.23 | 1000 | 0.7 |
| *g* | 0.9 | 1.09 | 0.77 | 0.32 | 0.58 | 1000 | 0.7 |
| *gp* | 0.19 | 0.22 | 0.17 | 0.05 | 0.14 | 1000 | 0.7 |

#### CYTO+ISS+Age+Sex+Batch+Site

| | index.orig | training | test | optimism | index.corrected | n | auc |
|---|---|---|---|---|---|---|---|
| *Dxy* | 0.4 | 0.47 | 0.35 | 0.12 | 0.28 | 1000 | 0.7 |
| *R2* | 0.16 | 0.21 | 0.12 | 0.1 | 0.06 | 1000 | 0.7 |
| *Intercept* | 0 | 0 | 0.13 | −0.13 | 0.13 | 1000 | 0.7 |
| *Slope* | 1 | 1 | 0.69 | 0.31 | 0.69 | 1000 | 0.7 |
| *Emax* | 0 | 0 | 0.1 | 0.1 | 0.1 | 1000 | 0.7 |
| *D* | 0.12 | 0.17 | 0.09 | 0.08 | 0.04 | 1000 | 0.7 |
| *U* | −0.01 | −0.01 | 0.02 | −0.03 | 0.02 | 1000 | 0.7 |
| *Q* | 0.13 | 0.18 | 0.07 | 0.11 | 0.02 | 1000 | 0.7 |
| *B* | 0.21 | 0.2 | 0.22 | −0.02 | 0.23 | 1000 | 0.7 |
| *g* | 0.9 | 1.1 | 0.75 | 0.35 | 0.55 | 1000 | 0.7 |
| *gp* | 0.19 | 0.22 | 0.16 | 0.06 | 0.14 | 1000 | 0.7 |

#### IA1+CYTO+ISS+Age+Sex+Batch+Site

| | index.orig | training | test | optimism | index.corrected | n | auc |
|---|---|---|---|---|---|---|---|
| *Dxy* | 0.46 | 0.52 | 0.4 | 0.12 | 0.34 | 1000 | 0.73 |
| *R2* | 0.2 | 0.26 | 0.16 | 0.1 | 0.1 | 1000 | 0.73 |
| *Intercept* | 0 | 0 | 0.12 | −0.12 | 0.12 | 1000 | 0.73 |
| *Slope* | 1 | 1 | 0.71 | 0.29 | 0.71 | 1000 | 0.73 |
| *Emax* | 0 | 0 | 0.1 | 0.1 | 0.1 | 1000 | 0.73 |
| *D* | 0.16 | 0.21 | 0.12 | 0.09 | 0.07 | 1000 | 0.73 |
| *U* | −0.01 | −0.01 | 0.02 | −0.03 | 0.02 | 1000 | 0.73 |
| *Q* | 0.16 | 0.21 | 0.1 | 0.11 | 0.05 | 1000 | 0.73 |
| *B* | 0.2 | 0.19 | 0.21 | −0.02 | 0.23 | 1000 | 0.73 |
| *g* | 1.16 | 1.41 | 0.98 | 0.44 | 0.73 | 1000 | 0.73 |
| *gp* | 0.21 | 0.24 | 0.19 | 0.05 | 0.16 | 1000 | 0.73 |

#### IA7+CYTO+ISS+Age+Sex+Batch+Site

| | index.orig | training | test | optimism | index.corrected | n | auc |
|---|---|---|---|---|---|---|---|
| *Dxy* | 0.59 | 0.66 | 0.53 | 0.13 | 0.46 | 1000 | 0.8 |
| *R2* | 0.33 | 0.4 | 0.26 | 0.14 | 0.19 | 1000 | 0.8 |
| *Intercept* | 0 | 0 | 0.11 | −0.11 | 0.11 | 1000 | 0.8 |
| *Slope* | 1 | 1 | 0.68 | 0.32 | 0.68 | 1000 | 0.8 |
| *Emax* | 0 | 0 | 0.11 | 0.11 | 0.11 | 1000 | 0.8 |
| *D* | 0.27 | 0.35 | 0.21 | 0.13 | 0.14 | 1000 | 0.8 |
| *U* | −0.01 | −0.01 | 0.04 | −0.04 | 0.04 | 1000 | 0.8 |
| *Q* | 0.28 | 0.36 | 0.18 | 0.18 | 0.1 | 1000 | 0.8 |
| *B* | 0.18 | 0.16 | 0.2 | −0.03 | 0.21 | 1000 | 0.8 |
| *g* | 1.76 | 2.14 | 1.44 | 0.71 | 1.05 | 1000 | 0.8 |
| *gp* | 0.28 | 0.31 | 0.25 | 0.06 | 0.22 | 1000 | 0.8 |

#### IA11+CYTO+ISS+Age+Sex+Batch+Site

| | index.orig | training | test | optimism | index.corrected | n | auc |
|---|---|---|---|---|---|---|---|
| *Dxy* | 0.61 | 0.69 | 0.54 | 0.16 | 0.45 | 1000 | 0.8 |
| *R2* | 0.35 | 0.44 | 0.27 | 0.17 | 0.18 | 1000 | 0.8 |
| *Intercept* | 0 | 0 | 0.13 | −0.13 | 0.13 | 1000 | 0.8 |
| *Slope* | 1 | 1 | 0.62 | 0.38 | 0.62 | 1000 | 0.8 |
| *Emax* | 0 | 0 | 0.13 | 0.13 | 0.13 | 1000 | 0.8 |
| *D* | 0.29 | 0.39 | 0.22 | 0.17 | 0.12 | 1000 | 0.8 |
| *U* | −0.01 | −0.01 | 0.06 | −0.07 | 0.06 | 1000 | 0.8 |
| *Q* | 0.3 | 0.4 | 0.16 | 0.24 | 0.06 | 1000 | 0.8 |
| *B* | 0.17 | 0.15 | 0.2 | −0.04 | 0.22 | 1000 | 0.8 |
| *g* | 1.86 | 2.38 | 1.45 | 0.93 | 0.94 | 1000 | 0.8 |
| *gp* | 0.29 | 0.32 | 0.25 | 0.07 | 0.21 | 1000 | 0.8 |

#### IA34+CYTO+ISS+Age+Sex+Batch+Site

| | index.orig | training | test | optimism | index.corrected | n | auc |
|---|---|---|---|---|---|---|---|
| *Dxy* | 0.72 | 0.86 | 0.57 | 0.28 | 0.44 | 996 | 0.86 |
| *R2* | 0.47 | 0.66 | 0.24 | 0.42 | 0.05 | 996 | 0.86 |
| *Intercept* | 0 | 0 | 0.24 | −0.24 | 0.24 | 996 | 0.86 |
| *Slope* | 1 | 1 | 0.26 | 0.74 | 0.26 | 996 | 0.86 |
| *Emax* | 0 | 0 | 0.33 | 0.33 | 0.33 | 996 | 0.86 |
| *D* | 0.42 | 0.66 | 0.19 | 0.47 | −0.05 | 996 | 0.86 |
| *U* | −0.01 | −0.01 | 0.72 | −0.73 | 0.72 | 996 | 0.86 |
| *Q* | 0.43 | 0.67 | −0.53 | 1.2 | −0.77 | 996 | 0.86 |
| *B* | 0.15 | 0.1 | 0.2 | −0.1 | 0.25 | 996 | 0.86 |
| *g* | 2.8 | 5.69 | 1.41 | 4.29 | −1.48 | 996 | 0.86 |
| *gp* | 0.34 | 0.4 | 0.22 | 0.18 | 0.16 | 996 | 0.86 |

**Supplemental Figure 15: Performance comparison of individual model components and validation metrics.** (**a)** Boxplots showing the AUC values based on the immune atlas cell compartment and model using different combinations of variables. (**b)** Boxplots showing the AUC based only on different variable combinations regardless of the immune atlas cell compartment. **(c)** Boxplots showing the AUCs based on cell compartments with covariates but integrative cytogenetics with immune atlas and covariates separately. (**d)** Bootstrap validation and diagnostics tables for key prediction models. The tables contain Dxy (Somer's index), R2 (Nagelkerke index). An adjusted version of the Cox & Snell R-square that adjusts the scale of the statistic to cover the full range from 0 to 1. This pseudo-statistic measures the improvement in the fitted model compared to the null model. D is a classic discrimination index, showing the difference in quality between the best constant predictor (one that on average predicts the overall prevalence of an event) and the best-calibrated predictor. U is the unreliability index — a unitless index of how far the logit calibration curve intercept and slope are from (0, 1). Q is the overall quality. Q = D - U. It represents the logarithmic accuracy score. Emax is the maximum difference between raw predicted probabilities, and the recalibrated probabilities. B or Brier score shows the mean square error between predictions and observations. G or Gini's mean difference index, calculated on the log-odds ratio scale. A measure of the model's predictive discrimination based only on the fitted values. GP is the Gini index based on the probability scale.
